# nQuack: An R package for predicting ploidal level from sequence data using site-based heterozygosity

**DOI:** 10.1101/2024.02.12.579894

**Authors:** Michelle L. Gaynor, Jacob B. Landis, Timothy K. O’Connor, Robert G. Laport, Jeff J. Doyle, Douglas E. Soltis, José Miguel Ponciano, Pamela S. Soltis

## Abstract

**Premise:** Traditional methods of ploidal level estimation are tedious; leveraging sequence data for cytotype estimation is an ideal alternative. Multiple statistical approaches to leverage DNA sequence data for ploidy prediction based on site-based heterozygosity have been developed. However, these approaches may require high-coverage sequence data, use improper probability distributions, or have additional statistical shortcomings that limit inference abilities. We introduce nQuack, an open-source R package, that addresses the main shortcomings of current methods.

**Methods and Results:** nQuack performs model selection for improved ploidy predictions. Here, we implement expected maximization algorithms with normal, beta, and beta-binomial distributions. Using extensive computer simulations that account for variability in sequencing depth, as well as real data sets, we demonstrate the utility and limitations of nQuack.

**Conclusion:** Inferring ploidal level based on site-based heterozygosity alone is discouraged due to the low accuracy of pattern-based inference.

## INTRODUCTION

Whole-genome duplication (WGD), or polyploidy, is ubiquitous across the plant tree of life, with all extant angiosperms having evidence of at least one ancient WGD (Jiao et al., 2011; Soltis et al., 2015; Landis et al., 2018; One Thousand Plant Transcriptomes Initiative, 2019). Identifying ploidal diversity is a crucial first step to understanding the impact of WGD on patterns of biodiversity. Direct estimation is achieved through chromosome counting at either mitosis or meiosis. However, indirect estimation (e.g., flow cytometry, stomatal cell measurements, pollen size) can be used for broad surveys of select taxa when complemented with known chromosome numbers and/or ploidal levels (Masterson, 1994; Beaulieu et al., 2008; Sanders, 2021; Sliwinska et al., 2021). The application of flow cytometry to determine ploidal level in naturally occurring populations (Galbraith et al., 1983; Keeler et al., 1987) has been fundamental to understanding evolution and ecology of mixed-ploidy populations. Despite the utility of laboratory-based approaches and the extension of flow cytometry to dried samples (Galbraith et al., 1983; Keeler et al., 1987), the process remains specialized and may involve the use of laboratory equipment that is difficult to access. Therefore, using DNA sequence data for ploidal-level prediction affords a great opportunity to streamline estimation while revolutionizing our understanding of chromosome evolution.

To date, multiple statistical approaches to leverage DNA sequence data for the prediction of ploidy have been developed based on (1) k-mer and (2) site-based heterozygosity. Both of these general methods for ploidal-level prediction require statistical tests to assign ploidal level to a sample; the statistical approach varies among available software.

K-mer-based ploidal-level prediction relies on a k-mer profile, which classifies the frequency of each distinct k-mer found across the dataset. K-mers are strings of length k, often 21 bases (Vurture et al., 2017), that are composed of a specific sequence of nucleotides. Popular methods for k-mer-based ploidal-level prediction are tetmer (Becher et al., 2022) and smudgeplot, which plots minor allele frequency by total coverage to predict copy number variants (Ranallo-Benavidez et al., 2020). These methods have been recently expanded to single-cell ATAC-seq data (Takeuchi and Kato, 2023). However, a limitation of these methods is that at least 15-25x sequence coverage per homolog is required.

Site-based heterozygosity relies on biallelic single nucleotide polymorphisms (SNPs) within an individual and the expected number of copies of each base at that SNP. For example, in a diploid individual, at a biallelic site with alleles A and B, about 50% of all nucleotides sequenced are expected to represent allele A. Comparatively, in a triploid, at a site with alleles A and B, 33% of the nucleotides are expected to be allele A and 67% allele B, or vice versa (Figure 1), are expected. The most commonly used site-based heterozygosity software is nQuire (Weiß et al., 2018), but additional software exists for de novo sequences (Sun et al., 2023). As for k-mer-based estimation, sequence coverage per site of at least 20-25x is recommended for the use of nQuire (Weiß et al., 2018).

**Figure 1.**
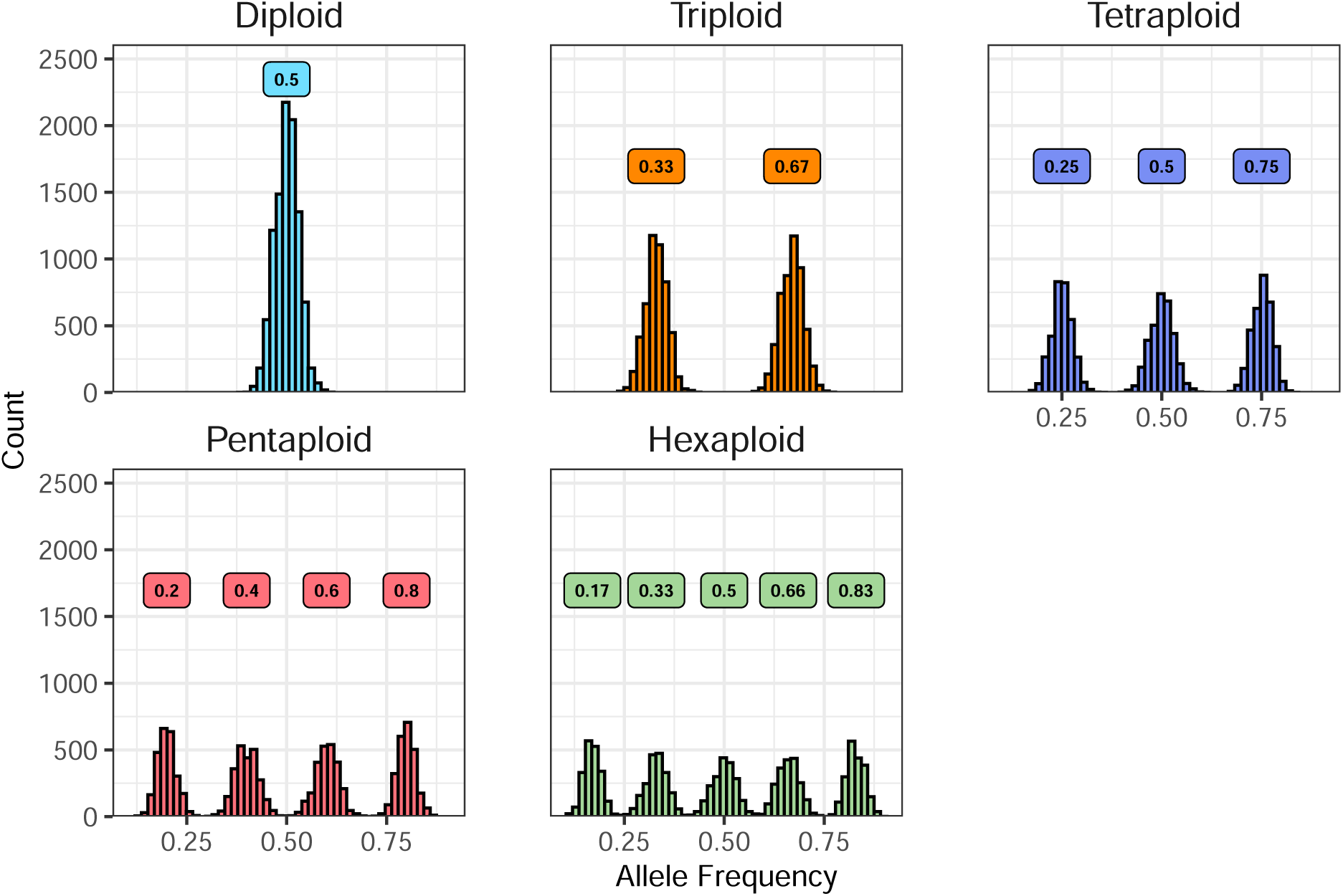
Expected allele frequencies at biallelic sites for diploid, triploid, tetraploid, pentaploid, and hexaploid.

In addition, the performance and limitations of nQuire are poorly understood in terms of accuracy. Combining nQuire’s model inference with additional data, such as genome size estimates, and with goodness-of-fit tests has been suggested (Viruel et al., 2019). Notably, nQuire’s accuracy and limitations were assessed using only genome resequencing data for only five samples, representing two taxonomic groups (Weiß et al., 2018). Numerous studies have since identified inconsistencies between nQuire’s estimates and indirect or direct ploidal estimates (Jantzen et al., 2022; Folk et al., 2023; Landis and Doyle, 2023).

Moreover, to concerns regarding accuracy, guidelines for data preparation are limited, since it is unknown how nQuire predictions are influenced by the number of sites, sequencing coverage, and amount of variance or noise in a dataset. In real data sets, this noise can be introduced through sequence error or general mapping error, as well as through the inclusion of non-single-copy loci.

Here we introduce nQuack, an R package that (1) provides expanded tools and implementations to improve site-based heterozygosity inferences of ploidal level, and (2) rigorously evaluates the accuracy of this method and an existing method, nQuire. Specifically, nQuack implements expected maximization algorithms with normal, beta, and beta-binomial distributions to identify the ploidal level (ranging from diploid to hexaploid) of samples based on DNA sequence data, building upon the framework proposed by nQuire. We designed three new implementations of the expected maximization algorithm which allow additional distributions to be tested. Although we implement the normal distribution, as used in nQuire, this distribution may be ill-suited for allele frequencies as they range from 0 to 1 and the normal distribution ranges from negative infinity to infinity. Our second implementation uses a beta distribution to match the constrained range of allele frequencies. Because sequence data provide allele counts, frequencies represent transformed data, which may lack original data attributes and misrepresent sampling variances and one or more sources of heterogeneity. Therefore, our final implementation includes the beta-binomial distribution, which allows raw allele counts to be leveraged.

We rigorously tested our new implementations to identify limitations to these new methods and provide guidance for users. We examine nQuire’s five samples in addition to 477 samples representing three additional taxonomic groups and three additional sequence data types (genotype-by-sequencing, target enrichment, and RadCap). To provide recommendations regarding coverage and the number of sites needed for each implementation and model type, we also test our model on 355 simulated samples, representing two simulation approaches that vary in the amount of variance introduced.

## METHODS AND RESULTS

### Likelihood calculations and model selection

The basis of our models is the expected allele frequency at variable biallelic sites for each ploidal level including diploid (0.5), triploid (0.33, 0.67), tetraploid (0.25, 0.5, 0.75), pentaploid (0.2, 0.4, 0.6, 0.8), and hexaploid (0.17, 0.33, 0.5, 0.67, 0.83), as described above (Figure 1; Appendix S1). To use the expected allele frequencies to determine the most likely ploidal level given a set of allele frequencies or allele counts, we developed three implementations of expected maximization algorithms with the normal, beta, and beta-binomial distributions, each with and without a uniform distribution to capture uniform noise components. The normal distribution implemented here differs from that of nQuire in our augmented-likelihood calculation (Appendix S1; see Supporting Information with this article), however, all model comparisons were investigated with both the nQuire-style implementation and our implementation of the normal distribution (Appendix S1). We found our implementation to have lower confidence in incorrect models compared to nQuire’s implementation, and therefore we focus only on our implementation of the normal distribution here.

The details of our implementations, though summarized here, can be found in Appendix S1. Given the expected frequencies, the likelihood for each ploidal level based on a set of observed allele frequencies (or allele counts) is defined as the sum of the product of the mixture proportion (alpha) and the relative likelihood of the observations, or probability density function, based on the expected frequency (mean) and variance of that mixture and the given distribution (Figure 2). To maximize the likelihood for a set of mixtures, values of alpha, variance, and mean can be modified through the expected maximization algorithm and optimized with the Nelder-Mead simplex optimization algorithm (Nelder and Mead, 1965). Furthermore, to allow model selection via information criteria, where divergence among models can be estimated by calculating the log-likelihood ratio, we allow ‘free’ and ‘fixed’ models, where all ‘fixed’ models are nested in a ‘free’ model. In our free model, all parameter values (alpha, variance, and mean) are estimated for a mixture of all potential ploidal levels. Although we have an expected value for the mean of each mixture, the expected values of alpha, as well as the variance, are not well-defined. We know that the proportions of each type of heterozygote may differ for an allopolyploid compared to an autopolyploid (see Lloyd and Bomblies, 2016), so we were interested in exploring models where alpha is free. Therefore, we tested three ‘fixed’ models: (1) where only alpha is free, (2) where only variance is free, and (3) where both alpha and variance are free. Therefore, for each implementation, we provide 32 model types, including three fixed models, at each of the five ploidal levels examined here and one ‘free’ model, all of which can be examined with and without a uniform distribution.

**Figure 2.**
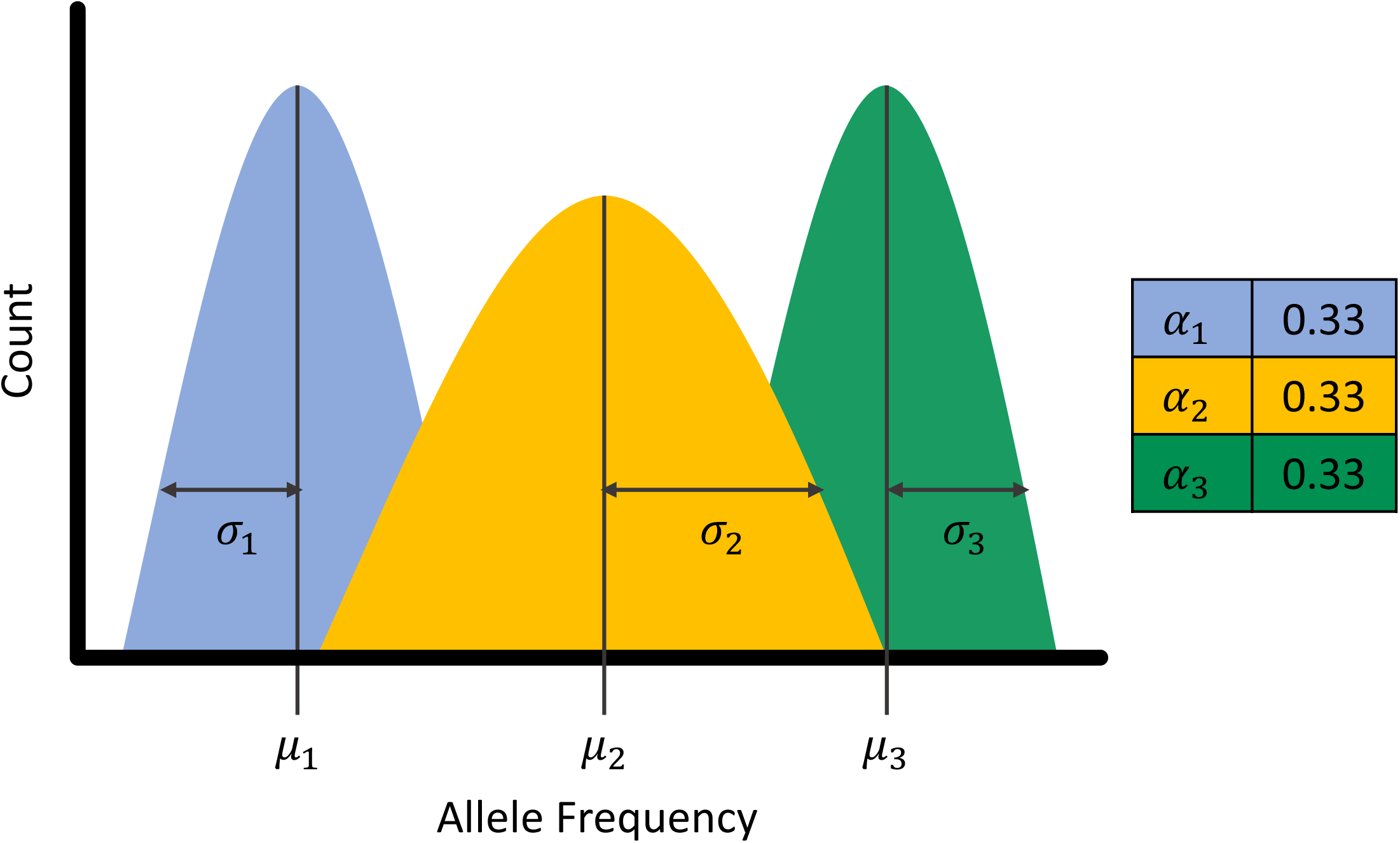
The basic components of a mixture model include mean (μ), variance (**σ**), and proportion (or alpha, α). The expected distributions for an autotetraploid, as defined by Lloyd and Bomblies 2016, can be seen here.

To evaluate each model, we examined the log-likelihood ratio and the Bayesian Information Criteria, or BIC score. The BIC score is the log-likelihood of a model penalized by both sample size and the number of parameters included, which leads to less error in model selection (Taper et al., 2021). We examined both the log-likelihood ratio and BIC score for all models and determined that BIC identified the correct ploidal level of more samples than the log-likelihood ratio; thus, we focused on BIC scores in all model comparisons. In theory, the BIC difference between the best and second-best model can be leveraged as an information criterion to assess confidence in model selection (Jerde et al., 2019; Taper et al., 2021).

### Model evaluation

To evaluate our models and determine guidelines for implementing these models, we examined 513,792 models based on both simulated and real samples. Simulated data representing all five ploidal levels varied in sequence coverage and number of sites, as well as the amount of random noise. Real samples include 482 samples of known ploidy (Table 1), inferred via indirect and direct estimates, and represent five taxonomic groups and four types of sequence data.

**Table 1.**
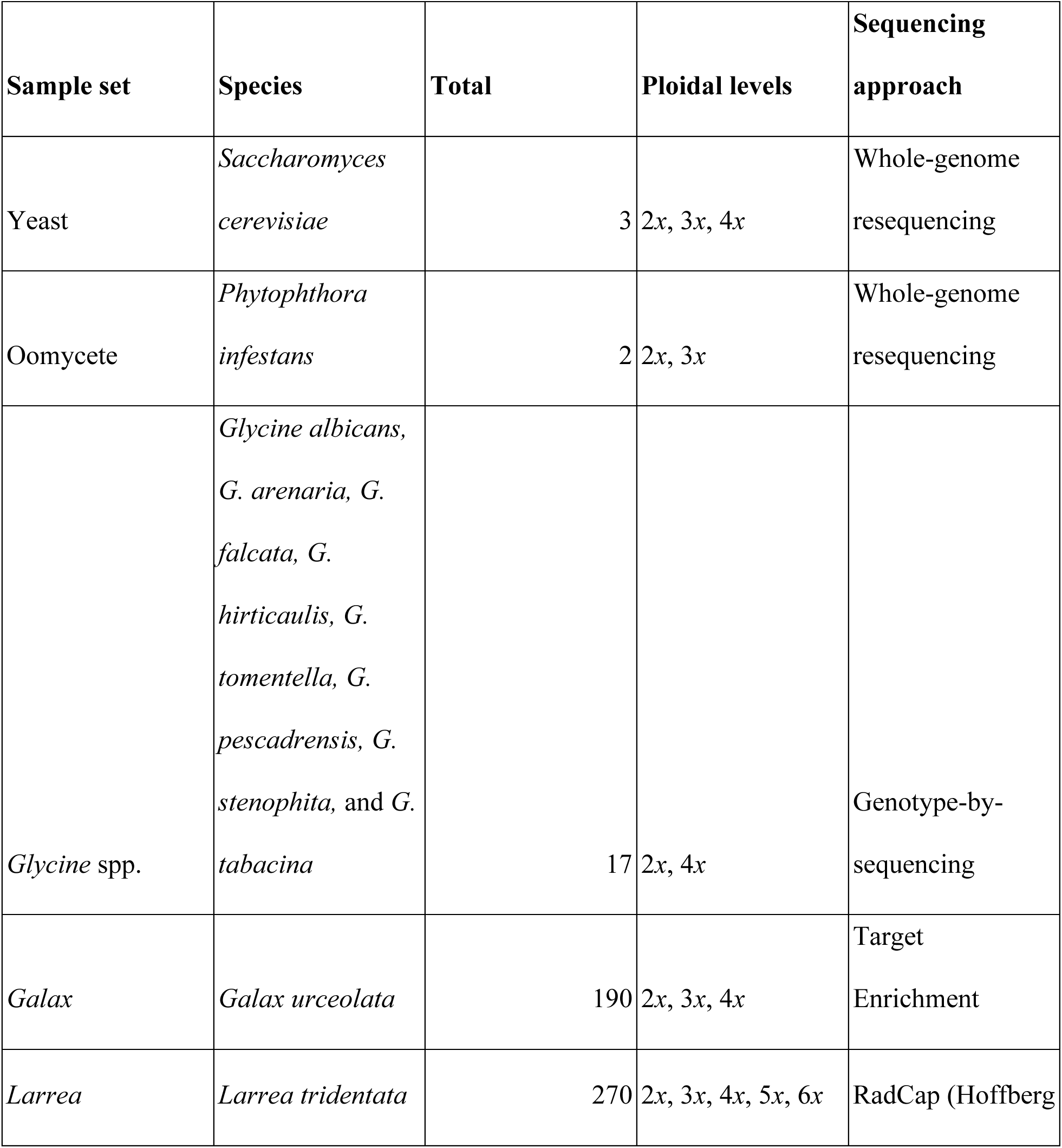
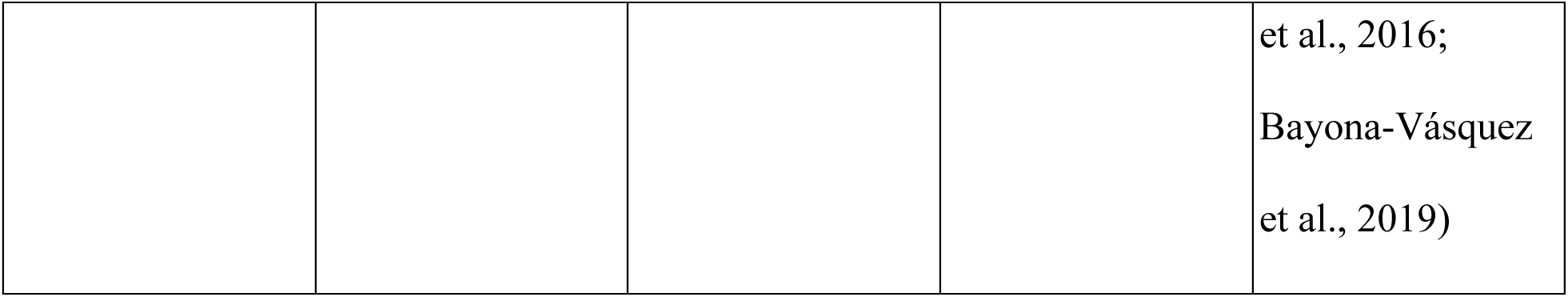
An overview of all included sample sets including the species, total number of samples, ploidal levels included, and sequencing approach. Additional information, including available accessions, can be found in Appendix S3 and Appendix S4.

### Simulated data

We simulated samples based on two approaches that represent two sampling scenarios: a “simplistic” one and a “realistic” scenario where the sampling is done at various levels of DNA sequence coverage (3-120x). Simplistic simulated samples are simple, with little to no variance introduced during the simulation process. The simplistic approach simulates heterozygous biallelic sites based on a binomial distribution where coverage among sites is equal and all expected frequencies have an equal probability of being sampled. For each ploidal level, we simulated 11 samples that differed in coverage per site (5, 10, 20, 30, 40, 50, 60, 70, 80, 90, or 100). For the 55 simulated samples, models were evaluated at six different numbers of sites, or the total number of SNPs (1250, 2500, 5000, 10000, 20000, 30000).

For our realistic simulations, we simulated samples where coverage across sites was variable and allele frequencies had higher variance than the simplistic simulations. The variance introduced in these simulations is meant to resemble noise introduced by sequencing errors and data processing errors (e.g., mapping errors). We simulated 60 different coverage amounts for each ploidal level; these simulations varied in the minimum and maximum coverage, as well as the expected number of samples within an interval, or lambda. Based on the minimum and maximum coverage, as well as the expected number of events (lambda), the total coverage for each site is sampled from a truncated Poisson distribution, as coverage across a genome resembles a Poisson distribution with multiple peaks (Pfenninger et al., 2022). For each of our 60 simulations, we set the minimum coverage as i, maximum coverage as (i +1)*3, and lambda as half of the sum of the minimum and maximum coverage (Appendix S2: Figure S1). The resulting mean coverage simulated by this method ranged from 3 to 120x. Given a randomly selected proportion (i.e., mean and associated variance), the copies of allele A were then defined with a binomial sample with the probability defined by the beta distribution (i.e., a beta-binomial) and the copies of allele B are equal to the remainder. We then followed the data processing steps applied to real data. First, the simulated data were filtered to remove any sites where only one allele was sampled by chance. Next, we filtered the sites based on the total coverage and sequencing coverage of each allele. This function can also filter sites based on truncated allele frequencies. Finally, we randomly sampled an allele with equal probability at each site. The resulting data set includes the total coverage per site and the coverage associated with a randomly sampled allele. For the 300 simulated samples, models were evaluated at six different numbers of sites, or the total number of SNPs (1250, 2500, 5000, 10000, 20000, 25000).

### Organismal data

We applied our model to real datasets available for samples of *Saccharomyces cerevisiae, Phytophthora infestans, Glycine* spp., *Larrea tridentata,* and *Galax urceolata*; for simplicity, we refer to these as yeast, oomycete, *Glycine* spp., *Larrea*, and *Galax*, respectively. Both the yeast and oomycete sample sets were included in nQuire (Weiß et al., 2018); thus, we chose to investigate these samples with nQuack. The type of DNA sequence data varied across these samples, including whole-genome resequencing, genotype-by-sequencing (Elshire et al., 2011), target enrichment, and RadCap data (Hoffberg et al., 2016; Bayona-Vásquez et al., 2019). RadCap (Hoffberg et al., 2016; Bayona-Vásquez et al., 2019) combines reduced-representation 3RAD library preparation (Hoffberg et al., 2016; Bayona-Vásquez et al., 2019) with probe-based target capture. These sample sets also vary in the number of samples, diversity in ploidal level, taxonomic diversity, and quality of the reference genome (Table 1, Appendix S3, Appendix S4).

We aligned reads from each sample to the associated reference genome for that species (Appendix S3) with bwa-mem2 version 2.2.1 (Vasimuddin et al., 2019), converted the .sam file to a .bam file, and sorted the results with samtools version 1.15 (Danecek et al., 2021). We identified and masked repeat regions with repeatmodeler version 2.0 (Flynn et al., 2020) and repeatmasker version 4.1.1 (Smit et al., 2015). Repetitive regions should be removed from alignments before the estimation of ploidal level, as these regions will have high coverage and will likely not represent the copy number variation found in coding or single-copy regions. Based on the masked genomes, we then created databases of repeat regions that were removed from each sample alignment. We also removed poorly mapped reads and any sites that had a 10% chance or more of being mapped to the wrong location (-q 10).

To allow multiple filtering approaches to be investigated, we first prepared a text file of the .bam alignment. After preparing text files with our function prepare_data(), we manually inspected each data set and specified the minimum filtering settings accordingly. Filtering strategies differed in minimum coverage and maximum coverage quantile, as well as the lower bound (C_L_) and upper bound (C_U_) for allele frequency truncation. For all filtering strategies, sequencing depth per allele was filtered based on a sequencing error rate of 0.01, where the coverage of each allele must be more than the total coverage times the error rate, but less than the total coverage times one minus the error rate. To avoid enhancement of signal from data duplication, we randomly sample an allele with equal probability at each site. After filtering, the resulting data set includes the total coverage per site and the coverage associated with a randomly sampled allele.

We examined four filtering strategies across sample sets, with at least two examined per set. For all sample sets, we examined the minimum filtering approach (D1) and the maximum filtering approach (D4). Because hexaploid samples are expected to have mixtures with means equal to 0.17 and 0.83, we investigated filtering approaches that differed in C_L_ and C_U_, to ensure we did not remove these peaks in our filtering process. The minimum filtering approach (D1) settings differed per sample set, with three groups of settings: yeast and oomycete, *Galax* and *Glycine* spp., and *Larrea*. Respectively, the settings for the minimum filtering approach were minimum coverage equal to 10, 2, and 3, maximum coverage quantile equal to 0.90, 0.90, and 1, C_L_ equal to 0.11, 0.1, and 0.11, and C_U_ equal to 0.89, 0.9, and 0.89. The maximum filtering approach (D4) represents nQuire’s default settings, where minimum coverage is 10, C_L_ is 0.15, C_U_ is 0.85, and there is no maximum coverage cutoff. The maximum filtering approach (D4) was applied with nQuire’s create function on all samples except for the *Larrea* sample set, which was prepared with a maximum depth quantile of 0.9 and error correction of 0.01. For *Galax* and *Larrea*, we examined two additional filtering approaches to examine the intermediate between the minimum and maximum filtering approaches. First, we increased the minimum coverage to 10, but retained the C_L_ and C_U_ in the minimum filtering approach (D2). Second, we increased our allele truncation with C_L_ as 0.15 and C_U_ as 0.85, with the minimum coverage retained from the minimum filtering approach (D3). After filtering, the resulting data set includes the total coverage per site and the coverage associated with a randomly sampled allele.

### Model performance on simulated data

Overall, we found that no single model correctly assigned ploidal levels to all simulated samples (Figure 3). The amount of random noise in simulated data influenced which model correctly predicted the most simulated samples, with the best model differing for the simplistic and realistic simulated data (Appendix S5). When considering all five potential ploidal levels, the most accurate model for the simplistic simulated samples was the beta distribution with variance free and a uniform mixture. For this model, the first three ploidal levels can be differentiated at about 20x coverage; however, pentaploid and hexaploid samples cannot be differentiated until about 70x coverage. For the realistic simulated samples, when considering all five potential ploidal levels, the most accurate model is the beta-binomial with alpha free. For this model, diploids, triploids, tetraploids, and pentaploids can be differentiated at 30x coverage, but hexaploids cannot be accurately identified until 70x coverage.

**Figure 3.**
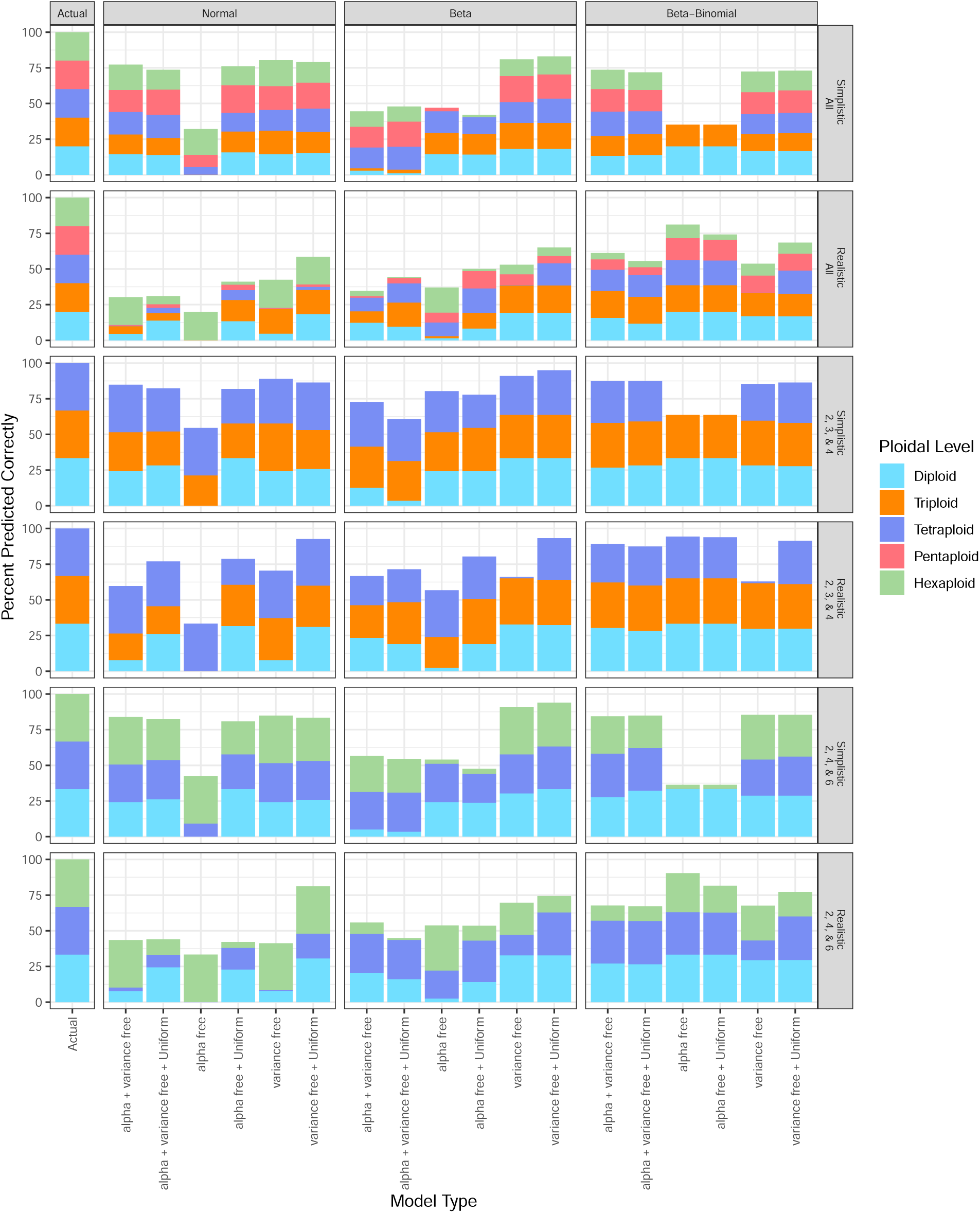
Variance in simulated data leads to a higher rate of incorrect ploidal level assignment. A larger percentage of samples will be properly assigned ploidal level when the number of mixtures examined is reduced. Some models are unsuitable for assigning specific ploidal levels, for example, diploids are not identified under the normal distribution when alpha is free.

Decreasing the number of ploidal levels considered may allow the proper assignment of ploidal levels to both simplistic and realistic samples (Figure 3). For example, when considering all ploidal levels with a normal distribution with variance free and uniform mixture, tetraploid realistic samples are identified incorrectly as hexaploids (Appendix S5: Figure S7). However, when a subset of mixtures is considered, tetraploids can be properly assigned as tetraploids for both simulation types (see Appendix S5: Figure S25 and Figure S43). The impact on sequence coverage requirements is minimal (see Appendix S5).

In some instances, we found the probability of the correct model choice to increase with the BIC difference between the best and second-best models; however, accuracy and BIC score difference often do not have a linear relationship (Appendix S6). We therefore caution against interpreting the difference in BIC scores between the best and second-best models as a measure of confidence or accuracy.

### Model performance on sample sets

As found with our simulated data, a single model was not ideal for all real samples. However, we were able to identify models that assigned ploidal level correctly to all samples or a large subset of samples for all data sets, with the best model for each sample set having at least 78% accuracy (Figure 4; Table 2). For those sample sets without pentaploid or hexaploid samples, we considered only diploid, triploid, and tetraploid mixtures, as this reduced assignment error. Our implementation of nQuire, as well as the best model identified with nQuack, had equal or greater accuracy than the original nQuire model (Table 2).

**Figure 4.**
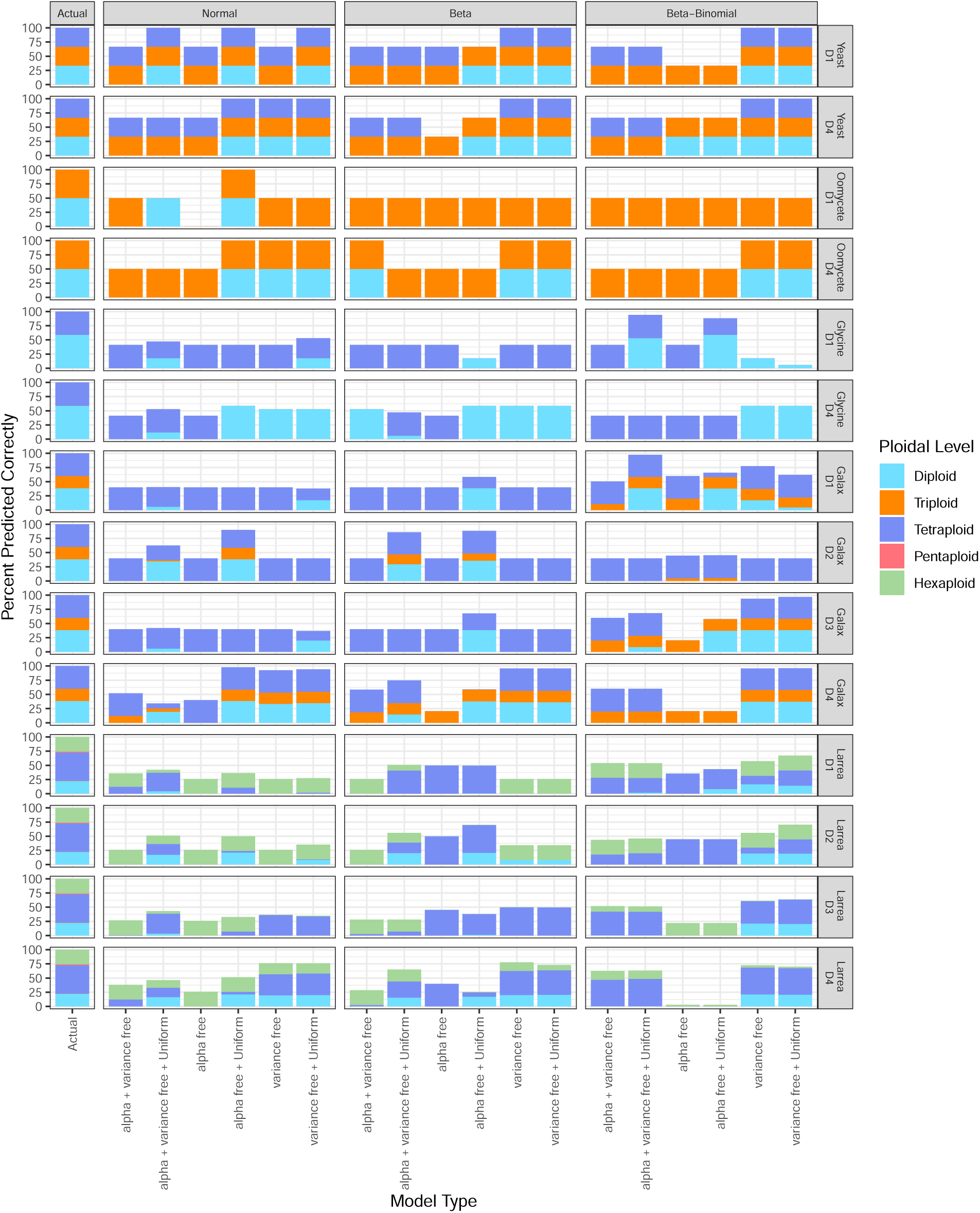
A large proportion of samples can be properly assigned ploidal level when only considering a subset of mixtures (2*x*, 3*x*, and 4*x* for yeast, oomycete, *Glycine*, and *Galax*; 2*x*, 4*x*, and 6x for *Larrea*). All samples were properly identified by at least one model for both yeast and oomycete sample sets. For *Glycine* and *Galax*, the best model identified 16 out of 17 samples and 186 out of 190 samples respectively. For *Larrea*, the best model was unable to identify 60 samples, for a total of 210 out of 270 samples correctly identified.

**Table 2.** Accuracy of nQuire compared to nQuack’s implementation of nQuire (normal distribution with variance free and a uniform mixture with the maximum filtering approach, D4), and the best model by nQuack. Percent of total samples accurately assigned in parentheses. nQuire was run on alignments before our recommended preprocessing steps.

| **Sample set** | **Total** | **nQuire** | **nQuack’s**<br>**nQuire** | **nQuack’s best**<br>**model** |
| --- | --- | --- | --- | --- |
| Yeast | 3 | 3 (100%) | 3 (100%) | 3 (100%) |
| Oomycete | 2 | 2 (100%) | 2 (100%) | 2 (100%) |
| *Glycine* spp. | 17 | 9 (53%) | 9 (53%) | 16 (94%) |
| *Galax* | 190 | 172 (91%) | 179 (94%) | 186 (97%) |
| *Larrea* | 270 | 189 (70%) | 205 (76%) | 210 (78%) |

We were able to properly assign ploidal levels to all five samples originally investigated by nQuire. For the yeast sample set, all three distributions had multiple model types that were able to properly assign ploidal level to all samples under both filtering approaches; the model type implemented in nQuire, variance free with a uniform mixture, was also able to accurately assign ploidal level to all samples with all three distributions. Notably, the normal distribution with alpha and variance free and a uniform mixture was only suitable when the allele truncation was the least constrained (D1). For the oomycete sample set, only one model was suitable when allele truncation was the least constrained: the normal distribution with alpha free and a uniform mixture. Surprisingly, for the oomycete sample set, the nQuire model type (variance free with a uniform mixture) was unable to properly assign ploidal level to the diploid sample when filtering did not match the filtering approach of nQuire. Additionally, the nQuire filtering approach (D4) allowed the proper assignment of both oomycete samples by at least two models from each distribution. Unlike all other sample sets, the maximum filtering approach (D4) increased the number of sites for both oomycete and yeast sample sets (Appendix S7); this is likely due to an excess of sites with high sequencing depth.

For *Glycine* spp., the nQuire filtering approach had low accuracy for all models (< 60 %); however, the minimum filtering approach allowed 16 of 17 samples to be assigned the correct ploidal level based on the beta-binomial distribution and the alpha- and variance-free model with a uniform mixture. We expected the alpha free model to be the best model for *Glycine* spp. samples due to the history of ancient polyploidization in *Glycine* spp. and the likely return to disomic inheritance (Walling et al., 2006), thus the proportions of each different heterozygote should be unequal. As expected, alpha as a free variable was informative for tetraploids; however, without a uniform mixture, diploids were incorrectly identified. Under the best model, the single incorrectly assigned diploid was an individual of *Glycine tomentella* (D5Bb) which is known to have a history of introgression (see Landis and Doyle, 2023). Hybridization can lead to an increased gene copy number; therefore, a more conserved filtering approach to only retain single-copy loci may be necessary to improve accuracy.

The best model for *Glycine* spp. also had high accuracy for *Galax* samples under the minimum filtering approach with 185 of 190 samples with properly assigned ploidal levels with only two tetraploids and three triploids misidentified. The tetraploid samples that were incorrectly identified had weak support; the absolute difference between the BIC score of the best model relative to the second-best model was less than 10, and these values were less than the BIC score difference of all accurate estimates. Although we caution against the interpretation of BIC score difference as a measure of accuracy generally, evaluating this method on samples with known ploidal level identified this potential usage for a set of unknown samples. When sample sequence data are more similar to the modeled data-generating process, these criteria may be informative. Here, we targeted single-copy loci with capture-based sequencing, thus avoiding variance among loci that would skew these models. However, BIC score differences were not informative for the incorrectly assigned triploid samples. Two of these three triploid samples were incorrectly identified by all models; both samples have a high abundance of sites with an allele frequency of approximately 0.5, suggesting unequal gene loss and retention across targeted sites. When low coverage sites remained (D1 & D3), the distribution with the best model remained the beta-binomial with 184 of 190 samples correctly predicted under the variance-free with uniform mixture model. When low-coverage sites were removed (D2 & D4), the best model shifted to the normal distribution with alpha free and a uniform mixture. The highest accuracy was found under the D4 filtering approach with the normal distribution with alpha free and a uniform mixture; this model accurately assigned ploidy to 186 of the 190 individuals, only failing to identify a single tetraploid and three triploids.

For the *Larrea* dataset, we were able to identify all triploids, tetraploids, pentaploids, and hexaploids under at least one model; however, the best model and filtering approach for each ploidal level differed. Based on the 18 different models and 4 different filtering approaches investigated for all cytotypes or only a subset of ploidal levels (2*x*, 4*x*, and 6*x*), we identified 22 and 39 instances, respectively, where all hexaploids were assigned the correct ploidal level. For tetraploid samples, all individuals were correctly identified in 2 instances for all cytotypes and 2 instances for only a subset of ploidal levels. Similar to the triploids in the *Galax* sample set, there were multiple diploid samples that our implemented models failed to identify correctly. These diploid samples were found to occur in mixed ploidal sites or at the edge of the species range, suggesting that ongoing mixed-ploidy introgression or divergence from the reference may skew the models’ ability to accurately assign ploidal levels due to increased gene copy number or mapping error, respectively. When considering all five potential ploidal levels, the best model was the beta distribution with alpha free and a uniform mixture, with 189 of 270 samples correctly assigned ploidal level under the D2 filtering approach; this prediction misidentified all hexaploids and pentaploids, as well as three tetraploids and four diploids. When we reduce the mixture of ploidal levels considered to only include diploids, tetraploids, and hexaploids, the best model shifts to the beta distribution with variance free under the maximum filtering approach with 210 samples correctly identified; the misidentified samples include all triploids and pentaploids, six diploids, 20 tetraploids, and 29 hexaploids. The original nQuire model was unable to estimate the correct ploidal level for only 6 diploids and 1 tetraploid from the diploid, triploid, and tetraploid *Larrea* samples; comparatively, our implementation of nQuire incorrectly assigned ploidal level to an increased number of tetraploid samples due to the inclusion of a hexaploid mixture model, which was identified as more likely for these samples. Alhough reducing the ploidal levels considered can increase the number of correctly assigned samples, we do not advise ignoring the presence of triploid, hexaploid, or pentaploid cytotypes in a system to improve model accuracy. Based on the 18 different models and 4 different filtering approaches investigated, we identified 22 and 39 instances where all hexaploids were identified correctly, when all cytotypes or only a subset of cytotypes were included. Overall, our approach increased the *Larrea* sample set accuracy compared to nQuire by 8% (Table 2).

## CONCLUSION

Here we provided expanded tools and implementations to improve site-based heterozygosity inferences of ploidal level. nQuack provides data preparation guidance and tools to decrease noise in input data. These tools include a maximum sequence coverage quantile filter and sequence error-based filter to remove biallelic sites that are likely not representative of copy number variance in the nuclear genome. We also consider only the frequency of allele A or B at each site, instead of both, as done in nQuire, as this would inflate the observation by enhancing the signal or noise found in the data. Our model builds upon, and improves, the nQuire framework by extending it to higher ploidal levels (pentaploid and hexaploid), correcting the augmented likelihood calculation, implementing more suitable probability distributions, and allowing additional ‘fixed’ models. We also decrease model selection errors by relying on BIC rather than likelihood ratio tests (Taper et al., 2021).

Through the intensive testing of our proposed methods, we found that many variables influence model accuracy. Based on our simulated data, we observed that each model implementation and model type can be influenced by the number of sites, sequencing coverage, and amount of variance or noise in a dataset. In real data sets, this noise can be introduced through sequencing error or general sequence mapping error. In addition, although we attempted to retain only single-copy loci by removing repetitive regions, additional filtering may increase accuracy to ensure estimates are not conflated by variation among loci at non-single-copy sites. By examining a large amount of real data, we determined that the most accurate model for each data set differed, suggesting that both filtering strategies and model selection must be explored on a set of known samples before applying these models to any sample with an unknown ploidal level to achieve accurate ploidal-level assignment.

We explored nQuack’s performance on an extensive set of simulated data and multiple real-world datasets. These analyses allowed us to benchmark model performance and identify data features that affect nQuack’s predictive power. However, the biological datasets we explored cannot represent the full diversity of polyploid systems, and additional tuning is required for real datasets. For example, these models would not be suitable in an allotetraploid with strict disomic inheritance as no AAAB or ABBB loci would occur; therefore, the most likely model could be identified as a diploid, though BIC score parameter corrections would allow the most probable model to be hexaploid or tetraploid. Additional biological systems will likely introduce more complexities and may work best under different filtering conditions. To identify which factors dictate which strategy is the most accurate, multiple mixed-ploidy systems with high-quality reference genomes, well-classified polyploidization events (e.g., mode of formation, timing of polyploidy events, chromosomal segregation patterns, etc.), and well-characterized reproductive history should be explored in future model iterations. Regarding summary statistics, non-parametric bootstrapping after model selection would allow for assessing the strength of the evidence in favor of every model and the robustness of model selection results. We provide functions to perform this non-parametric bootstrap sampling, however, completing a full non-parametric bootstrap for all of our real datasets was neither practical nor feasible due to computational limitations. Because all mathematical models are misspecifications of the true data-generating process (Dennis et al., 2019), errors are probable when selecting the model closest to the truth. Therefore, by resampling the data we can assess the reliability of the model choice. In addition, if analytical-based inferences continue to be pursued, a sliding window approach will likely improve ploidy inferences.

Our results open many interesting avenues for future research. Site-based heterozygosity models like the ones used here are in essence phenomenological statistical models, which focus on reproducing patterns rather than generating patterns based on a fundamental biological process. Although statistical models embodying fundamental biological processes are common in many areas of biology (for instance, in phylogenetics), in this particular case it is extremely difficult to capture the complexities of nature in an analytical-based inference, and future model exploration utilizing data-based inference to classify ploidal levels is warranted. Alternatively, demographic models like the ones we proposed elsewhere (Gaynor et al., 2023) may provide the ecological and evolutionary framework necessary to design process-based predictions for mixed ploidy. These models, however, require rigorous coupling with evolutionary and genomic theory.

Overall, this analysis reveals that it is critical to thoroughly examine proposed methods before inferring biological meaning. nQuack, as well as nQuire, should not be used to infer the ploidal level in a system for which very little is known, as these models are often positively misleading. We also suggest caution when relying on any site-based heterozygosity to predict ploidal level of a sample even when a known dataset is analyzed before applying the method to a sample of unknown ploidy due to the potential impact of biological processes (e.g., hybridization, divergence, etc.,) on model inference. Despite the caveats to this method, it can be easily implemented to leverage sequence data for ploidal estimation.

## ACKNOWLEDGMENTS

This research was supported by the NSF Graduate Research Fellowship (DGE-1842473) to M.L.G., an NSF Small Grant (DEB-1556371) to R.G.L, an NSF Plant Genome Fellowship (IOS-1711807) to J.B.L., and a USDA NIFA Hatch award (7002754) to J.J.D. We thank William Weaver, Eric Goolsby, and Matthew Gitzendanner for assistance with C++. We thank Alyssa Philips, Trevor Faske, and Matthew G. Johnson for their feedback and discussion.

## AUTHOR CONTRIBUTIONS

Original conceptualization by M.L.G, D.E.S, J.M.P, and P.S.S. Methodology designed by M.L.G and J.M.P. Software and formal analysis was written and conducted by M.L.G. Data were generated by M.L.G, J.B.L, J.J.D, T.K.O, and R.G.L. Original draft and visualization by M.L.G. All authors reviewed and contributed to the final manuscript.

## DATA AVAILABILITY STATEMENT

The R package nQuack is available https://github.com/mgaynor1/nQuack and https://mlgaynor.com/nQuack/. A full implementation tutorial (https://mlgaynor.com/nQuack/articles/BasicExample.html), as well as detailed tutorials on data preparation (https://mlgaynor.com/nQuack/articles/DataPreparation.html) and model inference (https://mlgaynor.com/nQuack/articles/ModelOptions.html), are available with the package documentation. For three sample sets, reference genomes and population genetics data are available via open repositories (see Appendix S3 and S4 for accessions). Sequence data for *Galax urceolata* and *Larrea tridentata* will be published in open repositories with future publications. An example data set, as well as the output of each step of our method, is available on our github (https://github.com/mgaynor1/nQuack/tree/main/data).

## SUPPORTING INFORMATION

Additional Supporting Information may be found online in the Supporting Information section at the end of the article.

Appendix S1. Detailed implementation of expected maximization algorithm and the models available in our method.

Appendix S2. Distribution of coverage for the realistic simulation approach (Figure S1).

Appendix S3. Genome statistics including species identity, number of contigs, total length in basepairs, contig minimum length, contig average length, contig maximum length, N50 (Mb), percent of GC content, BUSCO complete percentage, BUSCO duplicate percentage, BUSCO reference, and any accession information available.

Appendix S4. Extended information on sample sets including the number of samples of each ploidal level, the sequencer used, and any accession information available.

Appendix S5. Model comparisons for simulated data sets when considering all ploidal levels (Figure S2 - S19) or only a subset of ploidal levels: (1) only diploid, triploids, and tetraploids (Figure S20 - S38), or (2) only diploid, tetraploids, and hexaploids (Figure S39 - S57). BIC score difference between the best and second best models for simulated samples across different numbers of sites for eleven different coverage amounts (5, 10, 20, 30, …). The color of each point represents the best model. The shape of each point represents the approach used to simulate that sample. A larger BIC difference between the best and second-best models indicates model confidence. These plots can be used to guide users’ interpretation of these models and determine if these models will apply to their system.

Appendix S6. Probability of the correct model choice given the BIC difference between the best and second best model for all simulated diploid, triploid, tetraploid, pentaploid, and hexaploid samples. The probability of success was predicted based on a logistic regression where accuracy is a function of BIC difference. We expect the BIC difference between the best and second best model to increase with the probability of success when the BIC difference is indicative of the model’s accuracy. (Figure S58 - S60)

Appendix S7. Number of sites and mean sequence coverage included for all filtering approaches for each sample set. (Figure S61)

