## Appendix S1 for "nQuack: An R package for predicting ploidal level from sequence data using site-based heterozygosity"

### 1 Methods

Below we detail our implementation of the expected maximization (EM) algorithm for the normal, the beta, and the beta-binomial distributions. A major difference between these implementations is the necessary allele properties; for the normal and beta implementations, only allele frequencies are needed, while total coverage and allele sequencing coverage are needed for the beta-binomial. For simplicity, we first introduce our implementation with the normal and beta distribution.

#### 1.1 Statistical model for the observed biallelic heterozygosity

Let $n$ be the number of sites in the genome where biallelic heterozygosity is observed. Let $X_1, X_2, \ldots, X_n$ be the randomly sampled allele properties at each site; this may be the relative allelic frequencies or the sequencing depth of the allele. *We simply refer to these as allelic frequencies to begin.* The general statistical model assumes that the $i^{\text{th}}$ observation $X_i$ $(i = 1, \ldots, n)$ is a sample from a mixture distribution with $R$ components. The number of mixture components, $R$, is determined by types of biallelic heterozygotes possible at each ploidal level (Table 1;*i.e.*, diploid = AB, triploid = AAB or ABB, tetraploid = AAAB, AABB, or ABBB). For example, we expect one mixture component with a mean ($\mu$) of 0.5 for a diploid (50% allele A, 50% allele B). Here, we assume the mixture components each follow a normal, beta, or beta-binomial distribution, and fit five different statistical models: we consider one model per ploidal level (see Table 2). For the sake of generality and to accommodate any probability distribution, we let $f_r(x_i)$ denote the probability density function (pdf) of the $r^{\text{th}}$ mixture component.

Since $f_r(x_i)$ omits the specification of the parameters, we let $\boldsymbol{\theta}$ denote the vector whose elements are all the parameters of these mixture components. In addition to mean $\mu_r$ and variance $\sigma_r$, we denote the mixture proportions as $\alpha_r$, where $r = 1, 2, \ldots, R$ is defined such that $\sum_{r=1}^{R} \alpha_r = 1$. Therefore, $\boldsymbol{\theta}$ is defined as

$$\boldsymbol{\theta} = [\alpha_1, \alpha_2, \ldots, \alpha_R, \mu_1, \mu_2, \ldots, \mu_R, \sigma_1, \sigma_2, \ldots, \sigma_R].$$

Note that each of the $f_r(x_i)$ depends on a subset of $\boldsymbol{\theta}$.

The likelihood function needed for parameter estimation is defined as the joint probability density functions of the observations, $f(x_i)$ where $i = 1, 2, \ldots, n$:

$$L(\theta) = \prod_{i=1}^{n} f(x_i) = \prod_{i=1}^{n} \sum_{r=1}^{R} \alpha_r f_r(x_i). \tag{1}$$

The log-likelihood function is:

$$\ln L(\theta) = \sum_{i=1}^{n} \ln f(x_i) = \sum_{i=1}^{n} \ln \left[ \sum_{r=1}^{R} \alpha_r f_r(x_i) \right]. \tag{2}$$

#### 1.2 The Expectation Maximization (EM) algorithm

The EM algorithm is a technique for calculating the maximum likelihood estimates (MLEs) of the model parameters [3]. Here, we use the data augmentation algorithm to construct a joint likelihood function including the observations $X_i$, where $i = 1, 2, \ldots, n$, and latent variables. The latent variables are denoted $Z_i$ (defined below for our implementation) and the joint likelihood of the observations and the latent variables as $L(X_i, Z_i) = f(X_i, Z_i)$. The algorithm is implemented in two steps: (1) the 'E-step" and (2) the "M-step".

First, in the "E-step", all possible values of the missing data are weighted by their conditional probability given the current observations, $f(Z_i|X_i)$. In practice, such weighting amounts to calculating the expected augmented log-likelihood, averaged over this conditional distribution, or

$$E_{Z|X} \left[ \ln L(X_i, Z_i) \right].$$

The second step, the "M-step", consists of maximizing the expected log-likelihood.

The latent variables in our case are the allelic frequencies data, and the key missing information is which partition of the mixture each observation $X_i$ is comes. Therefore, we let $Z_i$ denote the mixture component from which the $i^{\text{th}}$ observation comes. Here, $Z_i$ can take on the values $1, 2, \ldots, R$. We then define a general augmented log-likelihood that can accommodate many special cases. We do so using an indicator random variable $I_{\{A\}}$ which is a Bernoulli random variable that takes on only two values, 1 if $A$ is true and 0 otherwise. Here, we will define the indicator function as follows: Let

$$I_{\{Z_i = r\}} = \begin{cases} 1 & \text{if} \quad Z_i = r \\ 0 & \text{otherwise.} \end{cases}$$

Note that $E_{Z|X} \left[ I_{\{Z_i = r\}} \right] = P(Z_i = r | X_i)$. This probability is in turn found using Bayes' theorem:

$$
\begin{aligned}
P(Z_i = r|X_i) &= \frac{P(X_i|Z_i = r)P(Z_i = r)}{P(X_i)} \\
&= \frac{f_r(x_i)\alpha_r}{f(x_i)} \\
&= \frac{f_r(x_i)\alpha_r}{\sum\limits_{r=1}^{R} \alpha_r f_r(x_i)} \equiv p_{ir}.
\end{aligned} \tag{3}
$$

With these observations and definitions in hand, the augmented data likelihood is then written as

$$
L(X_i, Z_i) = f(X_i, Z_i) = \prod_{i=1}^{n} \prod_{r=1}^{R} \alpha_r^{I_{\{Z_i=r\}}} f_r(x_i)^{I_{\{Z_i=r\}}}, \tag{4}
$$

so that the expected augmented log-likelihood becomes

$$
\begin{aligned}
E_{Z|X}\left[\ln L(X_i, Z_i)\right] &= E_{Z|X}\left[\sum_{i=1}^{n}\sum_{r=1}^{R} I_{\{Z_i=r\}}\left(\ln \alpha_r + \ln f_r(x_i)\right)\right] \\
&= \sum_{i=1}^{n}\sum_{r=1}^{R} E_{Z|X}\left[I_{\{Z_i=r\}}\right]\left(\ln \alpha_r + \ln f_r(x_i)\right) \\
&= \sum_{i=1}^{n}\sum_{r=1}^{R} p_{ir}\left(\ln \alpha_r + \ln f_r(x_i)\right).
\end{aligned} \tag{5}
$$

Finally, to implement the EM algorithm, one chooses initial values of $\boldsymbol{\theta}$, $\boldsymbol{\theta}^c$, where the superscript $^c$ stands for "current", then uses them to calculate $p_{ir}$ (eq.3). Denote this quantity as $p_{ir}^c$. Plug $p_{ir}^c$ into the final expression in eq. 8 and maximize it with respect of $\boldsymbol{\theta}$ to obtain updated values of the parameter estimates, $\boldsymbol{\theta}^{c+1}$. Next, use these updated estimates $\boldsymbol{\theta}^{c+1}$ to calculate an updated value of $p_{ir}$, denoted $p_{ir}^{c+1}$. Plug $p_{ir}^{c+1}$ into the augmented likelihood eq. 8 and maximize it again with respect to $\boldsymbol{\theta}$ to obtain a new set of updated values of the model parameters. Finally, repeat this process until $\boldsymbol{\theta}^c \approx \boldsymbol{\theta}^{c+1}$, or equivalently, until $|E_{Z|X}\left[\ln L(X_i, Z_i); \boldsymbol{\theta}^c\right] - E_{Z|X}\left[\ln L(X_i, Z_i); \boldsymbol{\theta}^{c+1}\right]| < \epsilon$, where $\epsilon$ is the convergence tolerance. Finally, note that at each iteration of this process, the maximization of eq. 8 is done numerically with a Nelder-Mead simplex optimization algorithm [4].

###### 1.2.1 EM algorithm for a mixture of Normal Distribution

The expected augmented log-likelihood eq. (8) in the normal mixture model can be maximized analytically. Notably, for the normal mixture model the MLE matches the moment estimates for the means. Starting, $p_{ir}^{(0)}$ is computed based on an initial value of the model parameters $\boldsymbol{\theta}^{(0)}$ and an initial value of $p_{ir}$ (eq. 3). Next, we compute the derivative of the augmented log-likelihood with respect to every parameter, then set these equal to 0 and solve for the model parameters. The first values of the MLEs are then obtained. For example, to estimate the mean $\mu_1$ of the first mixture component, the partial derivative of the log-augmented likelihood with respect to $\mu_1$ becomes:

$$\begin{aligned}\frac{\partial}{\partial \mu_1} \sum_{i=1}^{n} \sum_{r=1}^{R} p_{ir}^{(0)} \left(\ln \alpha_r + \ln f_r(x_i)\right) &\propto \sum_{i=1}^{n} \sum_{r=1}^{R} \frac{\partial}{\partial \mu_1} p_{ir}^{(0)} \ln f_r(x_i) \\ &= \sum_{i=1}^{n} \sum_{r=1}^{R} p_{ir}^{(0)} \frac{\partial}{\partial \mu_1} \left[ \ln \left(2\pi\sigma_r^2\right)^{-1/2} - \frac{1}{2\sigma_r^2}(x_i - \mu_r)^2 \right].\end{aligned}$$

To obtain the first iteration of the MLE for $\mu_1$ we simplify the right hand side (RHS) of the above equation, set it equal to 0 and solve for $\mu_1$:

$$\sum_{i=1}^{n} p_{ir}^{(0)} \frac{(x_i - \mu_1)}{\sigma_1^2} = 0 \implies \hat{\mu_1} = \frac{1}{S_r} \sum_{i=1}^{n} p_{ir}^{0} x_i,$$

where $S_r = \sum_{i=1}^{n} p_{ir}^{(0)}$

The same approach is used to obtain a first pass of the estimates of all the model parameters. Subsequently, these model parameters are used to compute a new estimate of $p_{ir}$, $p_{ir}^{(1)}$, which is in turn used to update the analytical estimates of each model parameter. This process is repeated a large number of times, say $k$, until the estimates between the $k^{\text{th}}$ and the $(k+1)^{\text{th}}$ iteration differ by less than the convergence tolerance level, $\epsilon > 0$ ([5]).

By exploiting the analytical MLE solutions of the normal statistical model, the implementation works fast and is "cheap" in terms of computation. However, there are two important considerations ignored: first, the normal distribution has the entire real line as its support (from $-\infty$, to $\infty$) yet the data are only between 0 and 1. Second, this approach ignores the truncation of the observed data, which is often included in data processing steps. Allele frequencies, the $x_i$, are often filtered to between two limits $c_L, c_U$ such that $0 \leq c_L < c_U < 1$. The assumption is that minor allele frequency of 0.1 or below represents sequencing error rather than true population values. Due to the minor allele cut-off, the max allele frequency then becomes 0.9. In some instances, allele truncation may remove sites that do not represent error; in these instances the expectation should be truncated to match the observation. We account for both of these limitations in the beta and beta-binomial EM algorithm implementation.

##### 1.2.2 EM algorithm for a mixture of Beta Distribution

Here, we use a truncated generalized beta distribution likelihood for the $f_r(x_i)$. We define the traditional shape ($a$) and scale ($b$) parameters as $a = \mu[\frac{\mu(1-\mu)}{\sigma^2} - 1]$ and $b = (1-\mu)[\frac{\mu(1-\mu)}{\sigma^2} - 1]$.

If the data are originally assumed to have a standard beta distribution such that the pdf of the $r^{\text{th}}$ mixture component is $g_r(x_i) = \frac{\Gamma(a_r+b_r)}{\Gamma(a_r)\Gamma(b_r)} x_i^{a_r-1}(1-x_i)^{b_r-1}$ and its corresponding cdf is denoted $F_g(x_i)$, then the pdf $f_r(x_i)$ we needed to write the truncated likelihood function with bounds $c_L$ and $c_U$ was

$$f_r(x_i) = \frac{g_r(x_i) I_{\{c_L < x_i < c_U\}}}{F_g(c_U) - F_g(c_L)}. \tag{6}$$

The implementation of the EM proceeds as described above with the analytical maximization step with numerical optimizations of the log-likelihood in eq. 8 in which the $f_r(x_i)$ are written as in equation 6.

##### 1.2.3 EM algorithm for a mixture of Beta-Binomial Distribution

Using beta-binomial likelihoods, $X_i$ for each observation is defined as the total coverage and the allele sequencing coverage for a randomly sampled allele. In the above distributions, observations are allele frequencies; however, this observation represented a transformation of the data, therefore important properties of the original data might be lost and not be reflected in the likelihood function. In this implementation, we investigate the possibility that the variability and heterogeneity in the data are better reflected with a count data model where the mean and the variance depend on each other.

The simplest probability model of how the data arise is a binomial distribution. Under this model, the number of trials would be the total coverage or sequencing depth and the number of successes would be the number of allele A sampled. The probability of a "success" (*i.e.* observing an A) would then be the parameter of interest. However, under the binomial case, such success probability is assumed to be fixed. Here, we propose that such success probability, call it $\pi$, is a random variable and beta distributed with pdf

$$\frac{\Gamma(a_r + b_r)}{\Gamma(a_r)\Gamma(b_r)} \pi^{a_r-1}(1-\pi)^{b_r-1},$$

where $a_r > 0$ and $b_r > 0$ are the two shape parameters of this beta distribution. Integrating the binomial count model over that beta distribution for the probability of success $\pi$ gives the well-known beta-binomial distribution (*e.g.*, [1, 2]). We then use this model as the probability model for each one of the mixture components. Let $N_i$ be the depth at site $i$ and $Y_i$ be the number of allele A at site $i$. Then, the individual $f_r$'s are for site $i$ under the beta-binomial model become:

$$f_r(y_i) = \frac{\Gamma(N_i + 1)}{\Gamma(y_i + 1)\Gamma(N_i - y_i + 1)} \frac{\Gamma(y_i + a_r)\Gamma(N_i - y_i + b_r)}{\Gamma(N_i + a_r + b_r)} \frac{\Gamma(a_r + b_r)}{\Gamma(a_r)\Gamma(b_r)}.$$

###### 1.2.4 Addition of a Uniform Mixture

Each mixture model can be extended by adding a uniform noise component. To account for this additional mixture, the general log-likelihood introduced in eq. 2 can be expanded:

$$\ln L(\theta) = \sum_{i=1}^{n} \ln \left[ \alpha_{R+1} U(x_i) + \left( \sum_{r=1}^{R} \alpha_r f_r(x_i) \right) \right]. \tag{7}$$

Similarly, the augmented log-likelihood becomes:

$$E_{Z|X} \left[ \ln L(X_i, Z_i) \right] = \sum_{i=1}^{n} \left[ p_{i(R+1)} \left( \ln \alpha_{R+1} + \ln U(x_i) \right) + \left( \sum_{r=1}^{R} p_{ir} \left( \ln \alpha_r + \ln f_r(x_i) \right) \right) \right]. \tag{8}$$

###### 1.2.5 Overdispersion and Sequencing Error

Instead of defining the shape and scale of the beta distribution with $\sigma$, we are able to define these distributions with overdispersion ($\tau$) and sequencing error rate ($e$). The parameter $\tau$ is well known in the statistical literature as the parameter giving rise to overdispersion, the statistical phenomenon wherein the variance of a distribution is larger than its mean [2]. The technical details are as follows:

Instead of writing the pdf of the beta distribution for the probability of successful detection $\pi$ as $\frac{\Gamma(a_r+b_r)}{\Gamma(a_r)\Gamma(b_r)} \pi^{a_r-1}(1-\pi)^{b_r-1}$, with mean and variance

$$\begin{aligned} \mathrm{E}(\pi) &= \frac{a}{(a+b)}, \\ \mathrm{Var}(\pi) &= \frac{a}{(a+b)} \frac{b}{(a+b)} \frac{1}{(a+b+1)}, \end{aligned}$$

we re-parameterized it as a function of the mean by letting $m = a/(a+b) = \mathrm{E}(\pi)$ and $\tau = 1/(a+b+1)$. With this re-parameterization, $\mathrm{Var}(\pi) = m(1-m)\tau$. Note that there is a one-to-one correspondence between $m$ and $\tau$ and $a$ and $b$, namely:

$$\begin{aligned} a &= \frac{(1-\tau)m}{\tau}, \\ b &= \frac{(1-m)(1-\tau)}{\tau}. \end{aligned}$$

This re-parameterization then allows the introduction of sequencing error as something affecting the mean $m$ of the probability of successful detection $\pi$. If we expect that on average, a target allele will be detected a proportion $m$ of the time, then the probability of sampling that allele is equal to $p$, it follows that the estimation of the sequencing error $e$ can be accommodated by writing $m$ as a function of $e$ and $p$ as follows:

$$\begin{aligned}
\mathrm{E}(\pi) = m = & \; P(\text{allele is in sample and no error was made}) \text{ or} \\
& P(\text{allele isn't in sample and an error was made}) \\
& = P(\text{allele is in sample})P(\text{no error was made}) \\
& + P(\text{allele isn't in sample})P(\text{an error was made}) \\
& = p(1-e) + (1-p)e.
\end{aligned}$$

With this reparameterization of the beta distribution in hand, and the one-to-one transformations between $m$ and $\tau$ and $a$ and $b$, we expanded the beta-binomial model above to utilize either $\mu$ and $\sigma$ or $\mu$, $\tau$, and $e$.

The resulting re-parameterization and expansion of the beta distribution model for $\pi$ allows the estimation of sequencing error and the ability to define how sequencing error modulates the amount of overdispersion. In other words, this methodology allows not only estimation but also understanding of the effects of sequencing error in shaping the variability of the original data. Therefore, we can transform starting parameter values to account for sequencing overdispersion and sequencing error for all implementations.

##### 1.2.6 Free and Fixed Models

For all implementations, starting parameters must be defined for $\alpha$, $\mu$, and $\sigma$. We then provide 4 types of inferences that differ in which parameter is included in the estimation. 'Free' models estimate $\alpha$, $\mu$, and $\sigma$. Alternatively, "fixed" models include implementations where only $\alpha$ , only $\sigma$, or only $\alpha$ and $\sigma$ are estimated. By allowing all parameters to be predicted, as well as some parameters to be constrained, we allow model selection via information criteria, where divergence among models can be estimated by calculating the log-likelihood ratio.

##### 1.2.7 Summary Statistics

Model selection was explored based on log-likelihood and BIC. Here we define $BIC = -2\ln(L(\theta) + n_\theta \ln(n_{x_i})$, where $n_\theta$ are the number of parameters estimated and $n_{x_i}$ are the number of sites for which allele properties were sampled. In cases where free and fixed models were estimated, we defined divergence with $\Delta\ln(L_{R,fixed}) = \ln(L_{free}) - \ln(L_{R,fixed})$.

### Tables

Table 1: Types of biallelic heterozygotes and the associated relative allelic frequencies given the ploidal level.

| Ploidal Level | Type | Frequency of A | Frequency of B |
|---|---|---|---|
| Diploid | AB | 0.50 | 0.50 |
| Triploid | AAB | 0.67 | 0.33 |
| | ABB | 0.33 | 0.67 |
| Tetraploid | AAAB | 0.75 | 0.25 |
| | AABB | 0.50 | 0.50 |
| | ABBB | 0.25 | 0.75 |
| Pentaploid | AAAAB | 0.80 | 0.20 |
| | AAABB | 0.60 | 0.40 |
| | AABBB | 0.40 | 0.80 |
| | ABBBB | 0.20 | 0.80 |
| Hexaploid | AAAAAB | 0.83 | 0.16 |
| | AAAABB | 0.67 | 0.33 |
| | AAABBB | 0.50 | 0.50 |
| | AABBBB | 0.33 | 0.67 |
| | ABBBBB | 0.17 | 0.83 |

Table 2: Expected mean(s) for each ploidal level.

| Ploidal Level | $\mu$ |
|---|---|
| Diploid | 0.50 |
| Triploid | 0.67, 0.33 |
| Tetraploid | 0.75, 0.50, 0.25 |
| Pentaploid | 0.80, 0.60, 0.40, 0.20 |
| Hexaploid | 0.83, 0.67, 0.50, 0.33, 0.17 |
