## Supplementary figures and images for "nQuack: An R package for predicting ploidal level from sequence data using site-based heterozygosity"

### Appendix S2

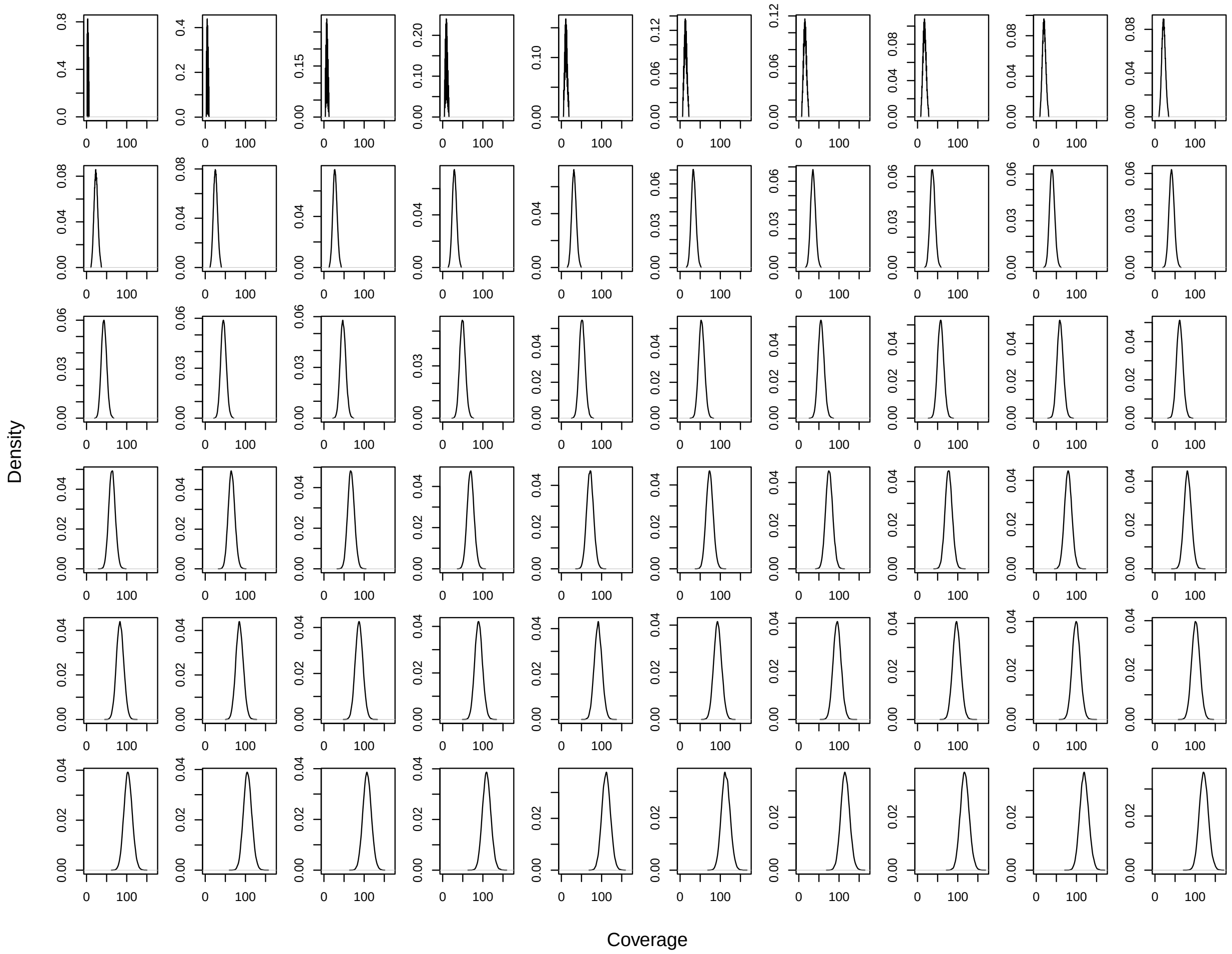


Figure S1. Distribution of coverage of the realistic simulation approach.
