## Appendix S5 for "nQuack: An R package for predicting ploidal level from sequence data using site-based heterozygosity"

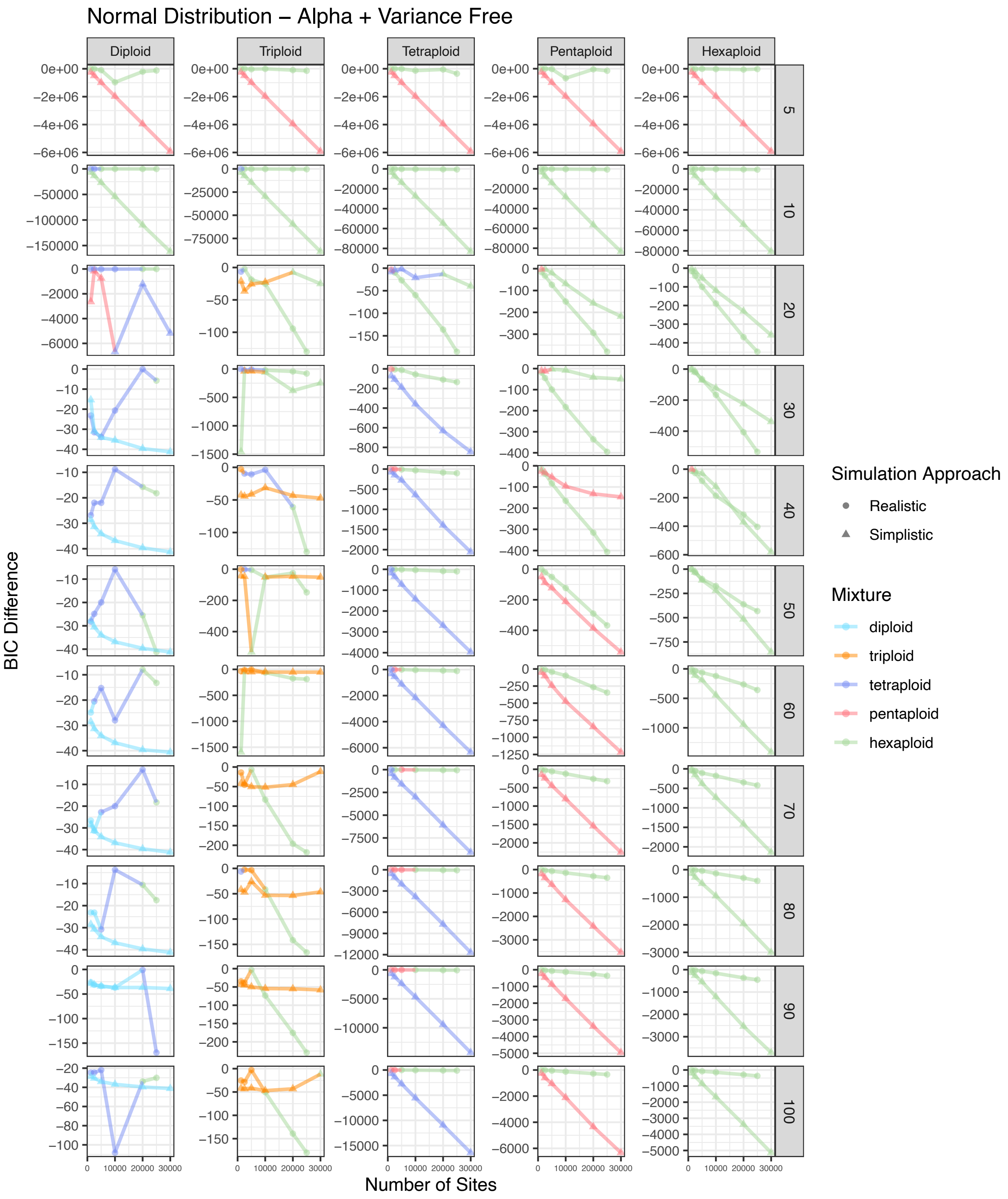


Figure S2. BIC score difference between the best and second best model for simulated diploid, triploid, tetraploid, pentaploid, and hexaploid samples across different numbers of sites for eleven different coverage amounts. The color of each point represents the best model. The shape of each point represents the approach used to simulate that sample. This represents a normal distribution with alpha and variance free.

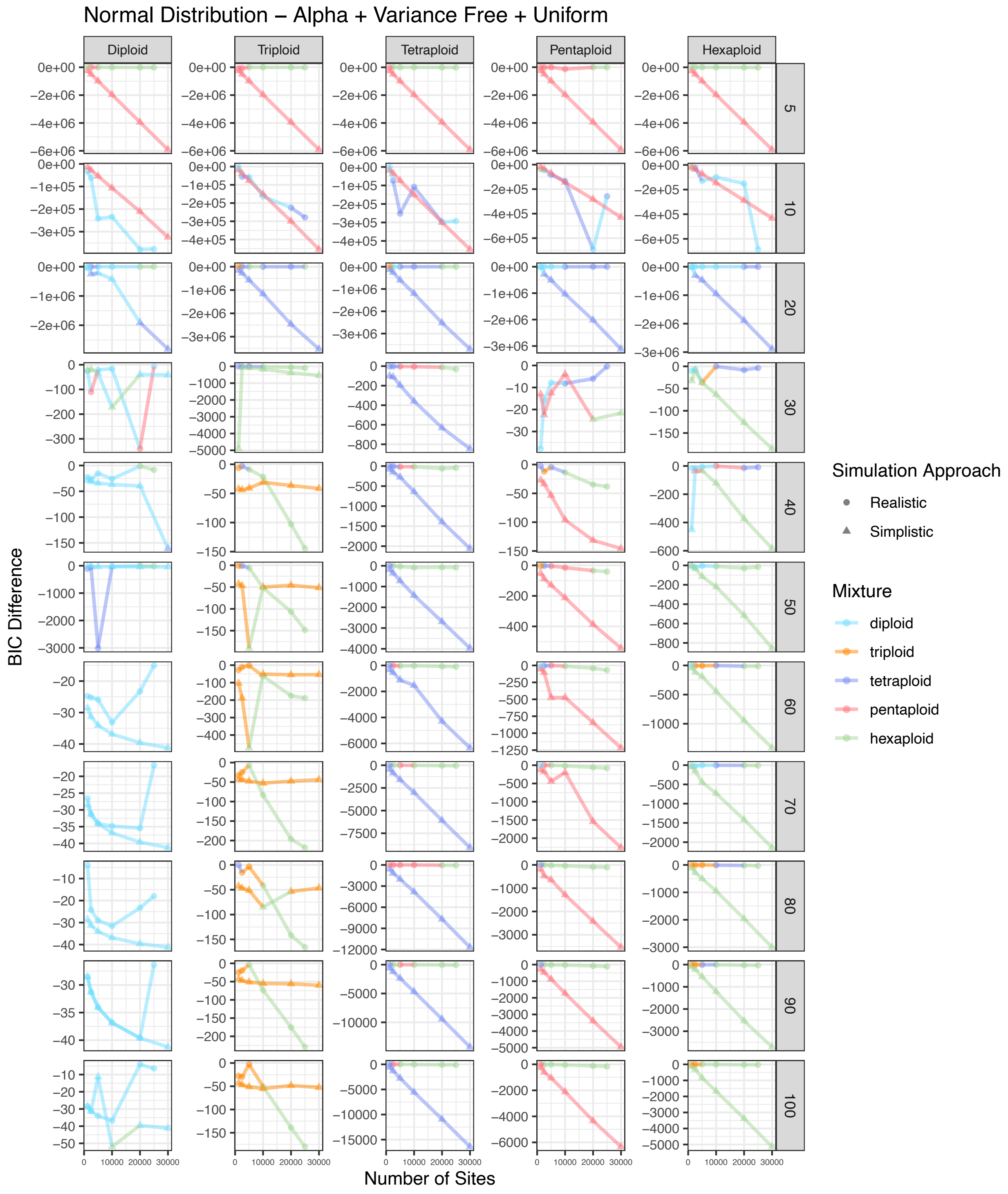


Figure S3. BIC score difference between the best and second best model for simulated diploid, triploid, tetraploid, pentaploid, and hexaploid samples across different numbers of sites for eleven different coverage amounts. The color of each point represents the best model. The shape of each point represents the approach used to simulate that sample. This represents a normal distribution with alpha and variance free with a uniform mixture.

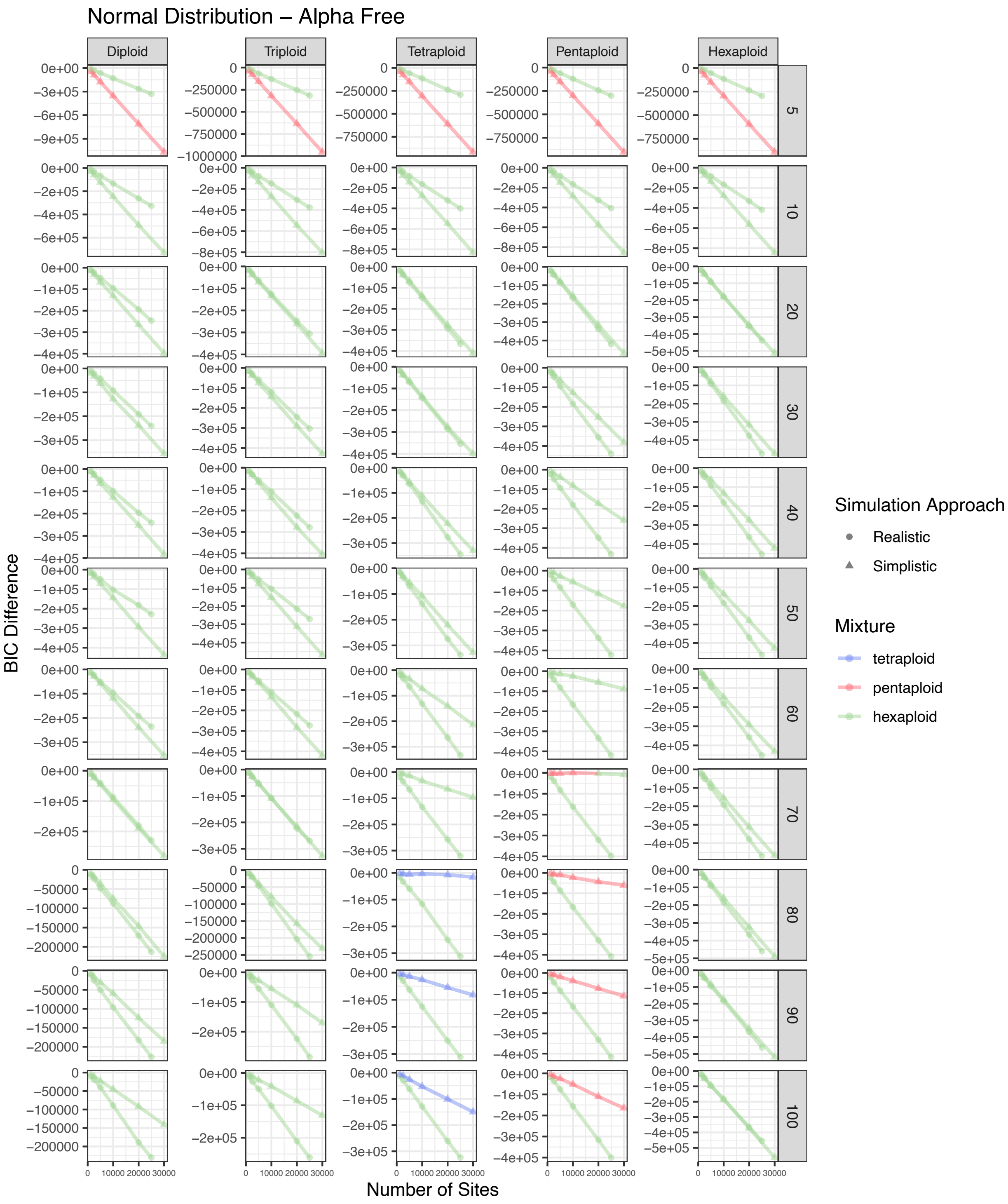


Figure S4. BIC score difference between the best and second best model for simulated diploid, triploid, tetraploid, pentaploid, and hexaploid samples across different numbers of sites for eleven different coverage amounts. The color of each point represents the best model. The shape of each point represents the approach used to simulate that sample. This represents a normal distribution with alpha free.

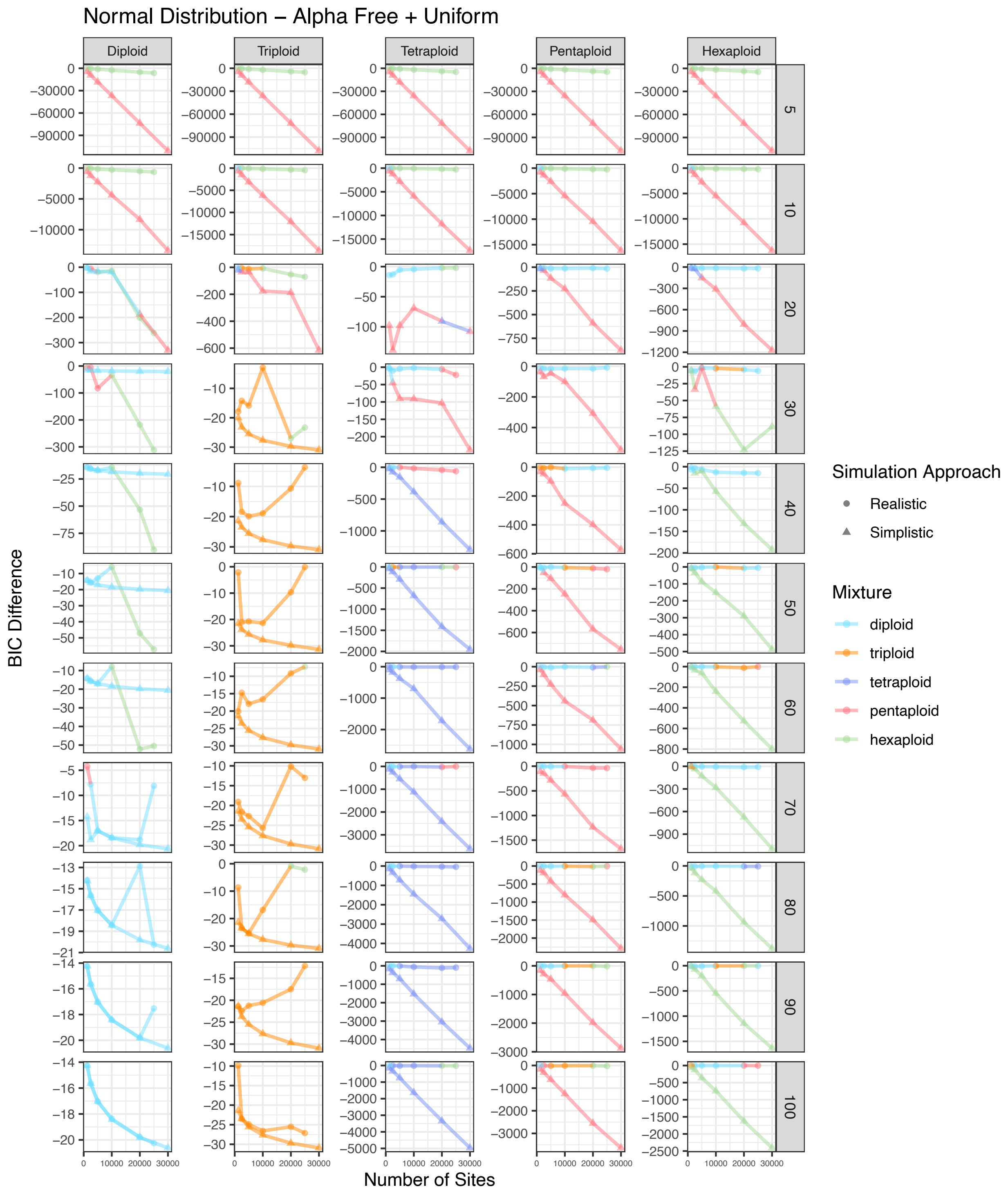


Figure S5. BIC score difference between the best and second best model for simulated diploid, triploid, tetraploid, pentaploid, and hexaploid samples across different numbers of sites for eleven different coverage amounts. The color of each point represents the best model. The shape of each point represents the approach used to simulate that sample. This represents a normal distribution with alpha free with a uniform mixture.

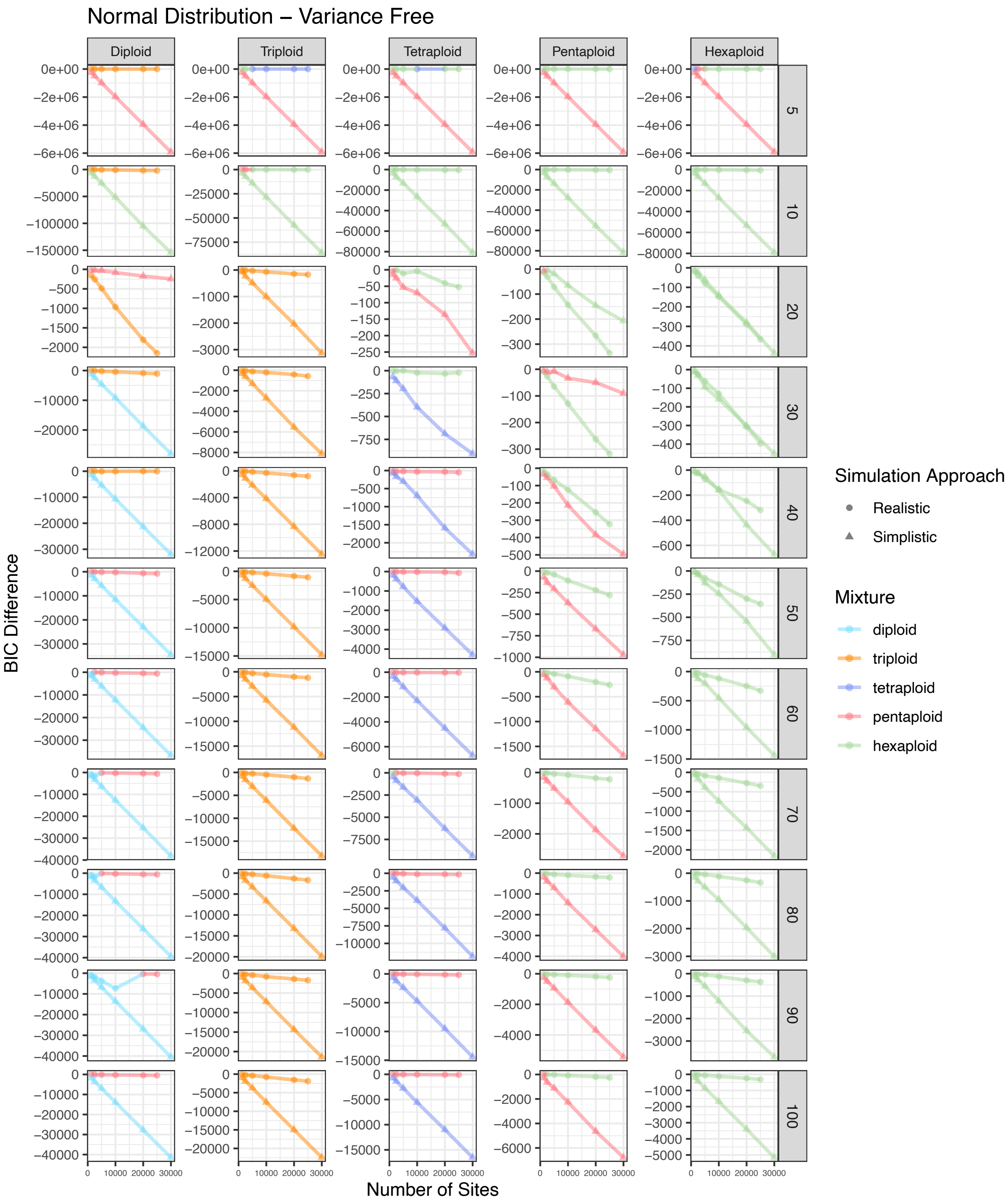


Figure S6. BIC score difference between the best and second best model for simulated diploid, triploid, tetraploid, pentaploid, and hexaploid samples across different numbers of sites for eleven different coverage amounts. The color of each point represents the best model. The shape of each point represents the approach used to simulate that sample. This represents a normal distribution with variance free.

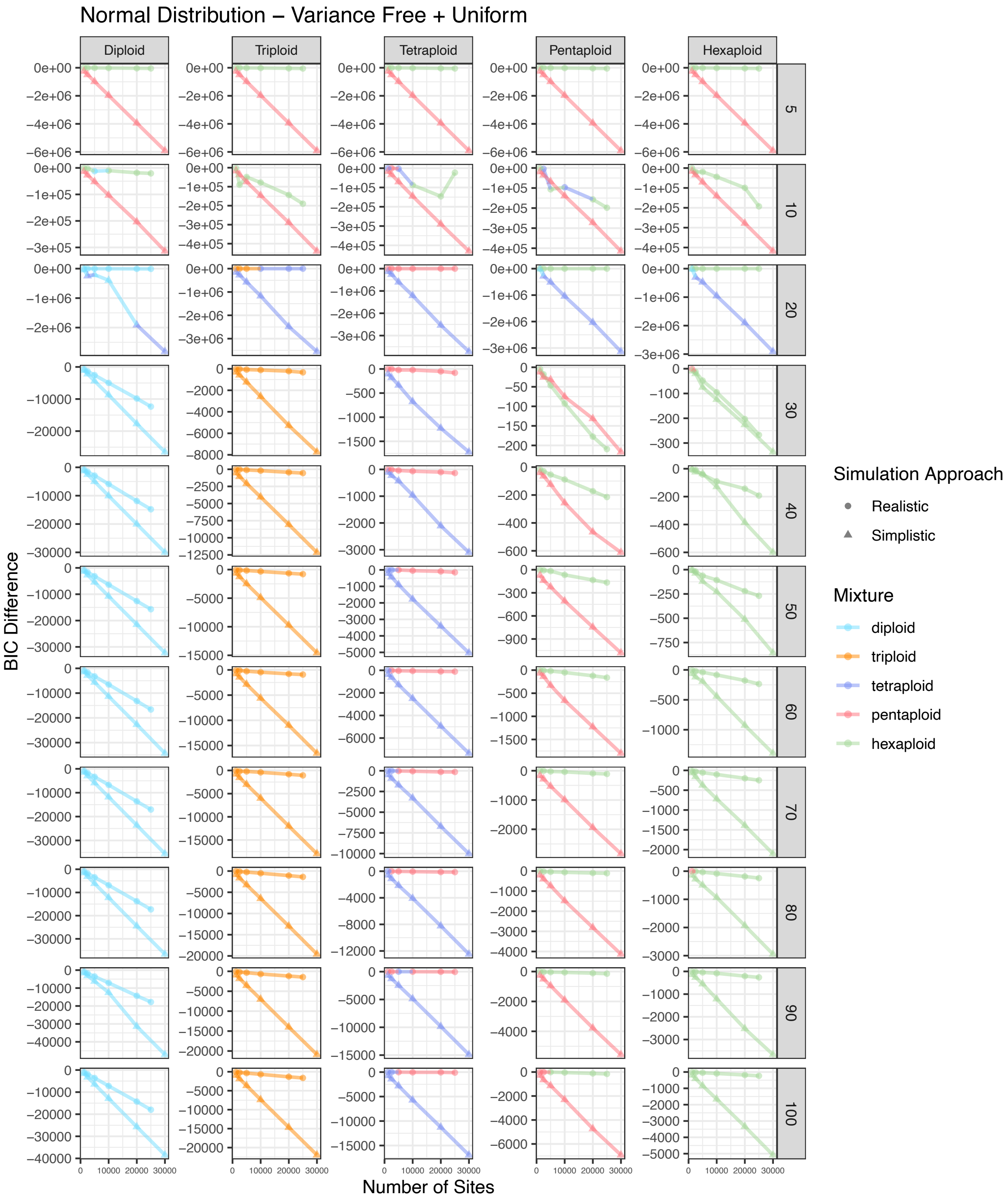


Figure S7. BIC score difference between the best and second best model for simulated diploid, triploid, tetraploid, pentaploid, and hexaploid samples across different numbers of sites for eleven different coverage amounts. The color of each point represents the best model. The shape of each point represents the approach used to simulate that sample. This represents a normal distribution with variance free with a uniform mixture.

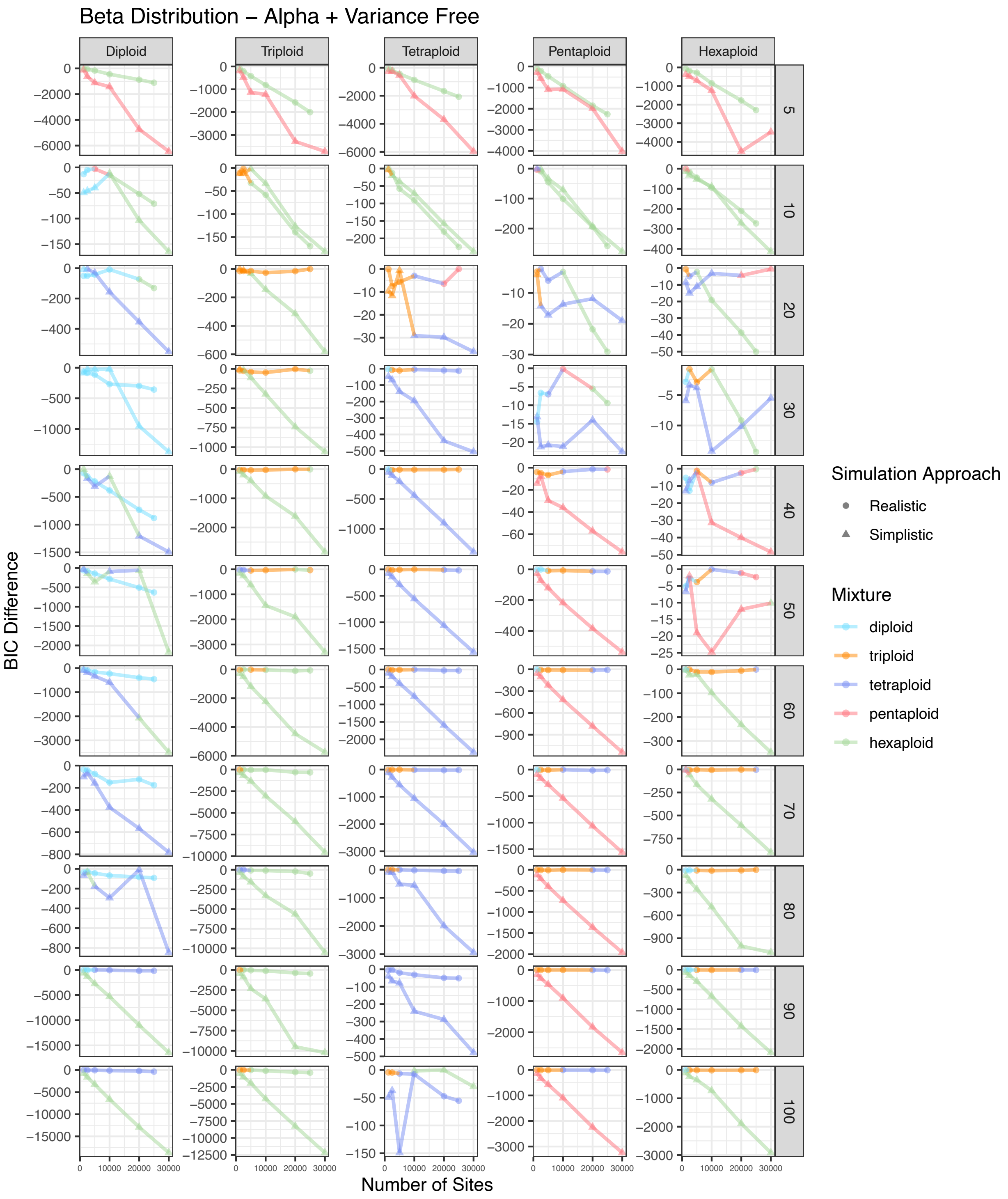


Figure S8. BIC score difference between the best and second best model for simulated diploid, triploid, tetraploid, pentaploid, and hexaploid samples across different numbers of sites for eleven different coverage amounts. The color of each point represents the best model. The shape of each point represents the approach used to simulate that sample. This represents a beta distribution with alpha and variance free.

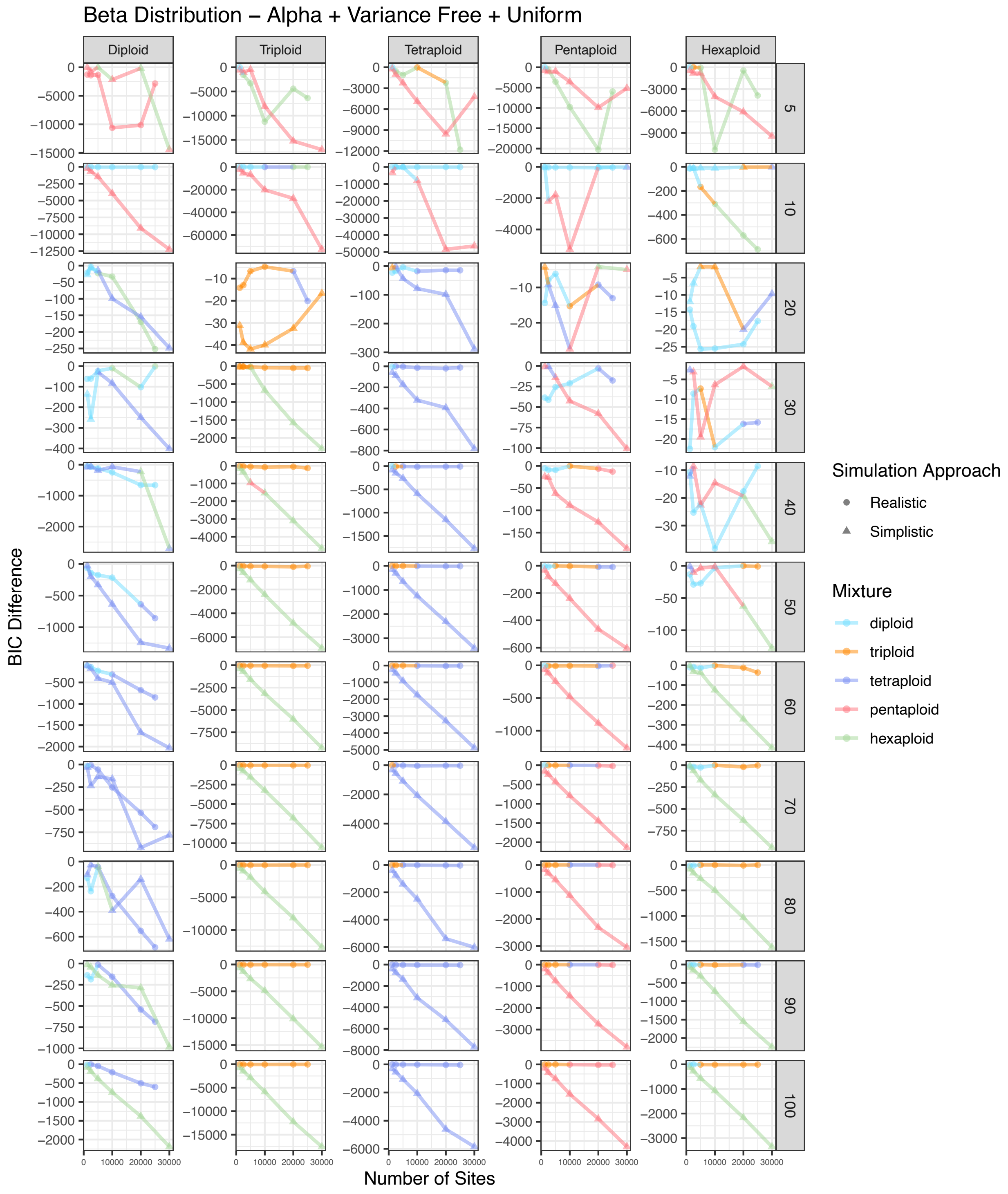


Figure S9. BIC score difference between the best and second best model for simulated diploid, triploid, tetraploid, pentaploid, and hexaploid samples across different numbers of sites for eleven different coverage amounts. The color of each point represents the best model. The shape of each point represents the approach used to simulate that sample. This represents a beta distribution with alpha and variance free with a uniform mixture.

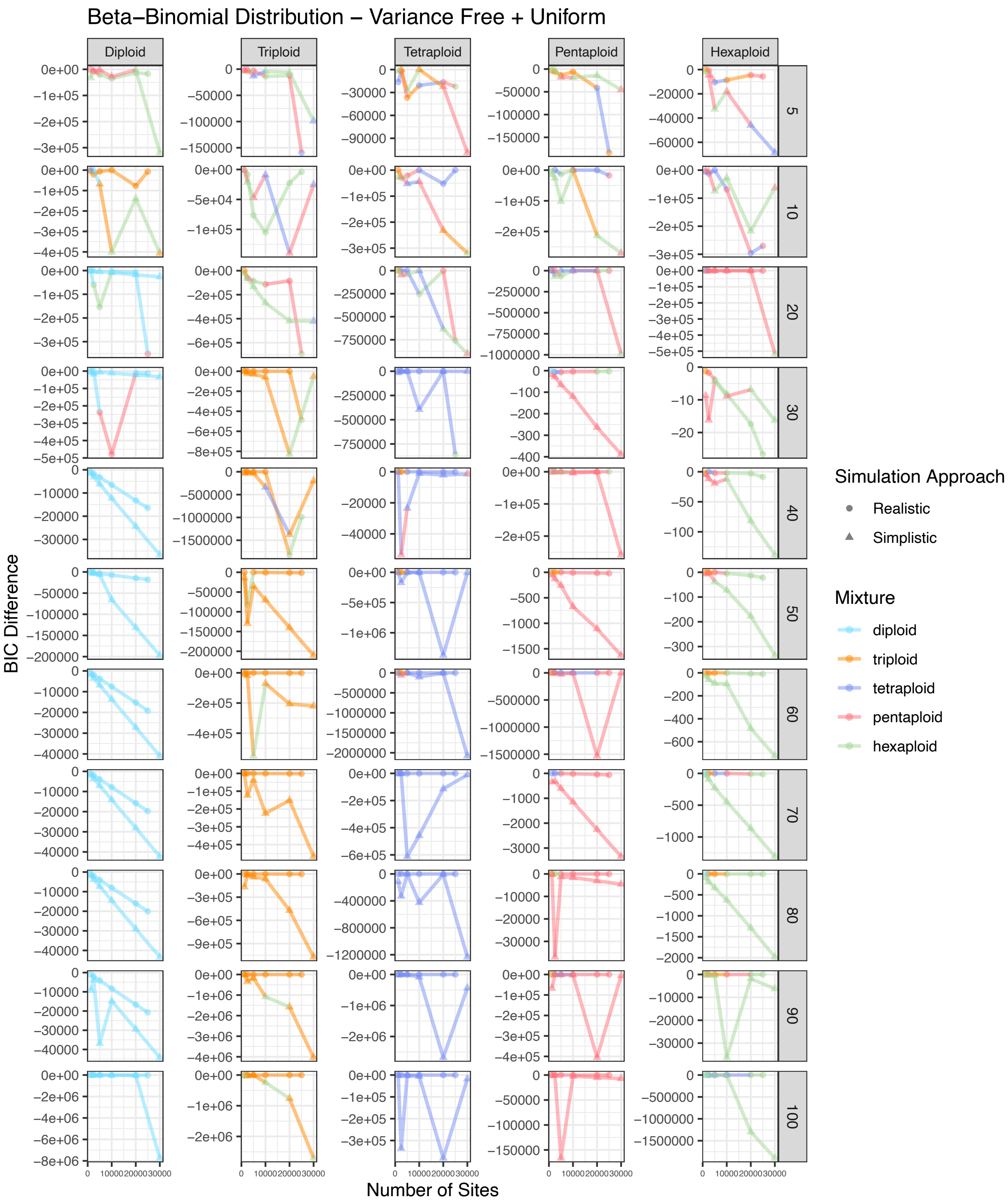

Beta–Binomial Distribution – Variance Free + Uniform
Diploid
Triploid
Tetraploid
Pentaploid
Hexaploid
5
10
20
30
40
50
60
70
80
90
100
BIC Difference
Number of Sites
Simulation Approach
Realistic
Simplistic
Mixture
diploid
triploid
tetraploid
pentaploid
hexaploid

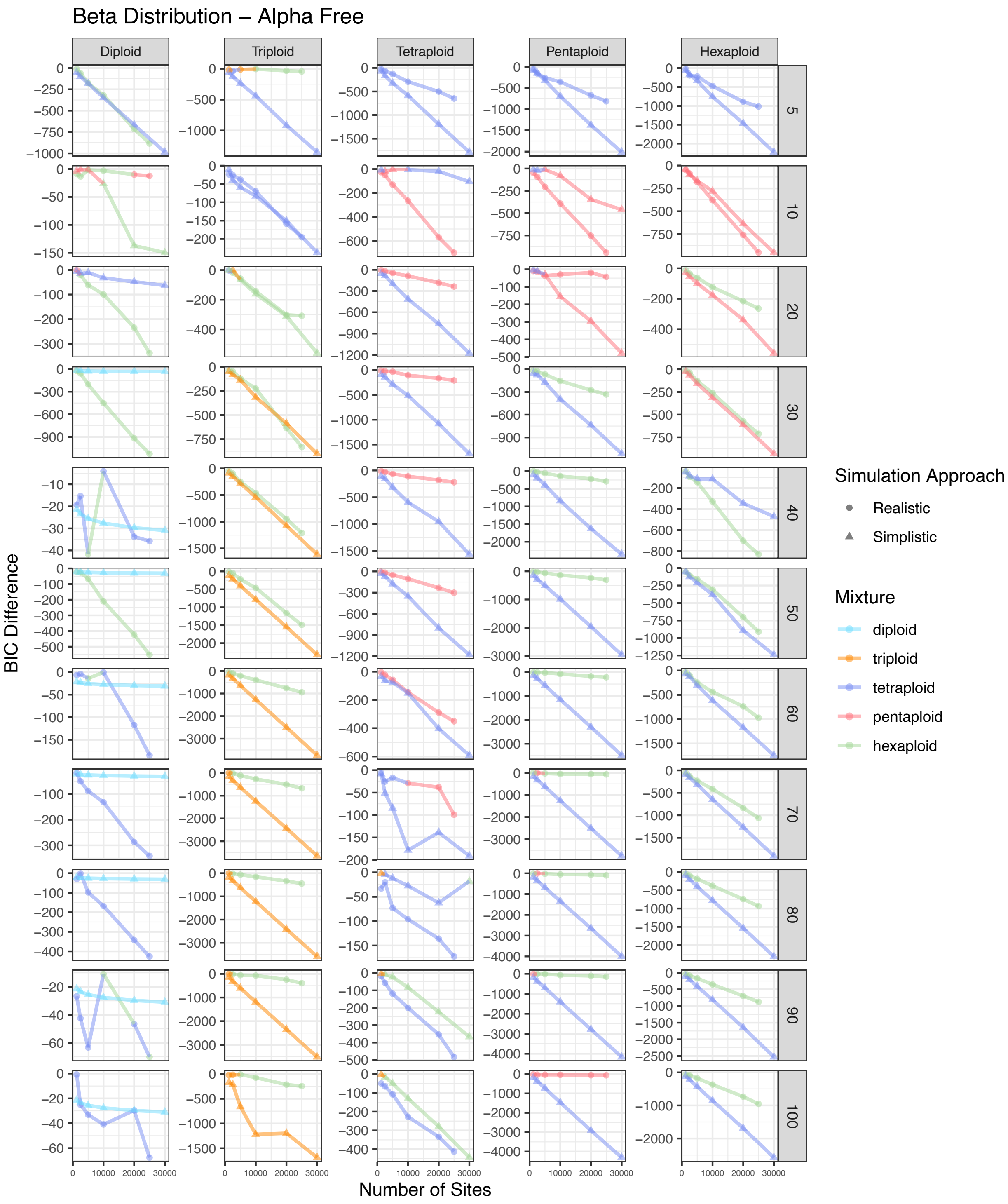


Figure S10. BIC score difference between the best and second best model for simulated diploid, triploid, tetraploid, pentaploid, and hexaploid samples across different numbers of sites for eleven different coverage amounts. The color of each point represents the best model. The shape of each point represents the approach used to simulate that sample. This represents a beta distribution with alpha free.

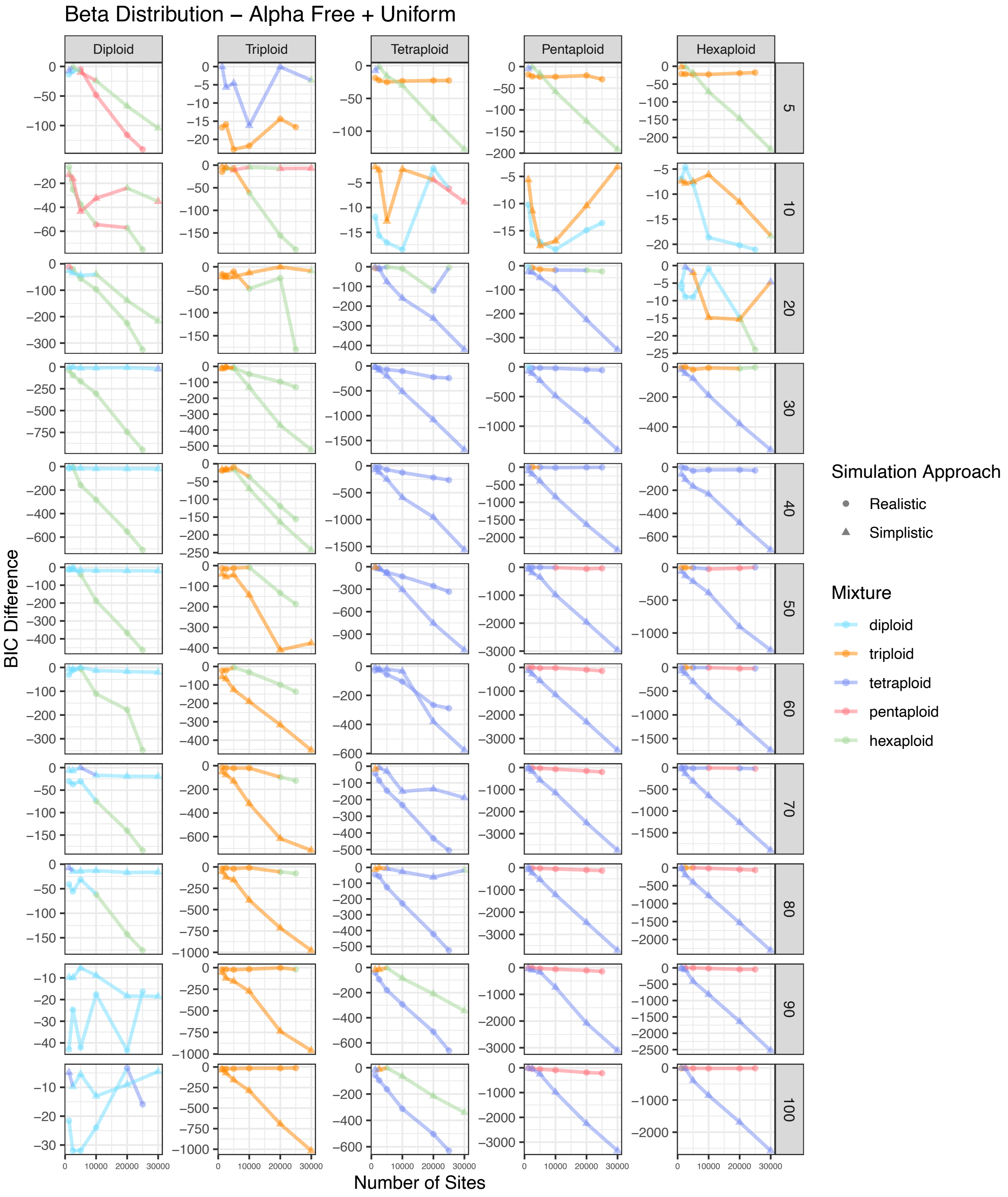


Figure S11. BIC score difference between the best and second best model for simulated diploid, triploid, tetraploid, pentaploid, and hexaploid samples across different numbers of sites for eleven different coverage amounts. The color of each point represents the best model. The shape of each point represents the approach used to simulate that sample. This represents a beta distribution with alpha free with a uniform mixture.

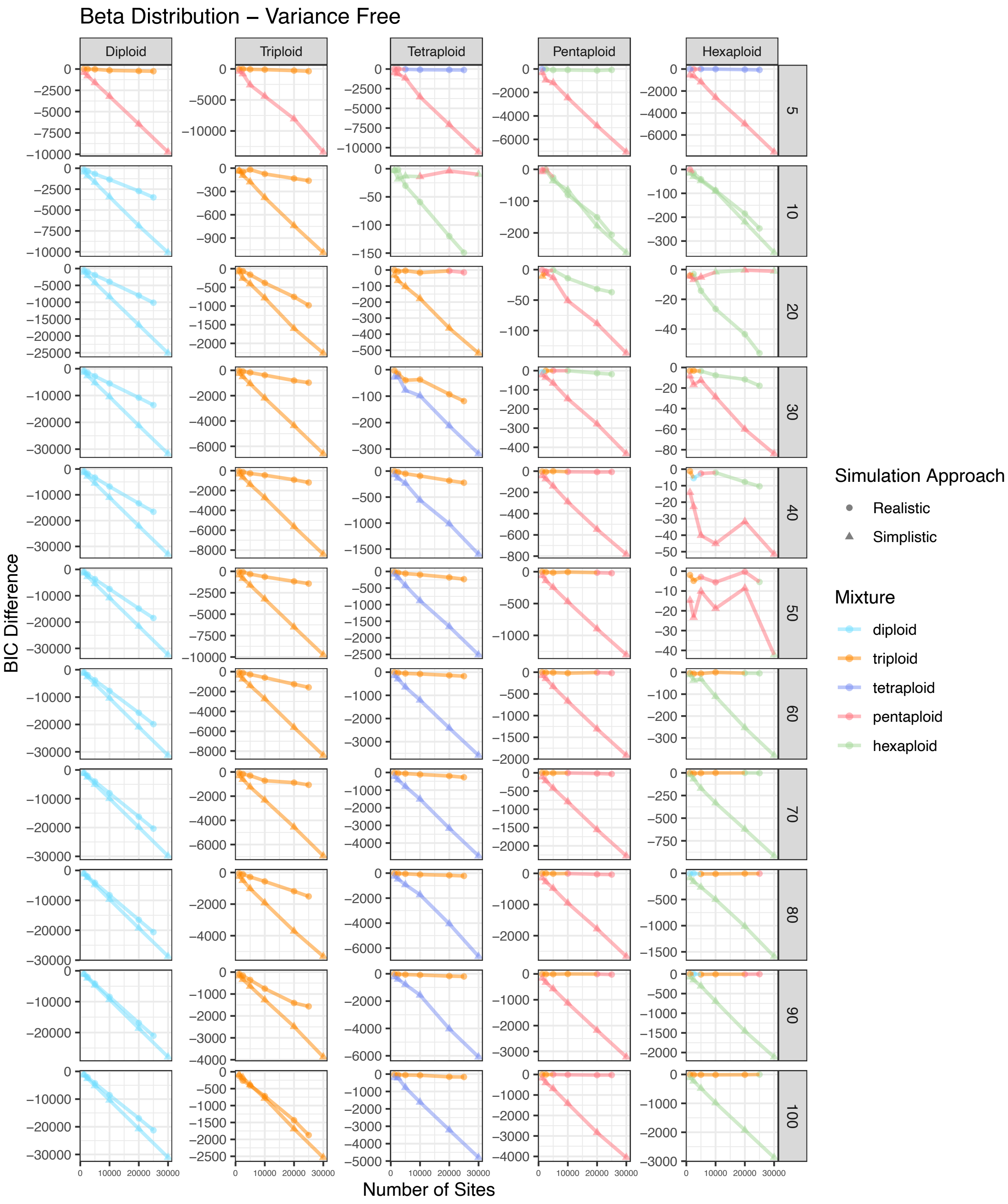


Figure S12. BIC score difference between the best and second best model for simulated diploid, triploid, tetraploid, pentaploid, and hexaploid samples across different numbers of sites for eleven different coverage amounts. The color of each point represents the best model. The shape of each point represents the approach used to simulate that sample. This represents a beta distribution with variance free.

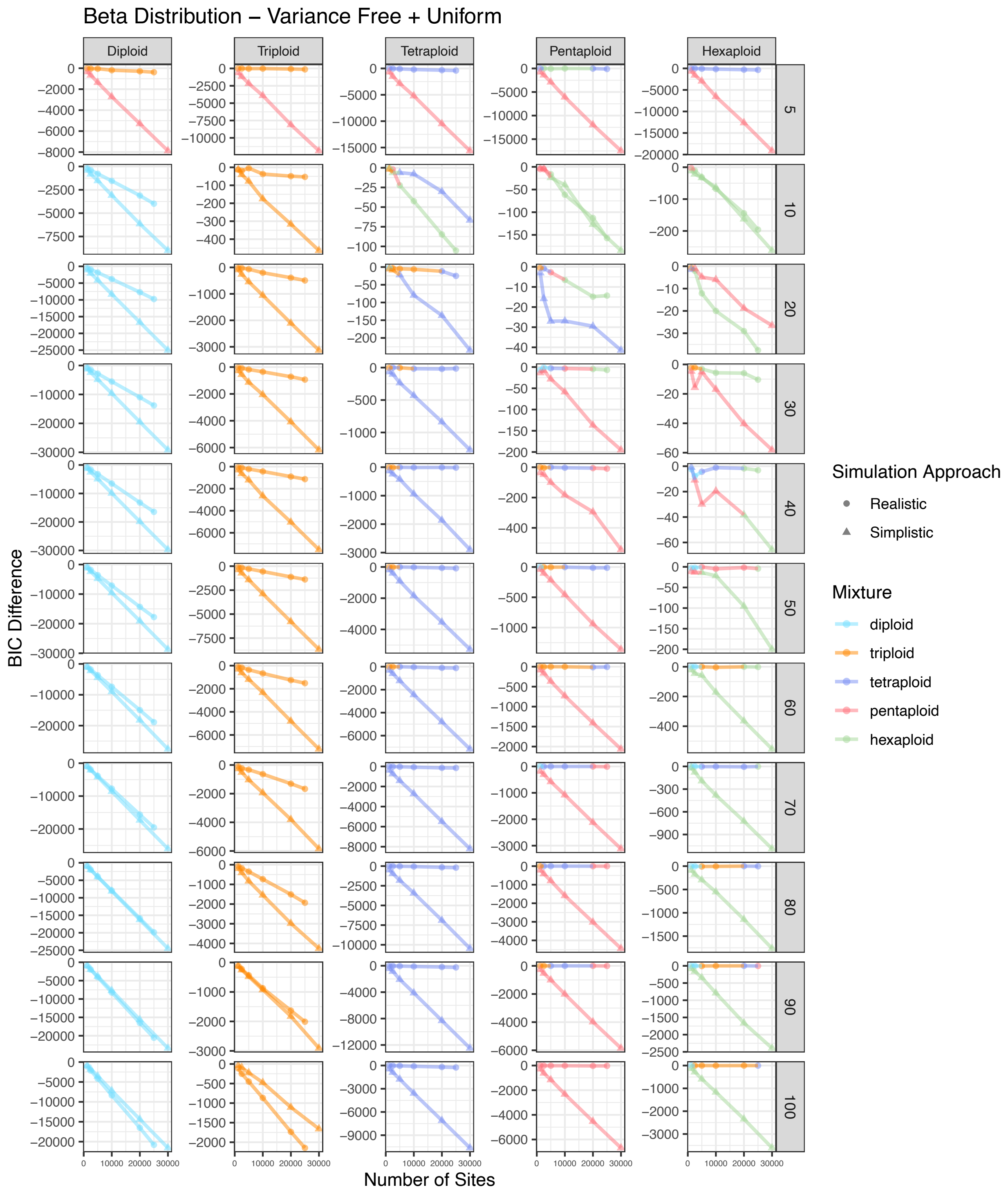


Figure S13. BIC score difference between the best and second best model for simulated diploid, triploid, tetraploid, pentaploid, and hexaploid samples across different numbers of sites for eleven different coverage amounts. The color of each point represents the best model. The shape of each point represents the approach used to simulate that sample. This represents a beta distribution with variance free with a uniform mixture.

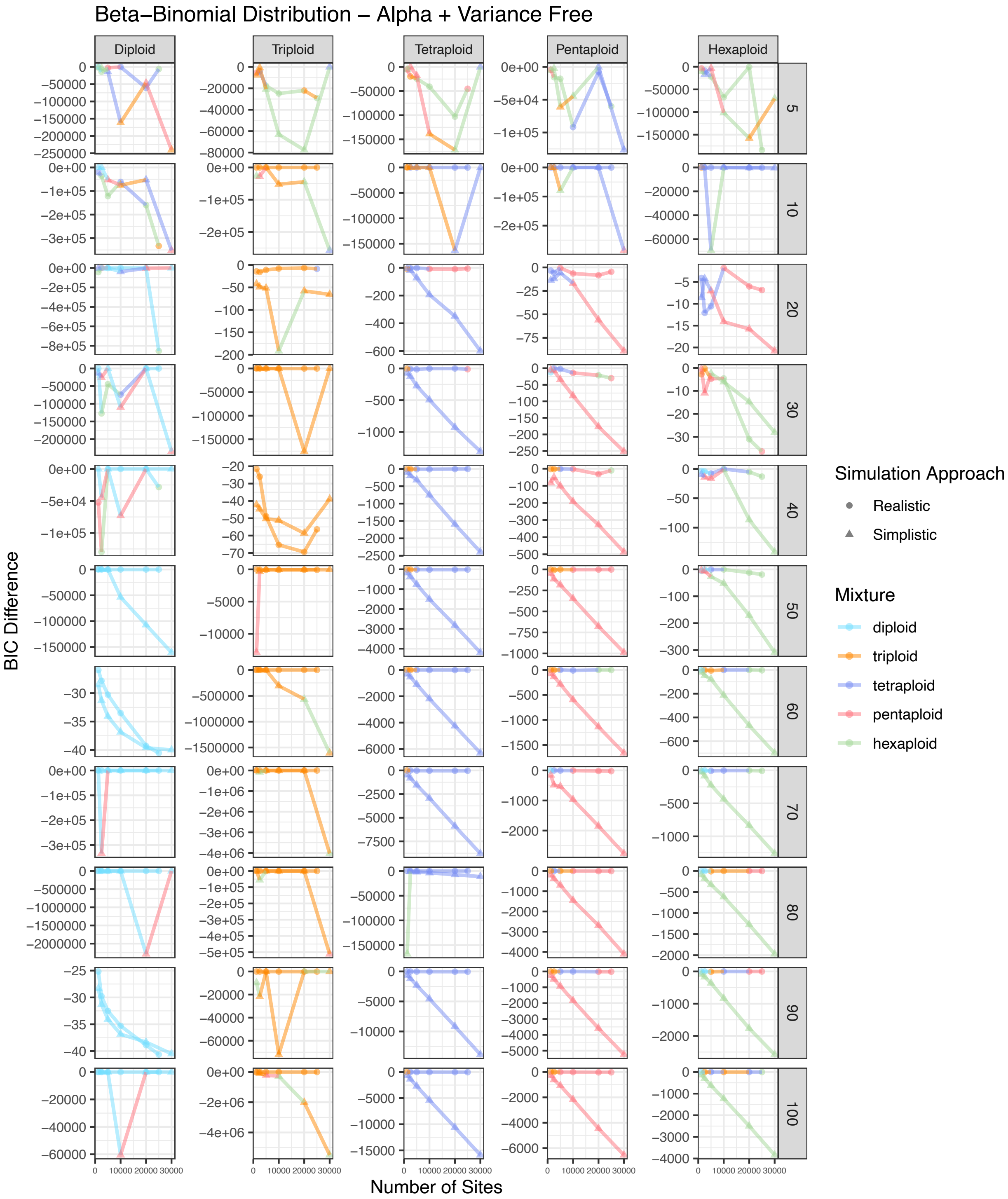


Figure S14. BIC score difference between the best and second best model for simulated diploid, triploid, tetraploid, pentaploid, and hexaploid samples across different numbers of sites for eleven different coverage amounts. The color of each point represents the best model. The shape of each point represents the approach used to simulate that sample. This represents a beta-binomial distribution with alpha and variance free.

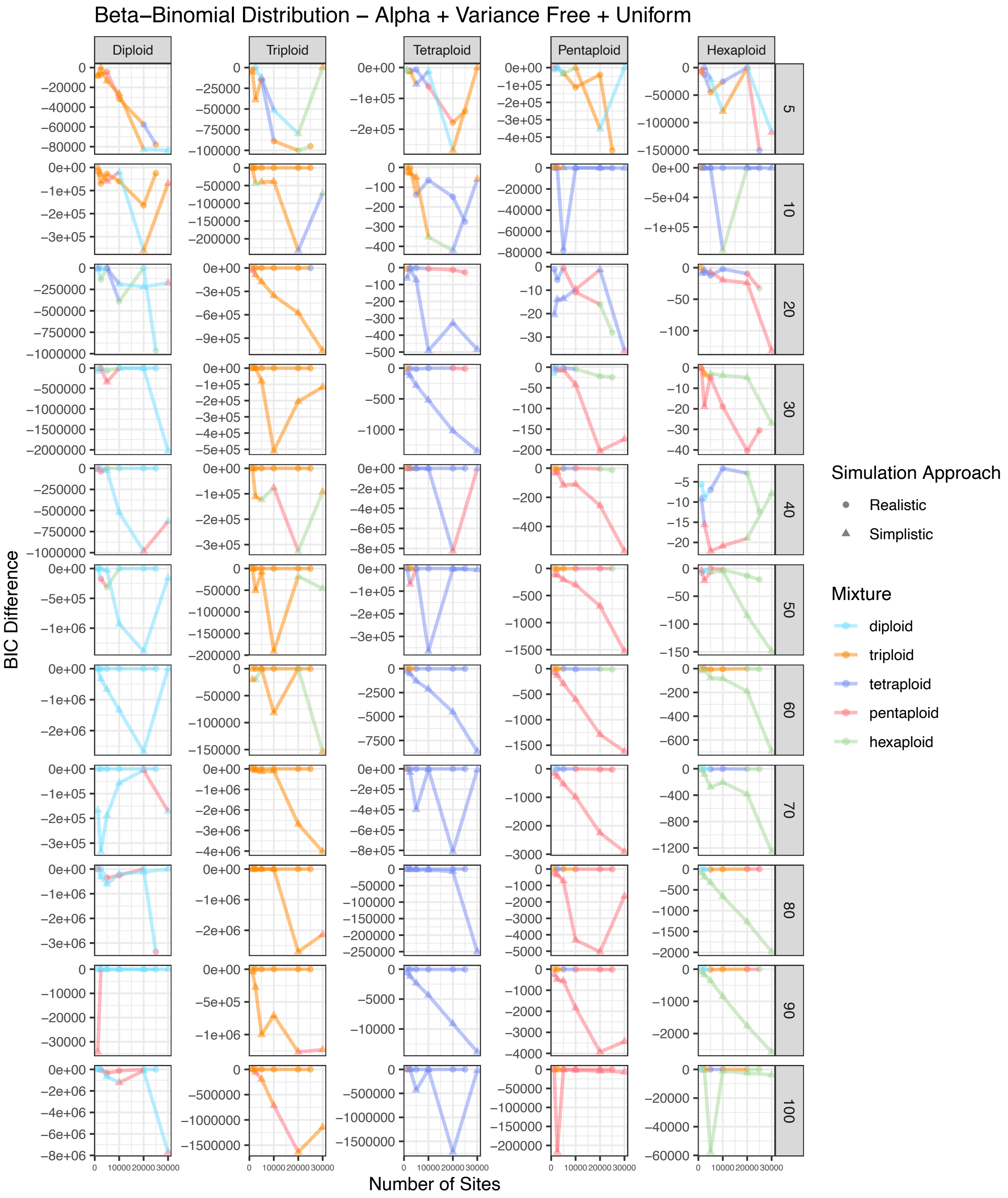


Figure S15. BIC score difference between the best and second best model for simulated diploid, triploid, tetraploid, pentaploid, and hexaploid samples across different numbers of sites for eleven different coverage amounts. The color of each point represents the best model. The shape of each point represents the approach used to simulate that sample. This represents a beta-binomial distribution with alpha and variance free with a uniform mixture.

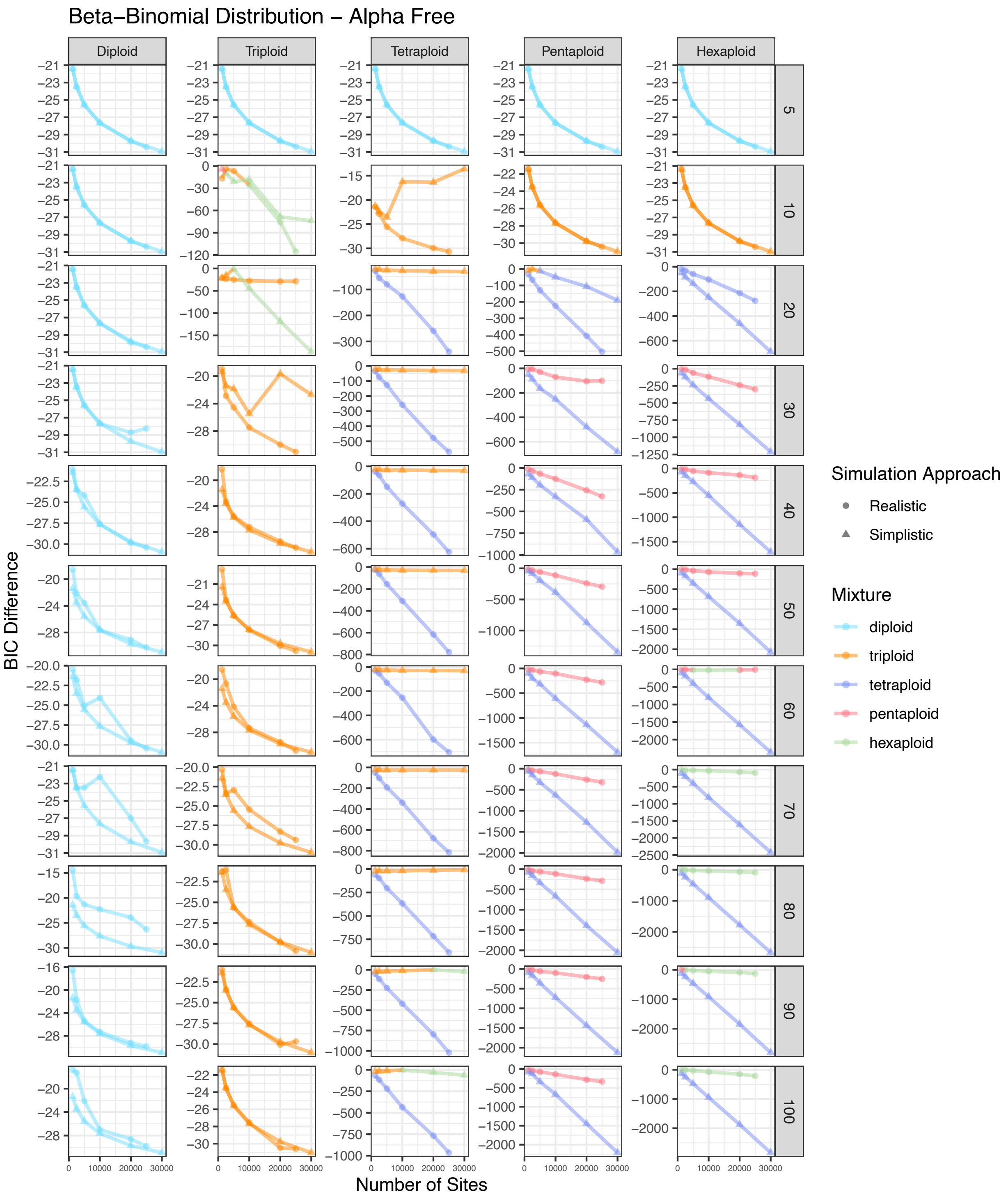


Figure S16. BIC score difference between the best and second best model for simulated diploid, triploid, tetraploid, pentaploid, and hexaploid samples across different numbers of sites for eleven different coverage amounts. The color of each point represents the best model. The shape of each point represents the approach used to simulate that sample. This represents a beta-binomial distribution with alpha free.

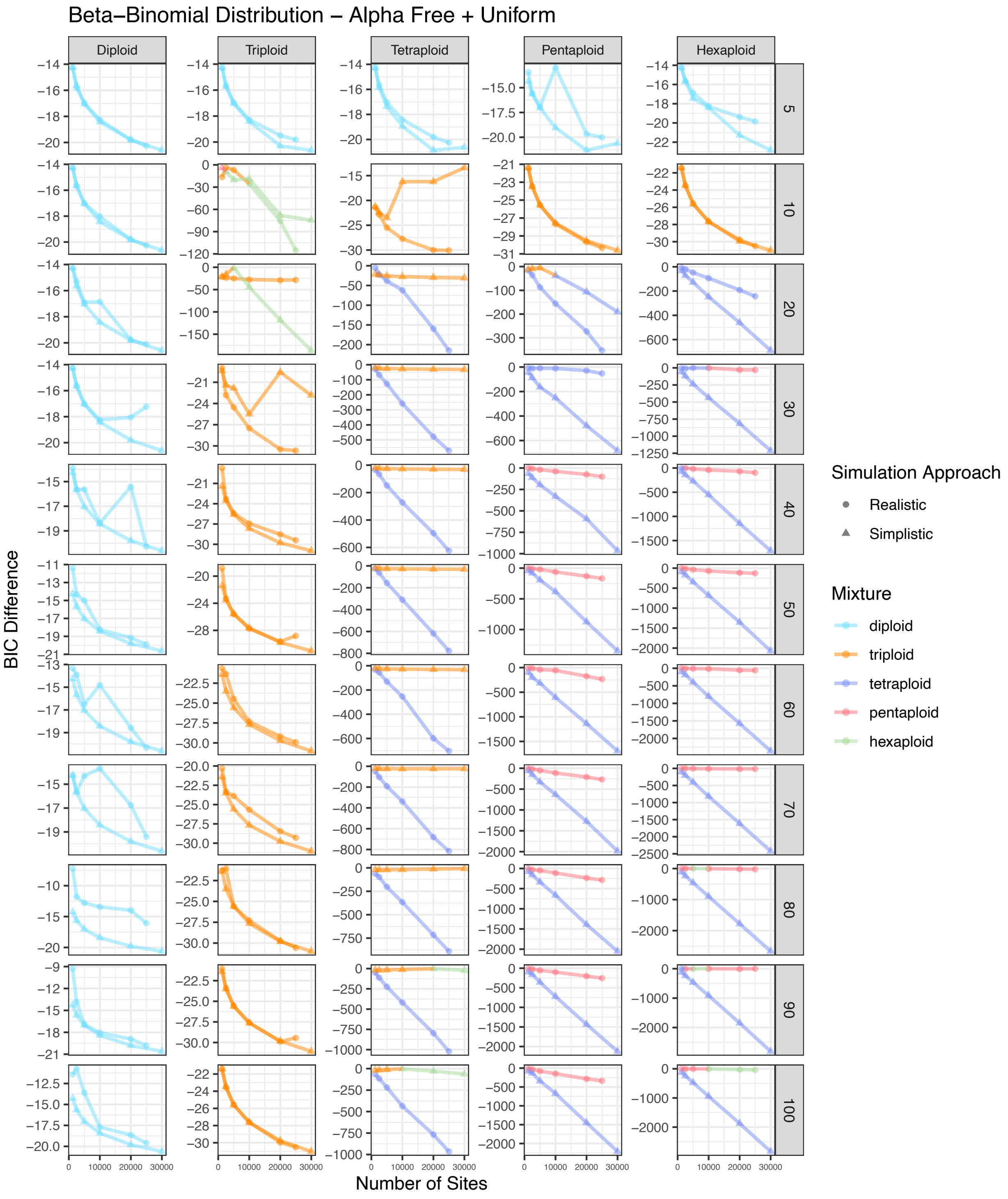


Figure S17. BIC score difference between the best and second best model for simulated diploid, triploid, tetraploid, pentaploid, and hexaploid samples across different numbers of sites for eleven different coverage amounts. The color of each point represents the best model. The shape of each point represents the approach used to simulate that sample. This represents a beta-binomial distribution with alpha free with a uniform mixture.

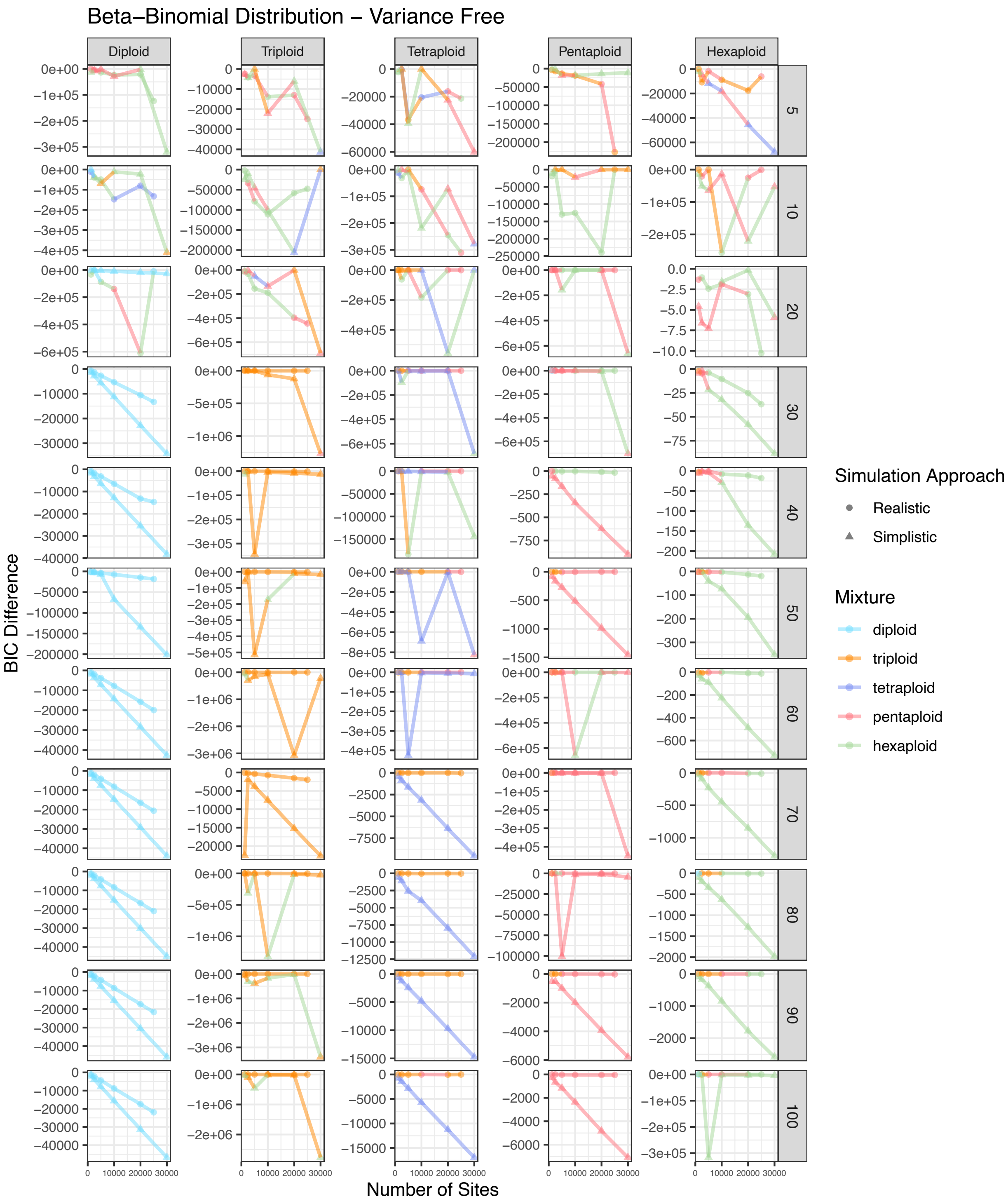


Figure S18. BIC score difference between the best and second best model for simulated diploid, triploid, tetraploid, pentaploid, and hexaploid samples across different numbers of sites for eleven different coverage amounts. The color of each point represents the best model. The shape of each point represents the approach used to simulate that sample. This represents a beta-binomial distribution with variance free.

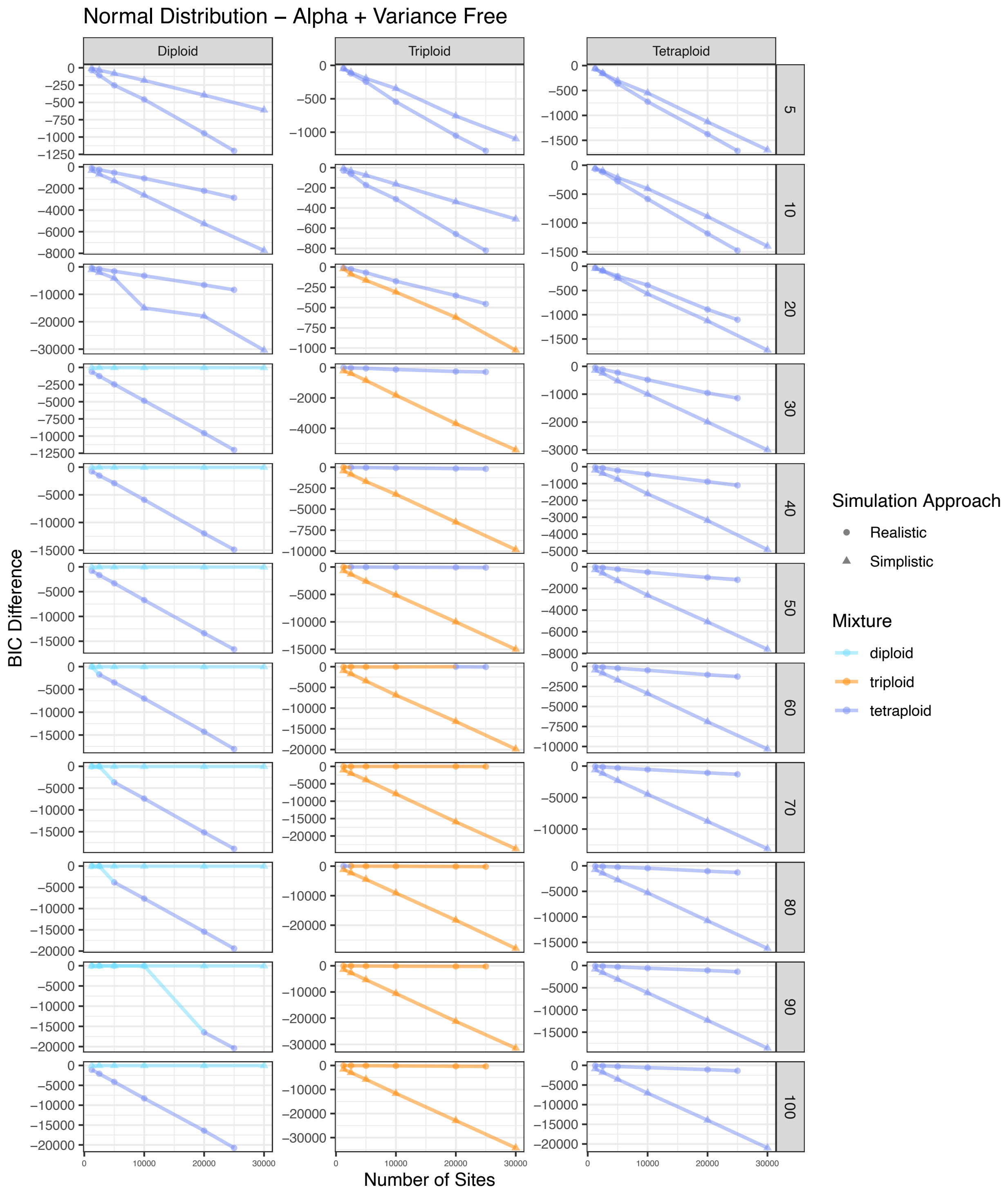


Figure S20. BIC score difference between the best and second best model for simulated diploid, triploid, and tetraploid samples across different numbers of sites for eleven different coverage amounts. The color of each point represents the best model. The shape of each point represents the approach used to simulate that sample. This represents a normal distribution with alpha and variance free.

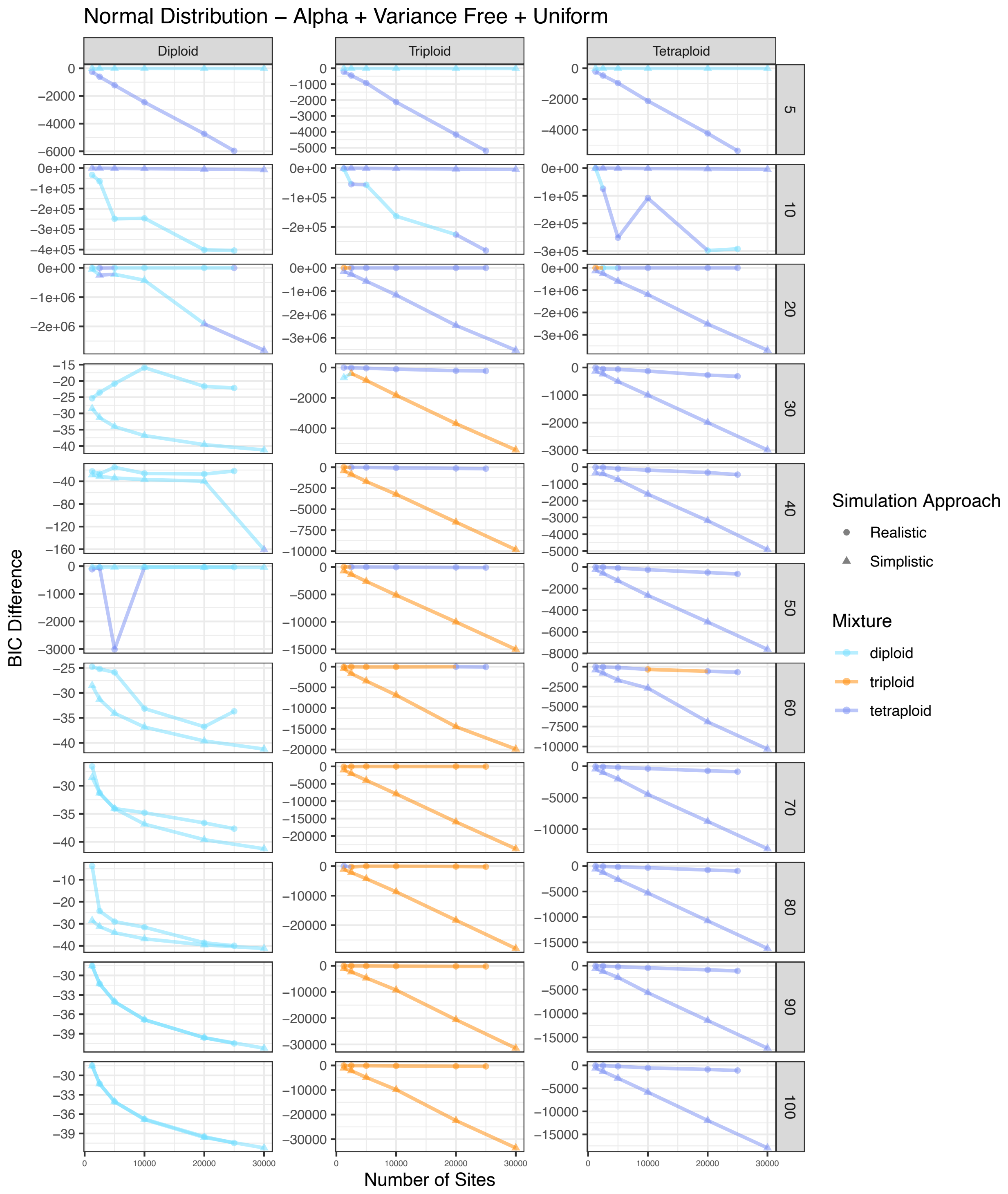


Figure S21. BIC score difference between the best and second best model for simulated diploid, triploid, and tetraploid samples across different numbers of sites for eleven different coverage amounts. The color of each point represents the best model. The shape of each point represents the approach used to simulate that sample. This represents a normal distribution with alpha and variance free with a uniform mixture.

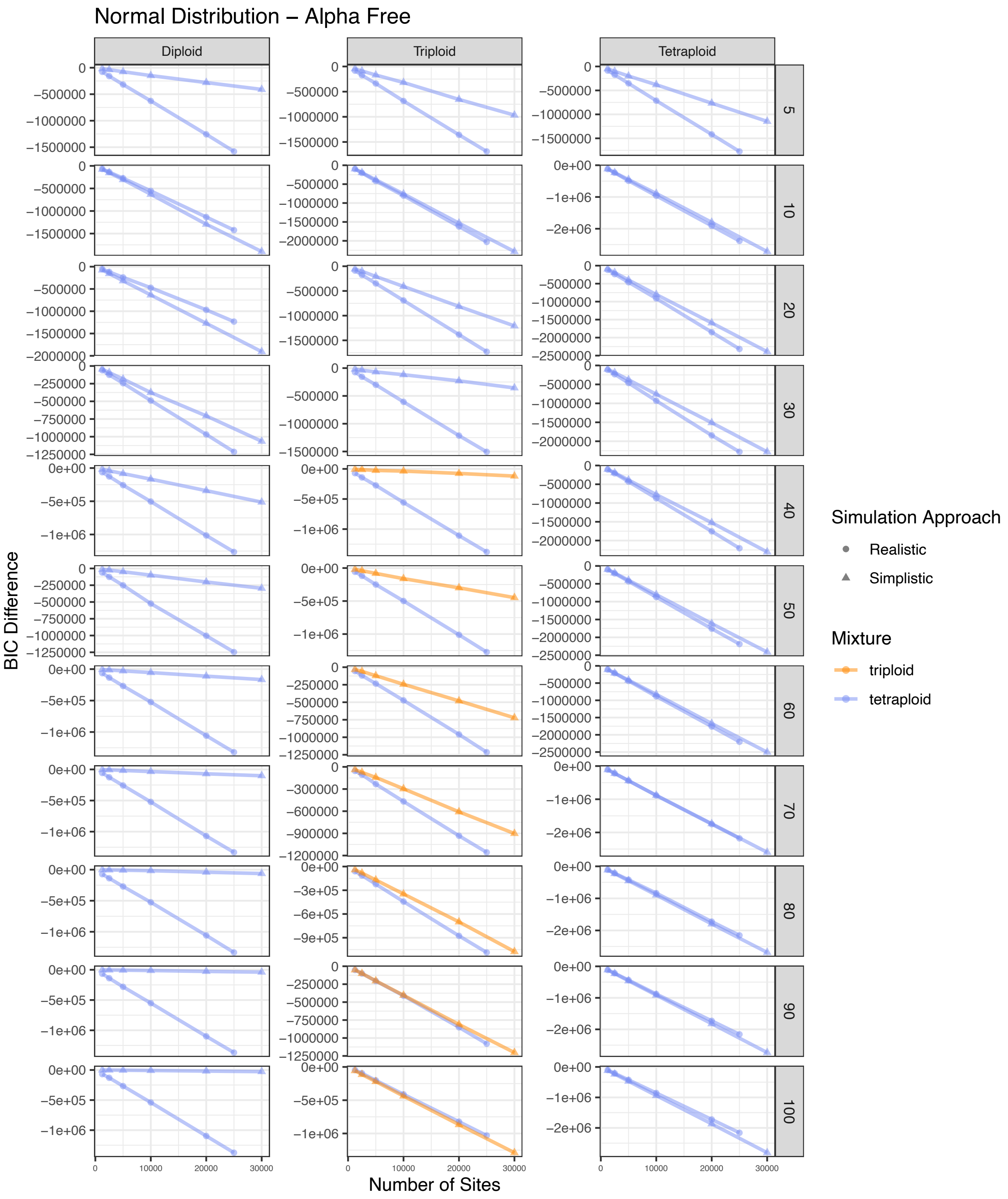


Figure S22. BIC score difference between the best and second best model for simulated diploid, triploid, and tetraploid samples across different numbers of sites for eleven different coverage amounts. The color of each point represents the best model. The shape of each point represents the approach used to simulate that sample. This represents a normal distribution with alpha free.

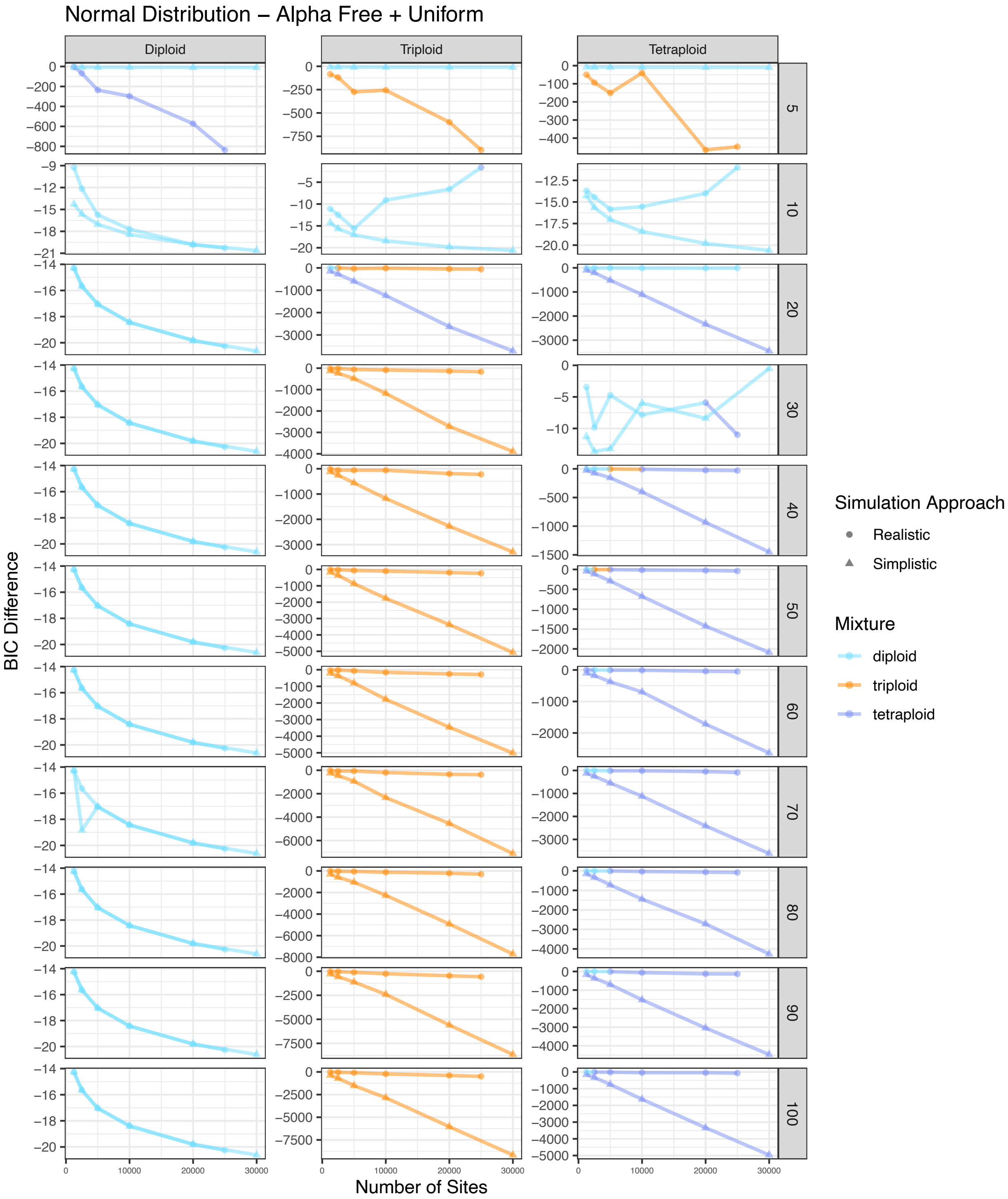


Figure S23. BIC score difference between the best and second best model for simulated diploid, triploid, and tetraploid samples across different numbers of sites for eleven different coverage amounts. The color of each point represents the best model. The shape of each point represents the approach used to simulate that sample. This represents a normal distribution with alpha free with a uniform mixture.

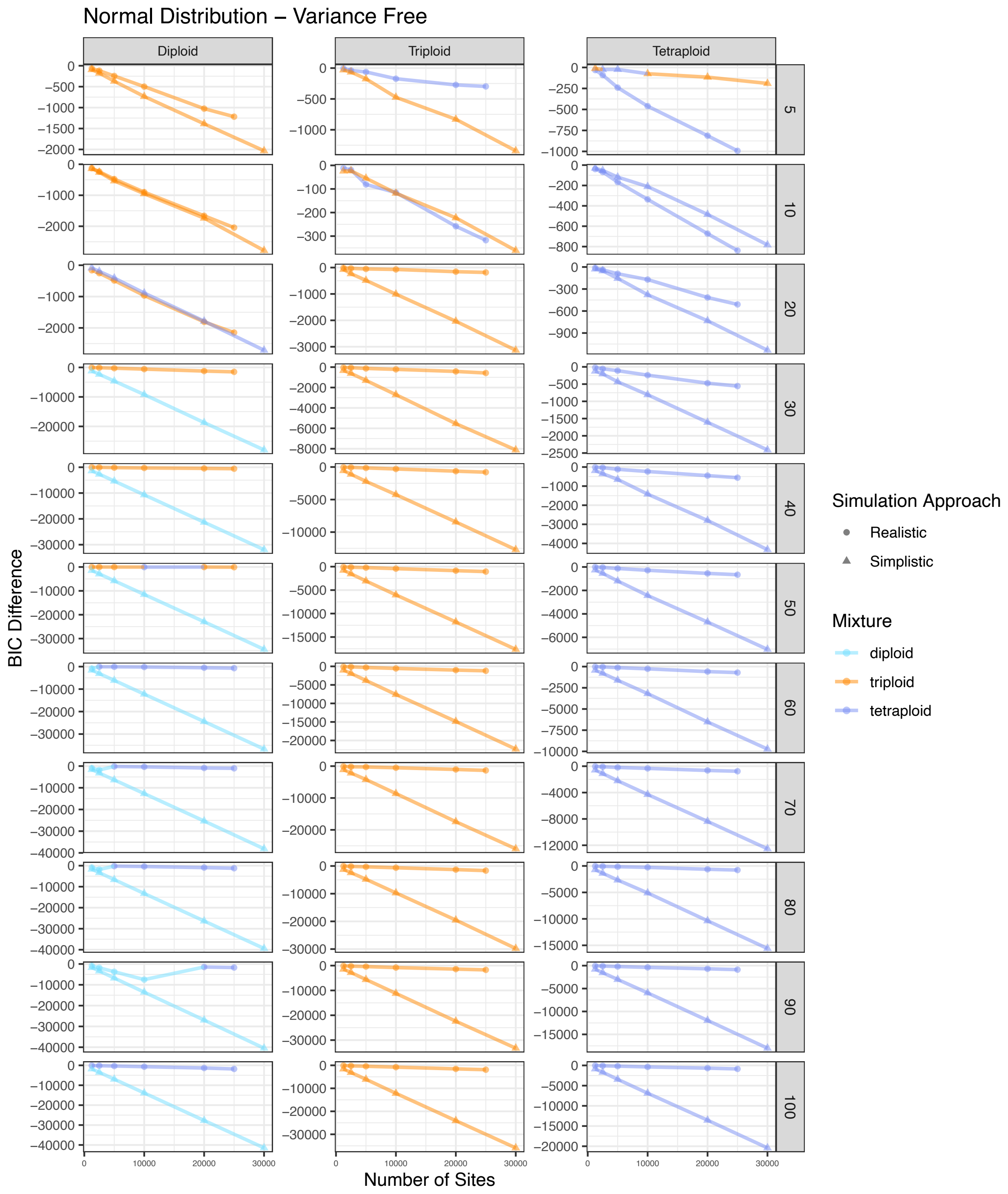


Figure S24. BIC score difference between the best and second best model for simulated diploid, triploid, and tetraploid samples across different numbers of sites for eleven different coverage amounts. The color of each point represents the best model. The shape of each point represents the approach used to simulate that sample. This represents a normal distribution with variance free.

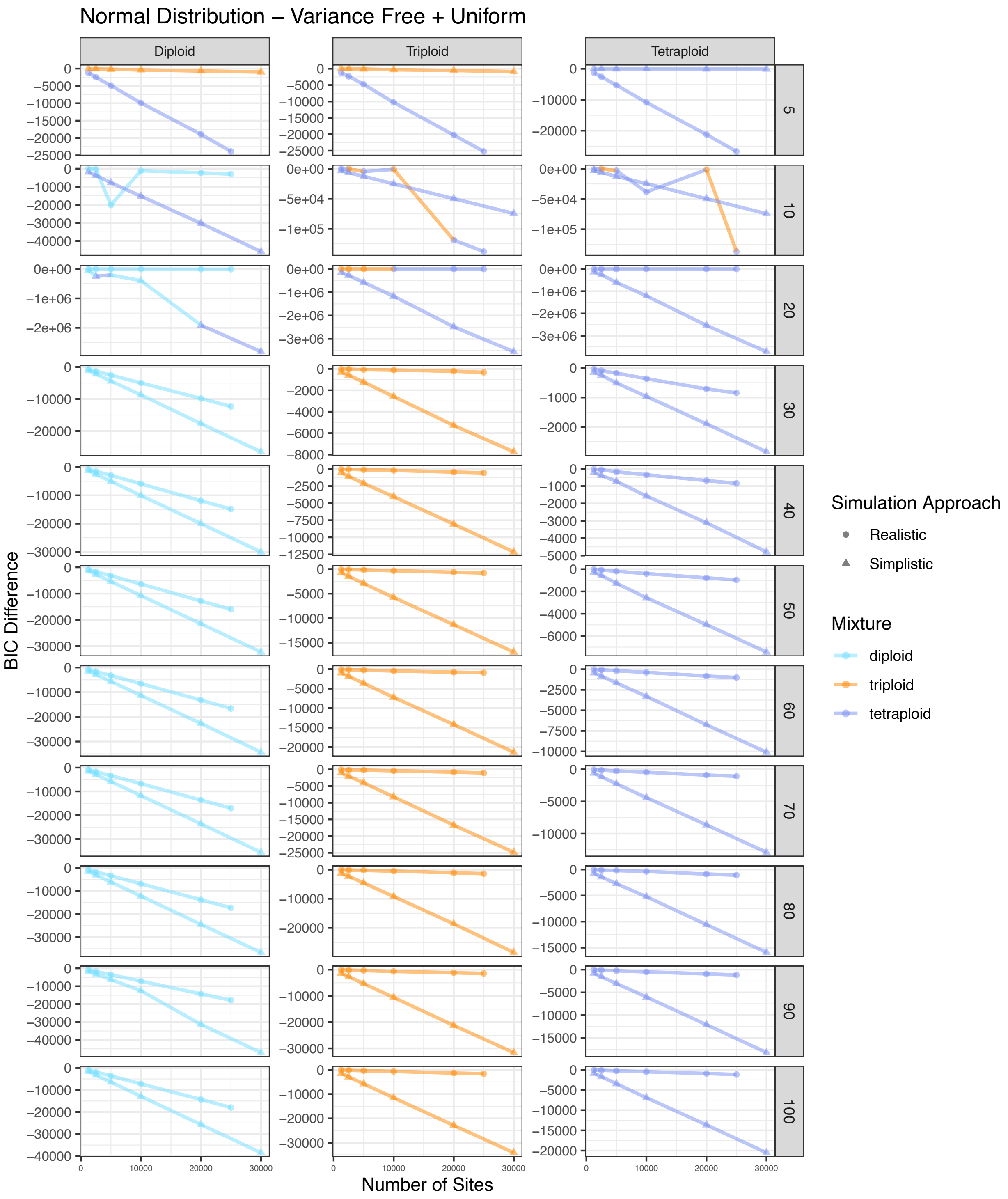


Figure S25. BIC score difference between the best and second best model for simulated diploid, triploid, and tetraploid samples across different numbers of sites for eleven different coverage amounts. The color of each point represents the best model. The shape of each point represents the approach used to simulate that sample. This represents a normal distribution with variance free with a uniform mixture.

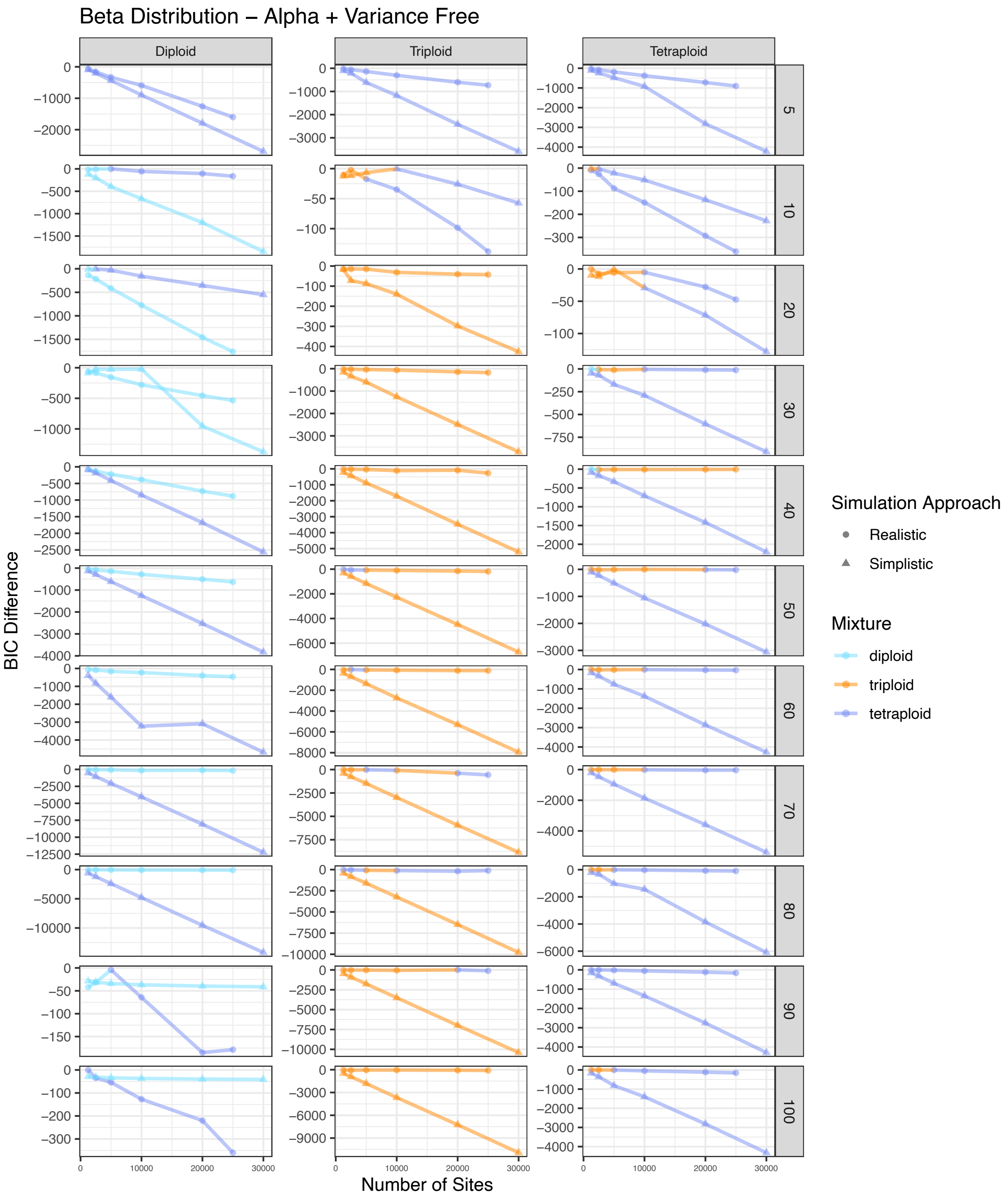


Figure S26. BIC score difference between the best and second best model for simulated diploid, triploid, and tetraploid samples across different numbers of sites for eleven different coverage amounts. The color of each point represents the best model. The shape of each point represents the approach used to simulate that sample. This represents a beta distribution with alpha and variance free.

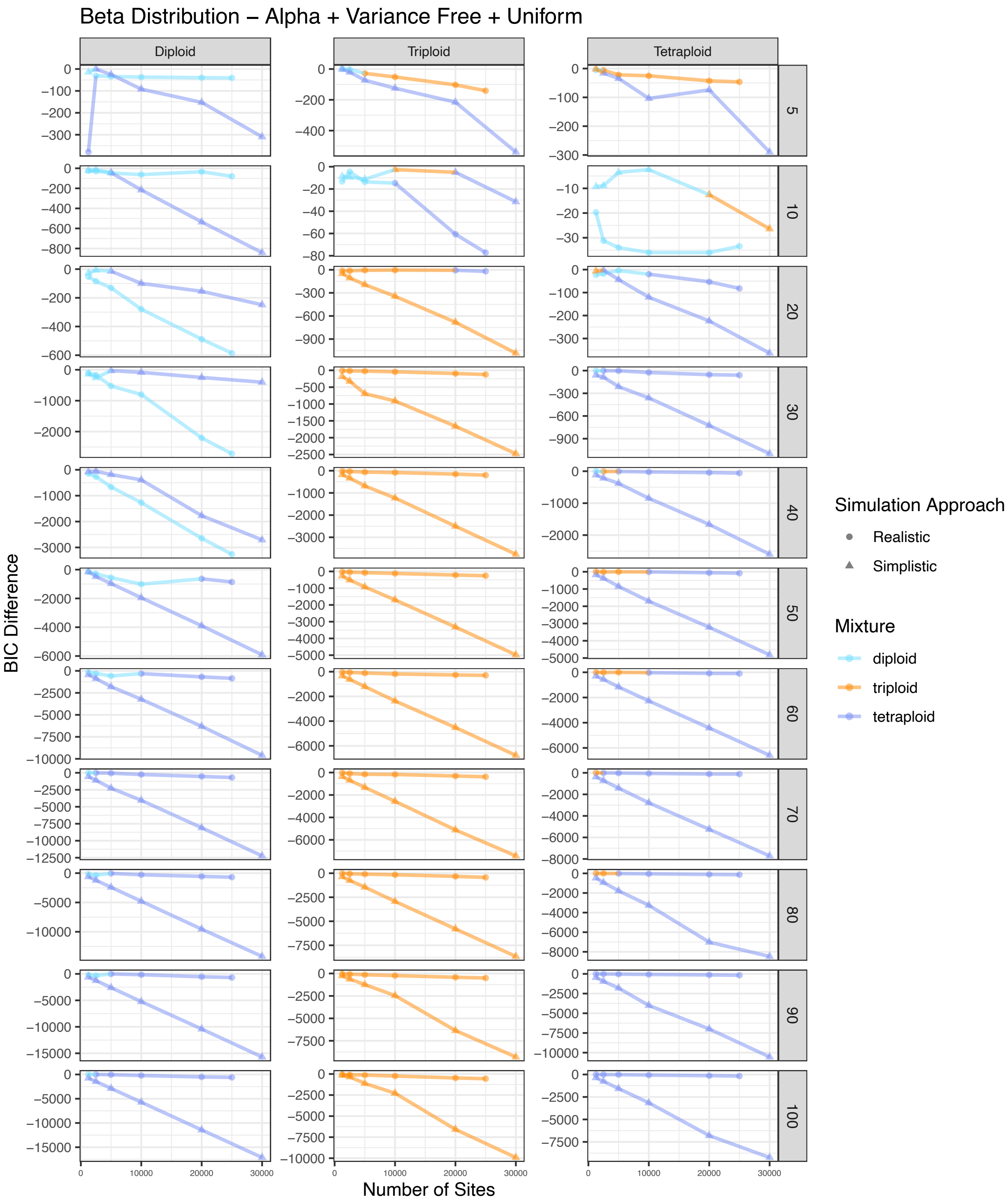


Figure S27. BIC score difference between the best and second best model for simulated diploid, triploid, and tetraploid samples across different numbers of sites for eleven different coverage amounts. The color of each point represents the best model. The shape of each point represents the approach used to simulate that sample. This represents a beta distribution with alpha and variance free with a uniform mixture.

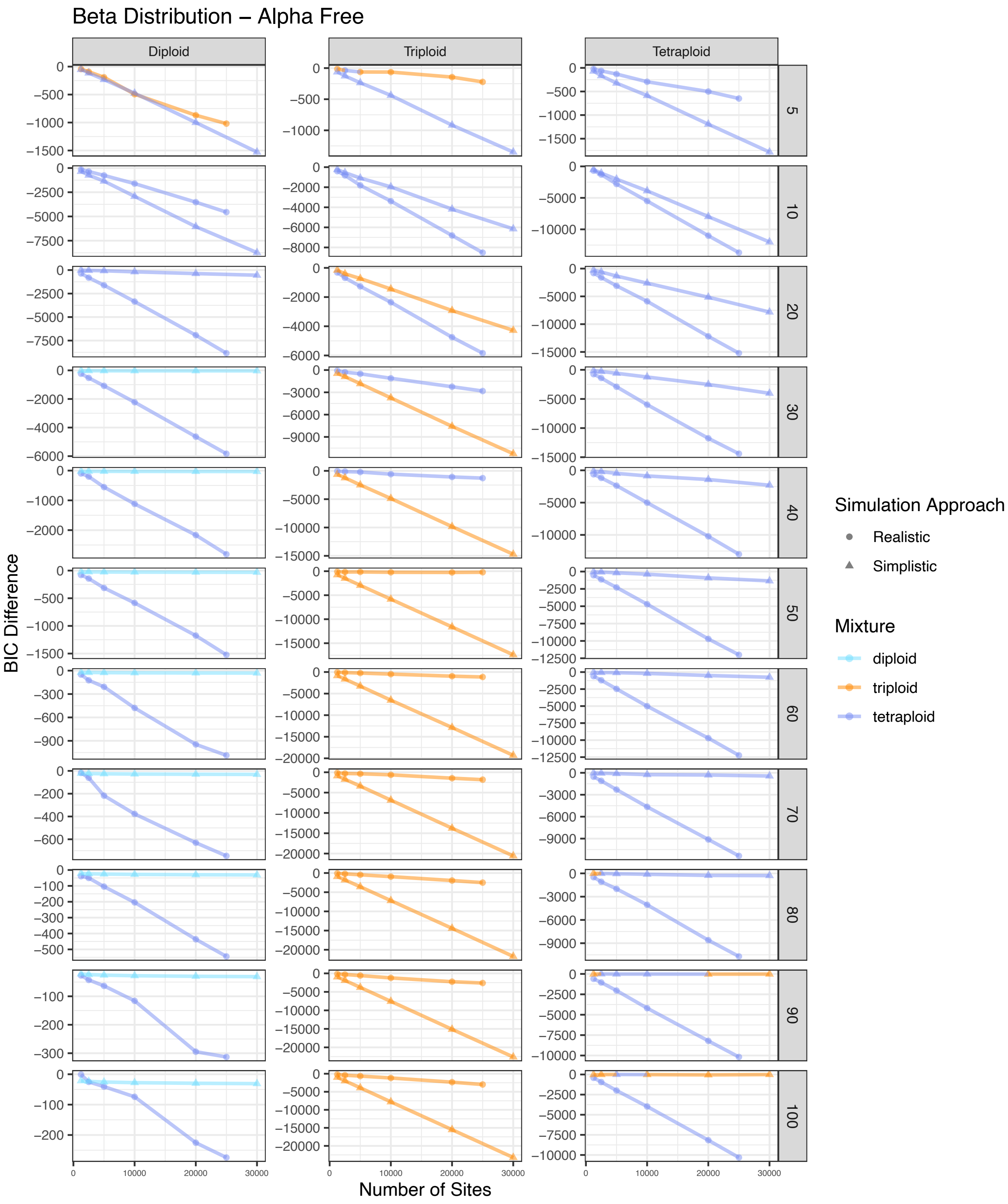


Figure S28. BIC score difference between the best and second best model for simulated diploid, triploid, and tetraploid samples across different numbers of sites for eleven different coverage amounts. The color of each point represents the best model. The shape of each point represents the approach used to simulate that sample. This represents a beta distribution with alpha free.

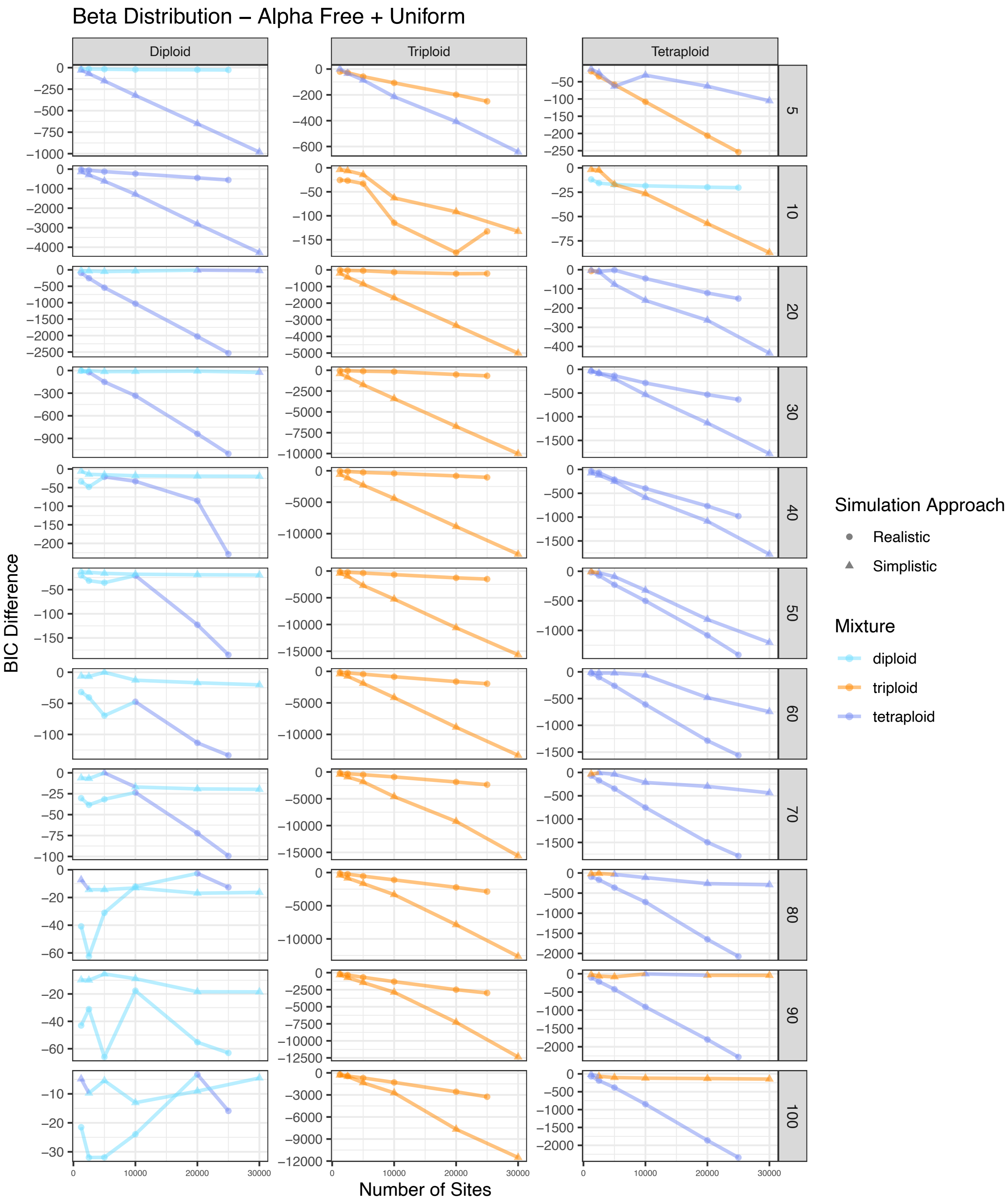


Figure S29. BIC score difference between the best and second best model for simulated diploid, triploid, and tetraploid samples across different numbers of sites for eleven different coverage amounts. The color of each point represents the best model. The shape of each point represents the approach used to simulate that sample. This represents a beta distribution with alpha free with a uniform mixture.

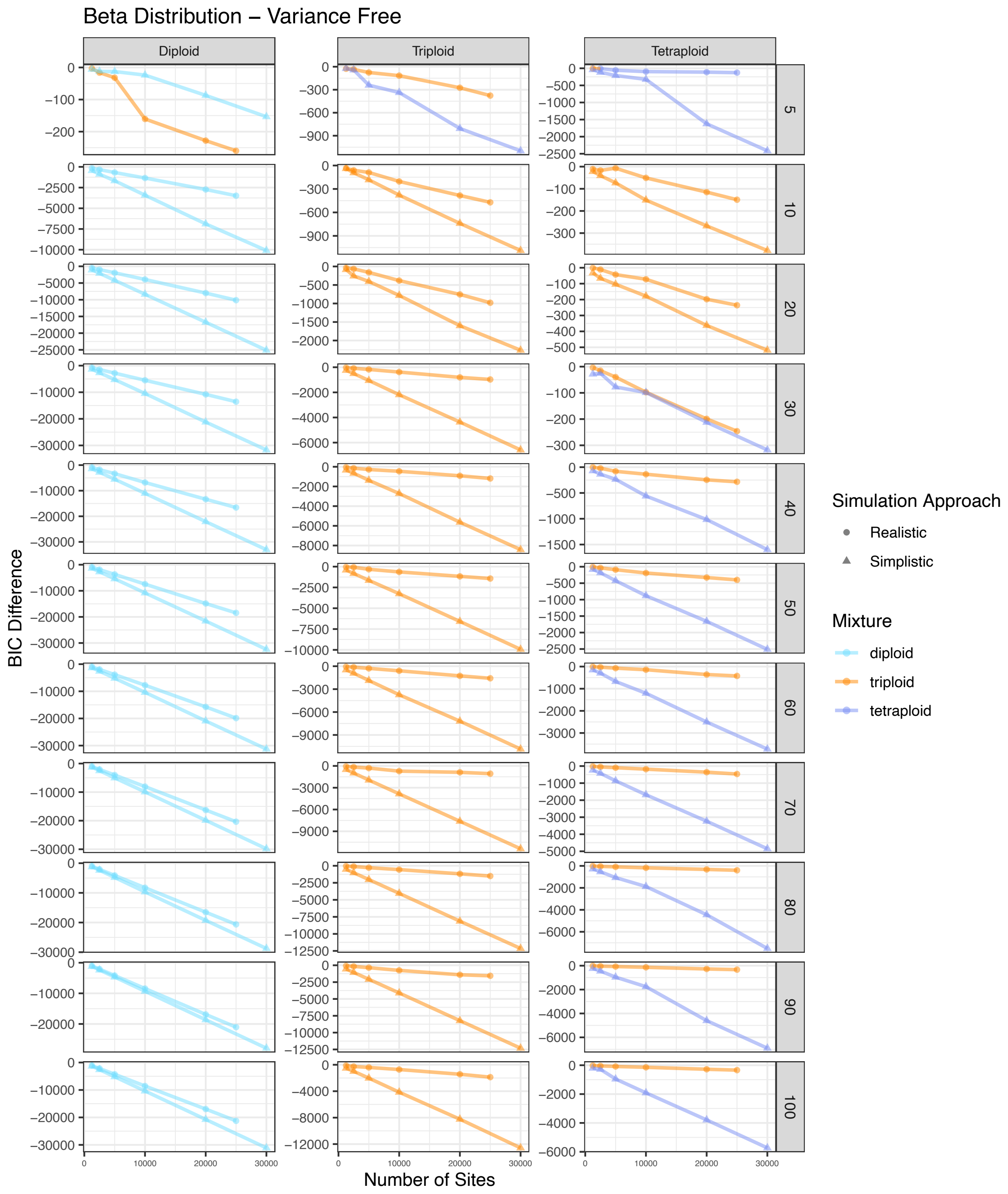


Figure S30. BIC score difference between the best and second best model for simulated diploid, triploid, and tetraploid samples across different numbers of sites for eleven different coverage amounts. The color of each point represents the best model. The shape of each point represents the approach used to simulate that sample. This represents a beta distribution with variance free.

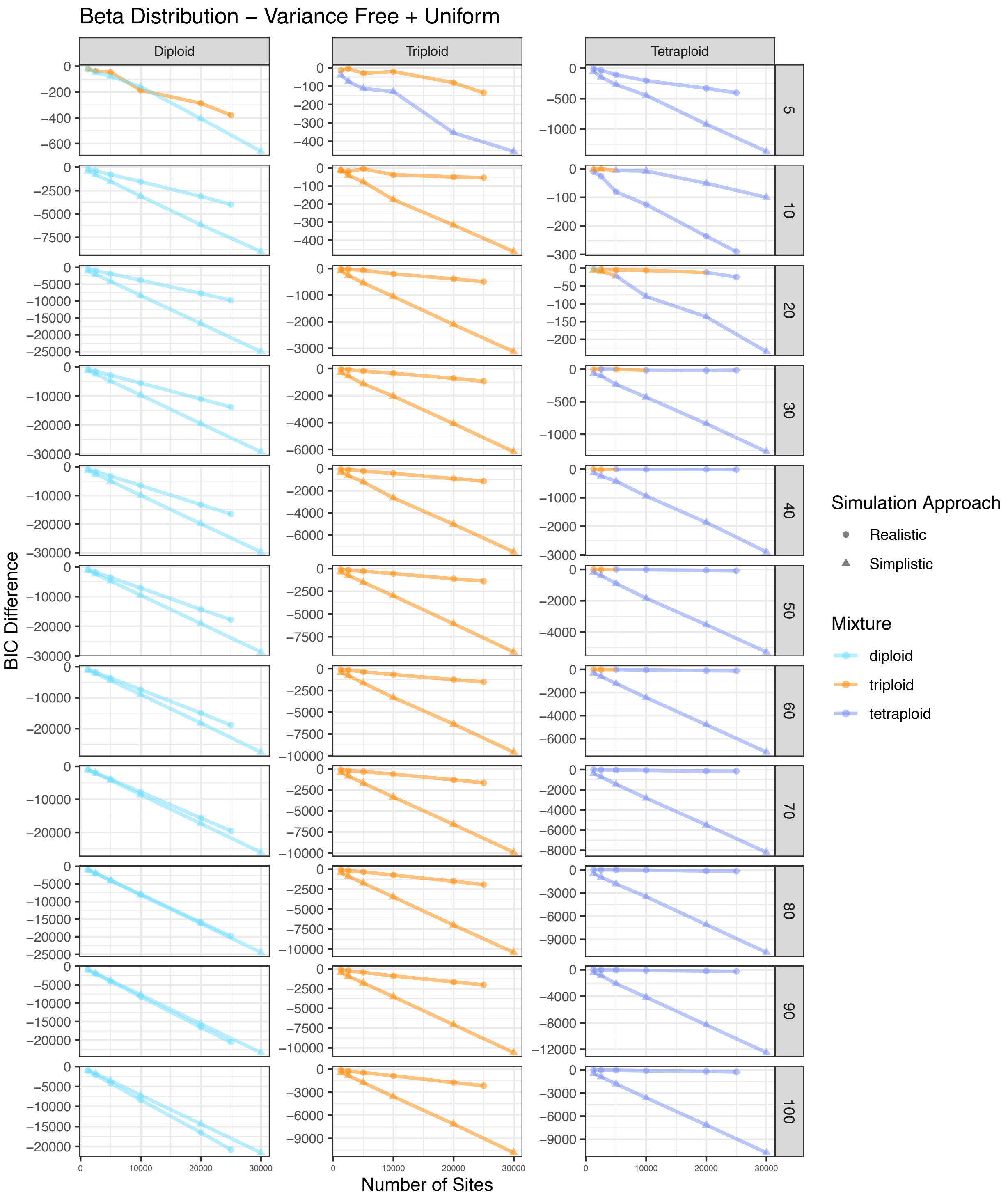


Figure S31. BIC score difference between the best and second best model for simulated diploid, triploid, and tetraploid samples across different numbers of sites for eleven different coverage amounts. The color of each point represents the best model. The shape of each point represents the approach used to simulate that sample. This represents a beta distribution with variance free with a uniform mixture.

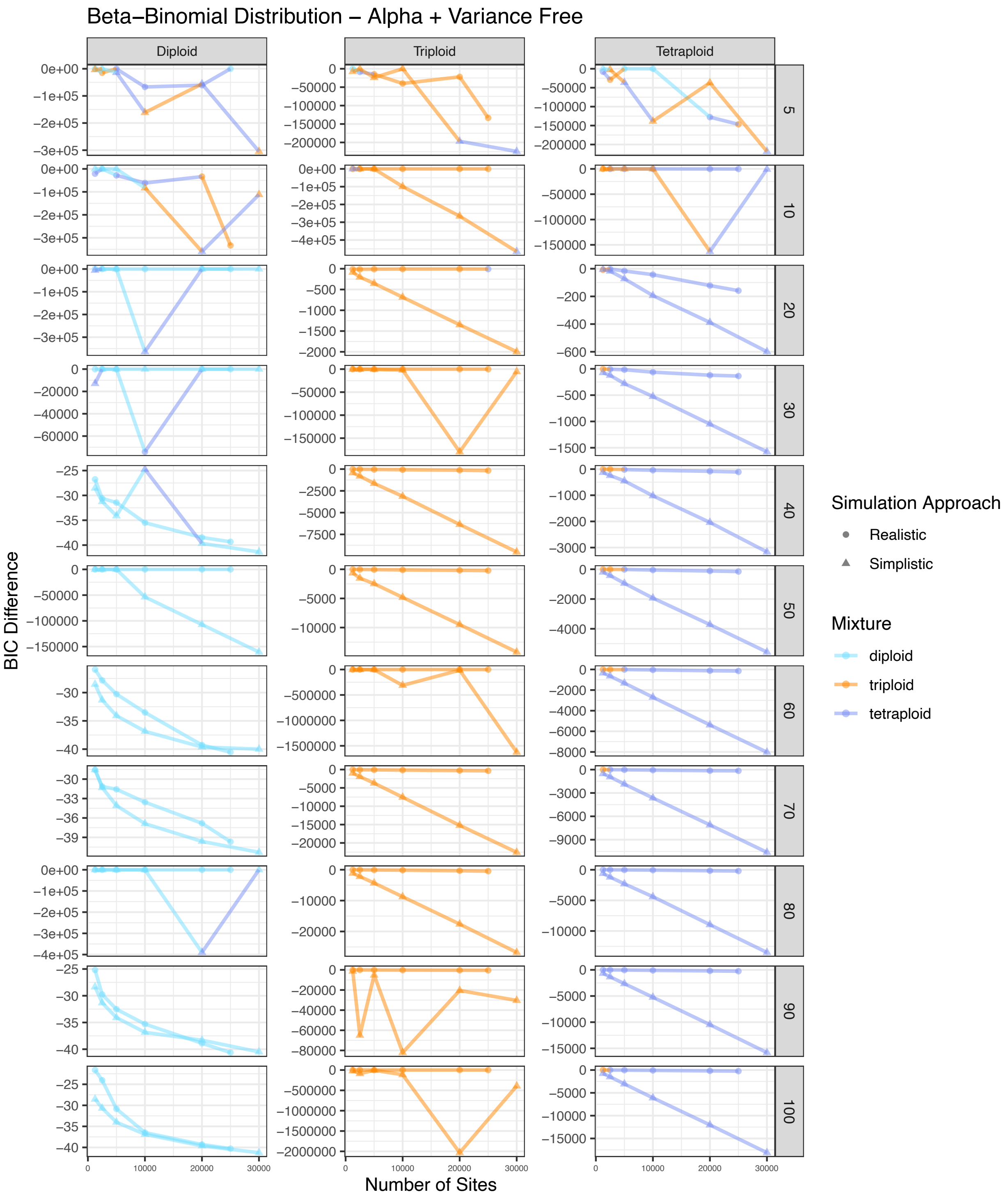


Figure S32. BIC score difference between the best and second best model for simulated diploid, triploid, and tetraploid samples across different numbers of sites for eleven different coverage amounts. The color of each point represents the best model. The shape of each point represents the approach used to simulate that sample. This represents a beta-binomial distribution with alpha and variance free.

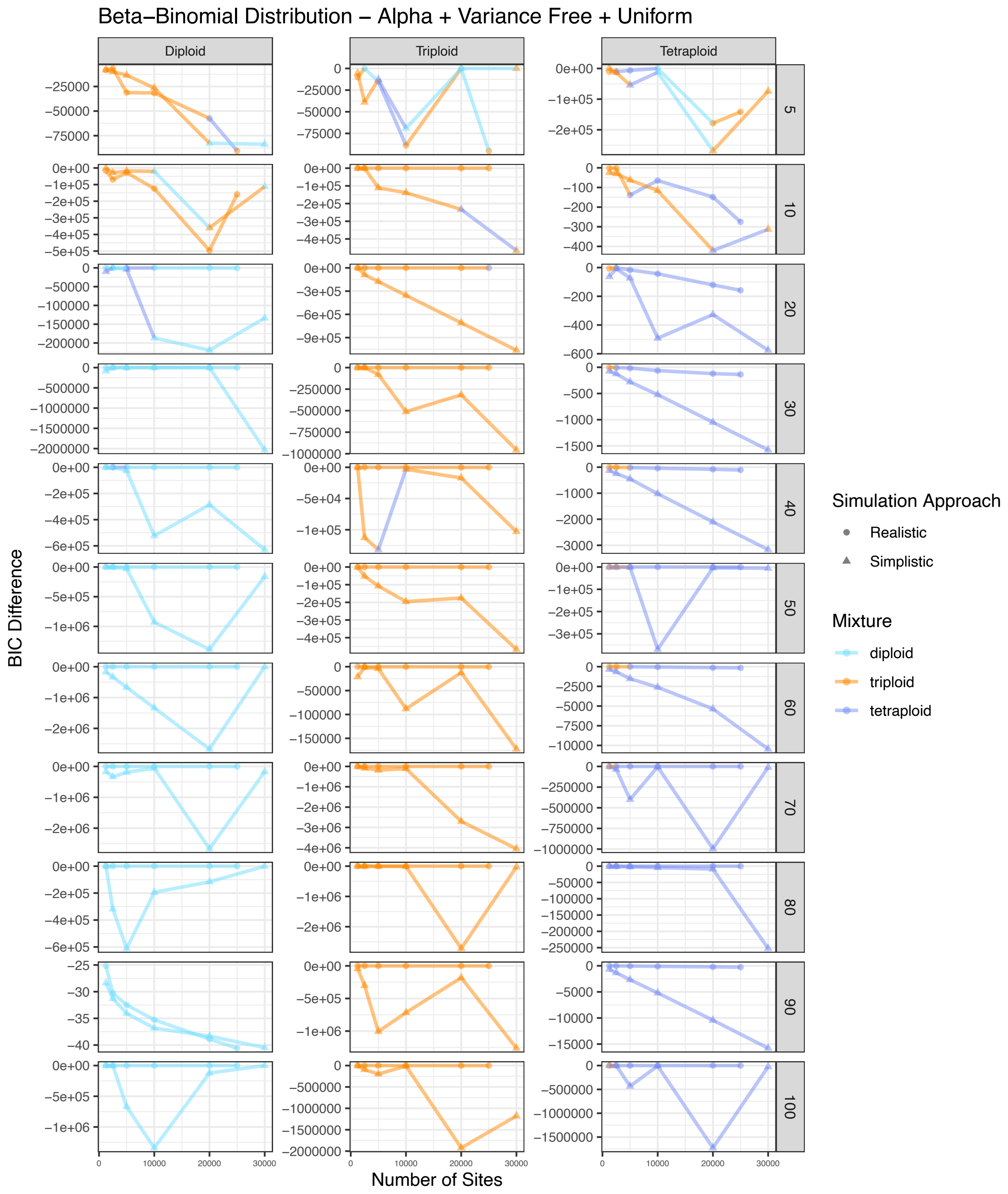


Figure S33. BIC score difference between the best and second best model for simulated diploid, triploid, and tetraploid samples across different numbers of sites for eleven different coverage amounts. The color of each point represents the best model. The shape of each point represents the approach used to simulate that sample. This represents a beta-binomial distribution with alpha and variance free with a uniform mixture.

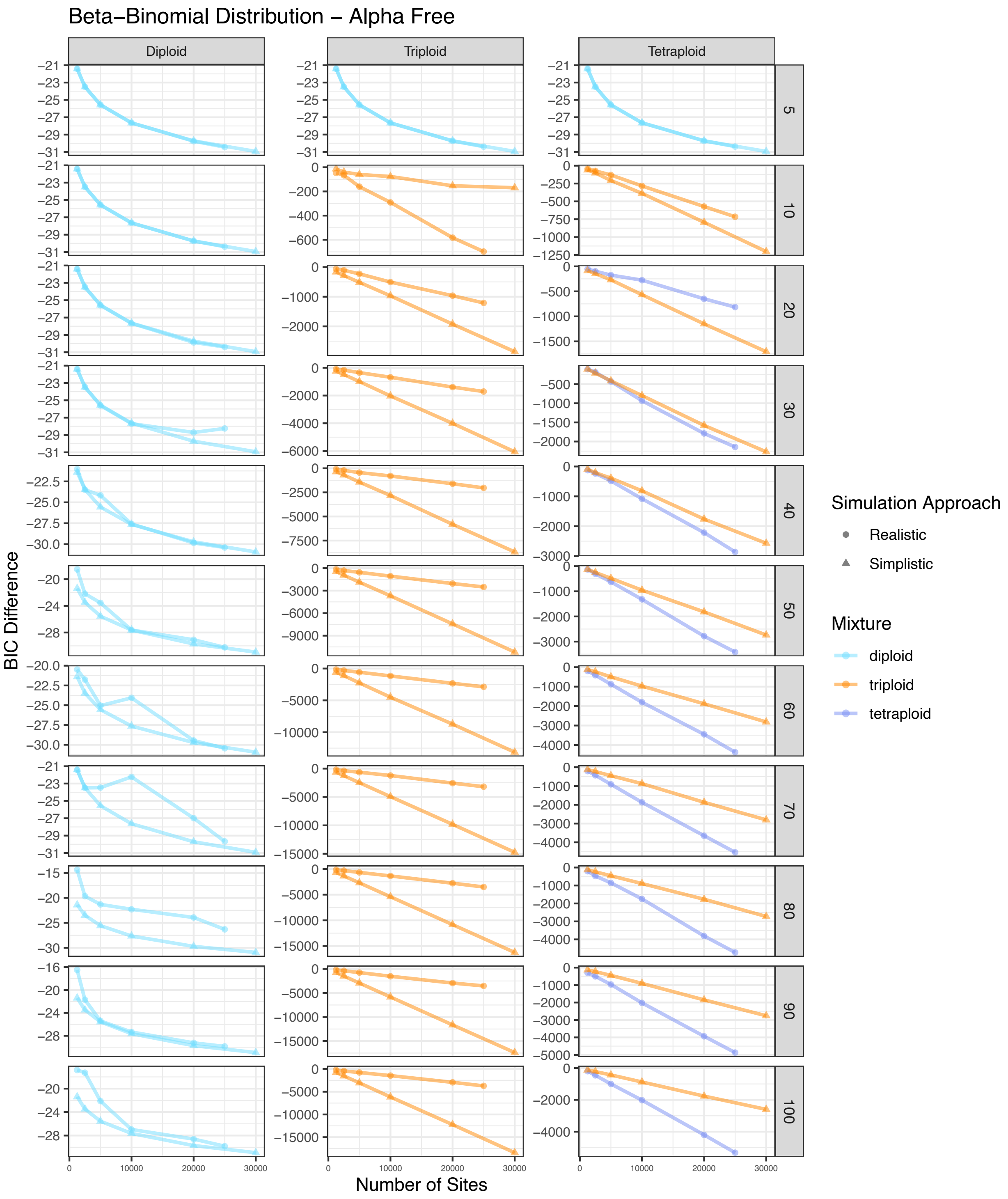


Figure S34. BIC score difference between the best and second best model for simulated diploid, triploid, and tetraploid samples across different numbers of sites for eleven different coverage amounts. The color of each point represents the best model. The shape of each point represents the approach used to simulate that sample. This represents a beta-binomial distribution with alpha free.

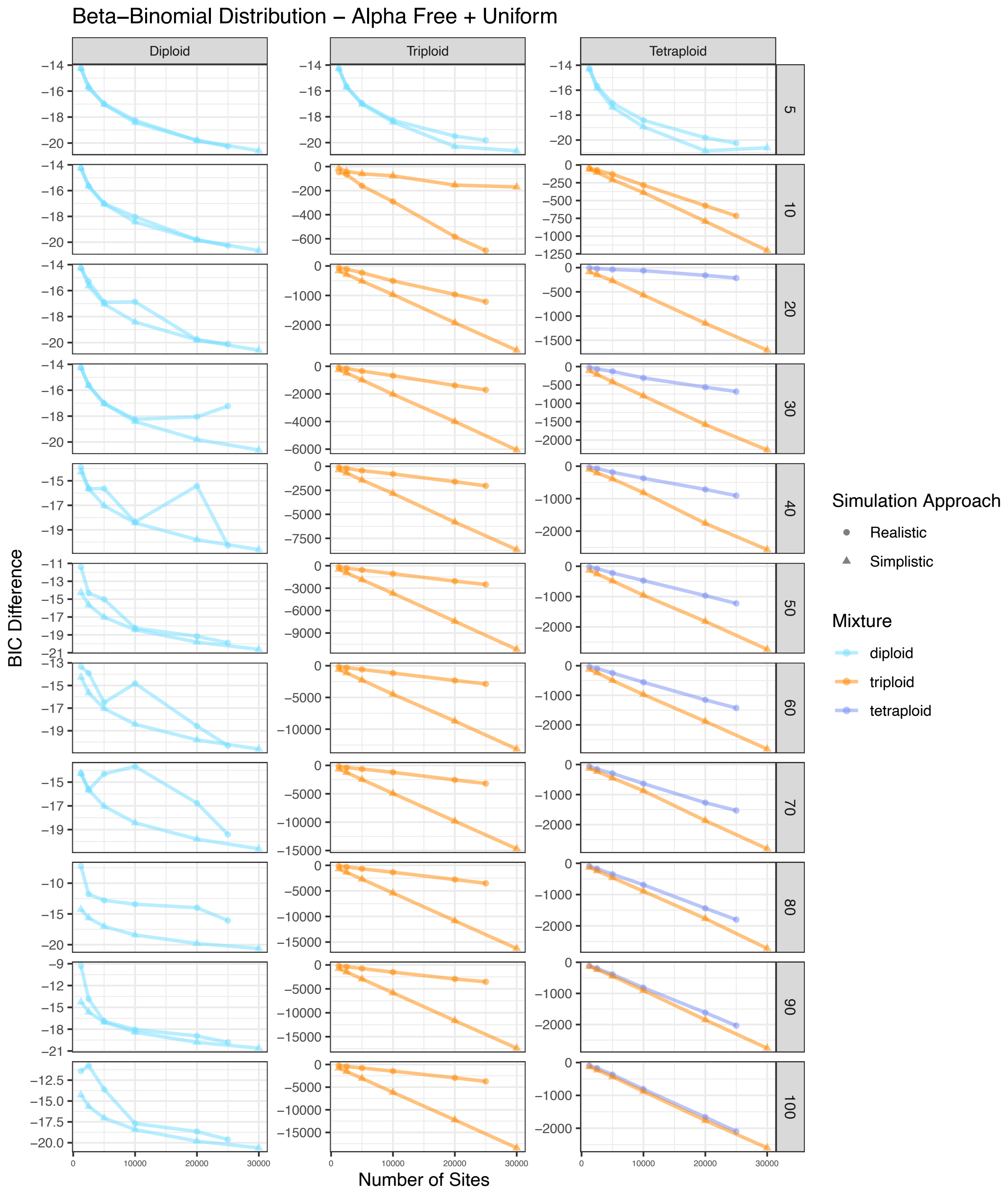


Figure S35. BIC score difference between the best and second best model for simulated diploid, triploid, and tetraploid samples across different numbers of sites for eleven different coverage amounts. The color of each point represents the best model. The shape of each point represents the approach used to simulate that sample. This represents a beta-binomial distribution with alpha free with a uniform mixture.

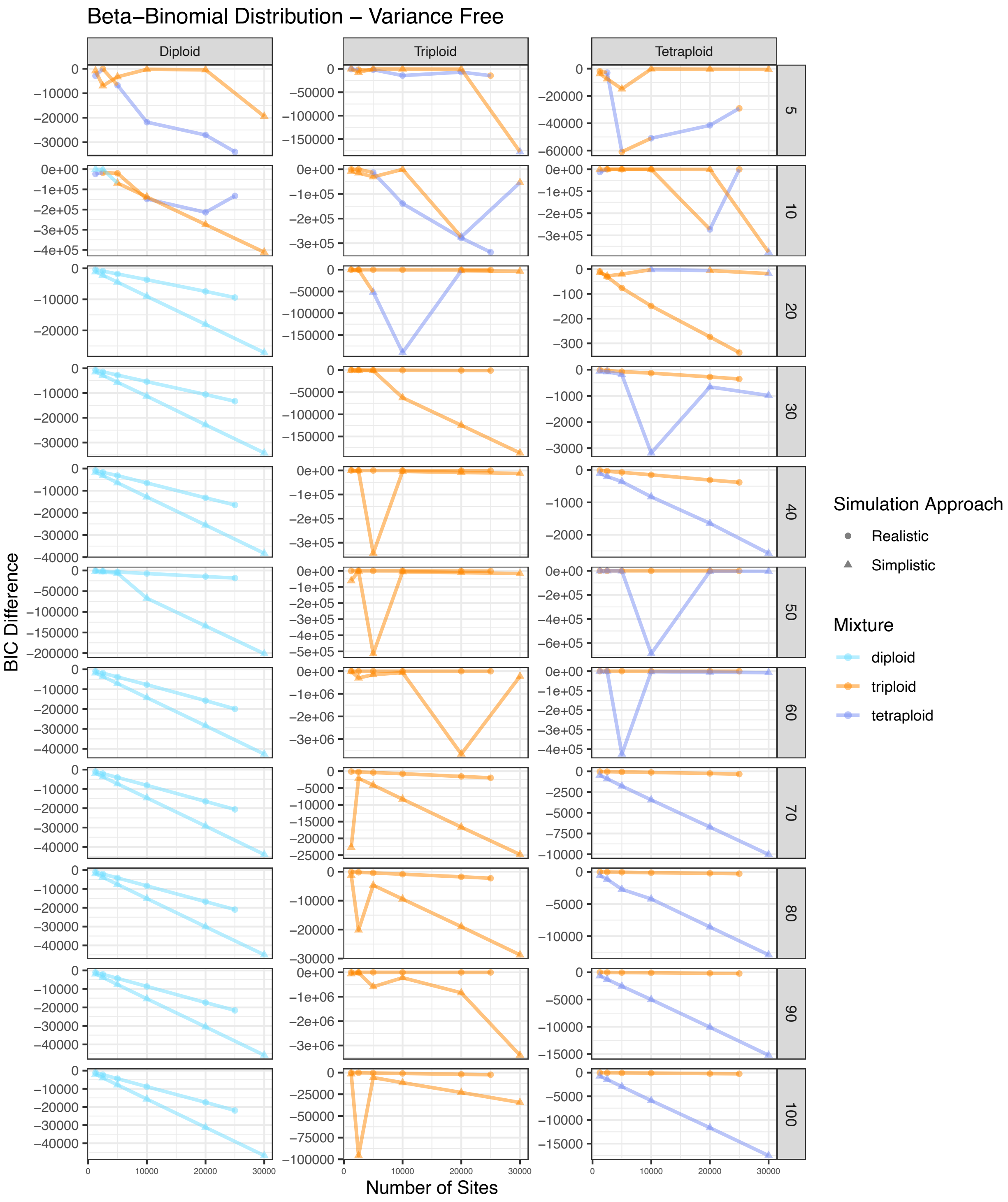


Figure S36. BIC score difference between the best and second best model for simulated diploid, triploid, and tetraploid samples across different numbers of sites for eleven different coverage amounts. The color of each point represents the best model. The shape of each point represents the approach used to simulate that sample. This represents a beta-binomial distribution with variance free.

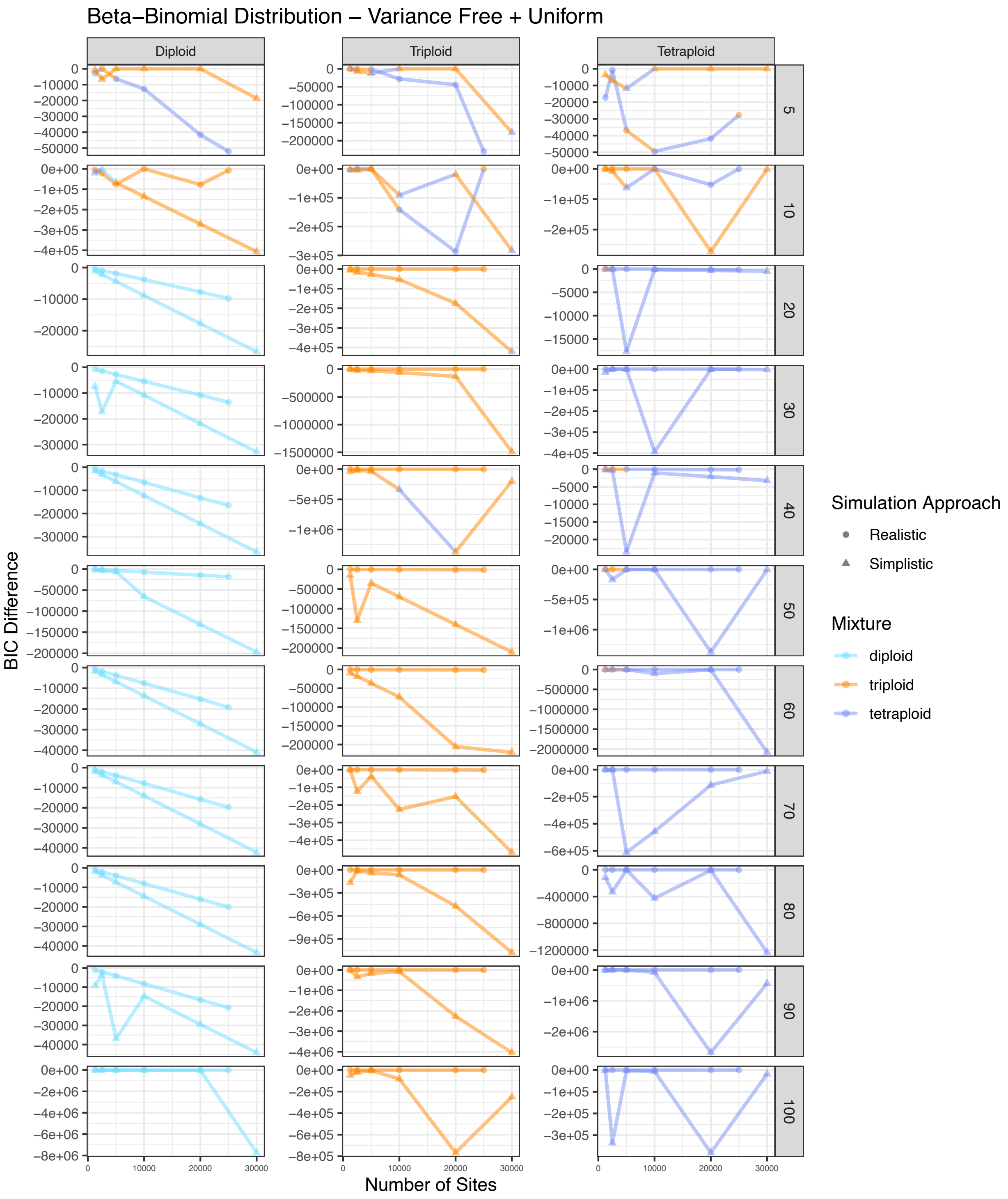


Figure S37. BIC score difference between the best and second best model for simulated diploid, triploid, and tetraploid samples across different numbers of sites for eleven different coverage amounts. The color of each point represents the best model. The shape of each point represents the approach used to simulate that sample. This represents a beta-binomial distribution with variance free with a uniform mixture.

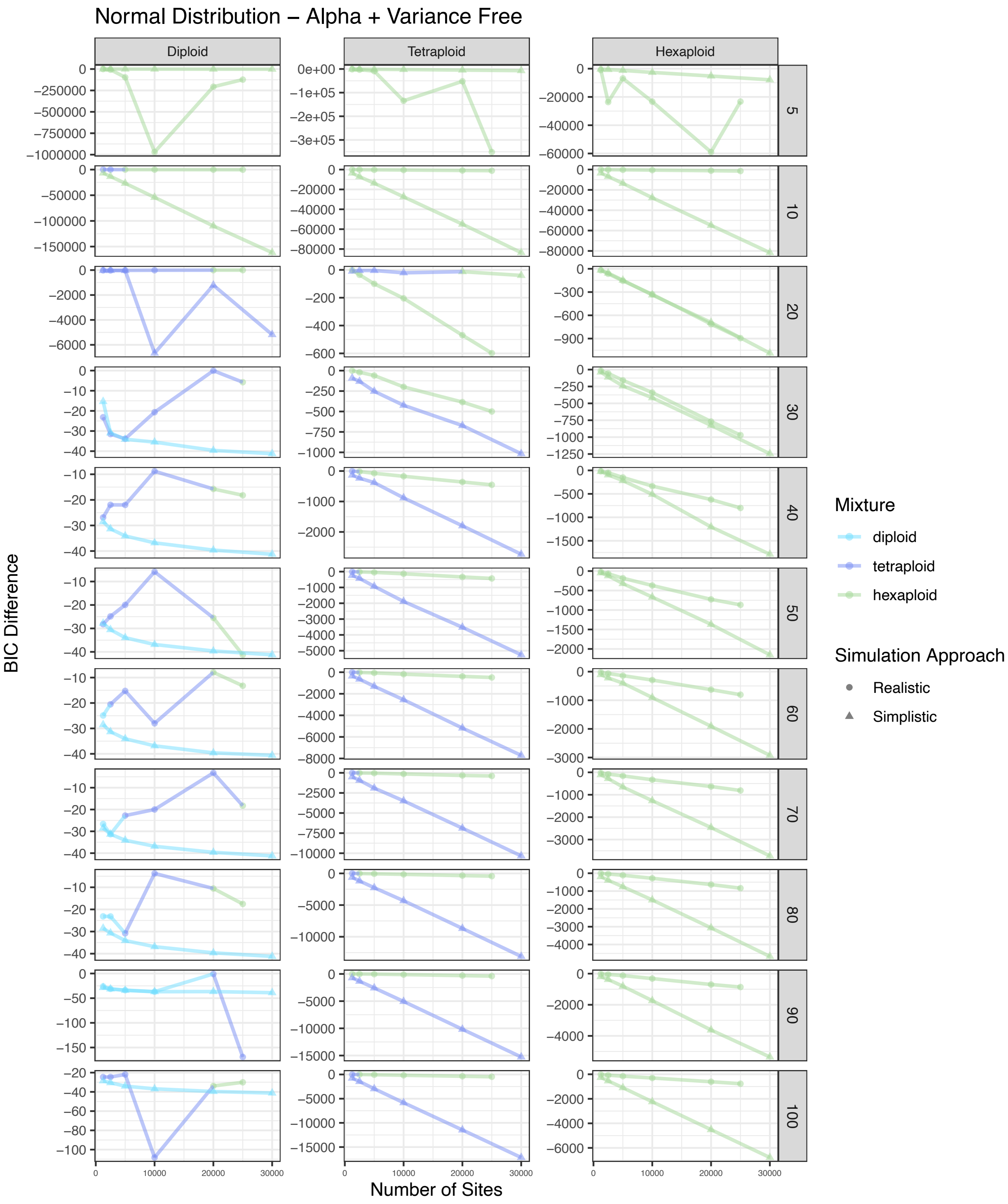


Figure S38. BIC score difference between the best and second best model for simulated diploid, tetraploid, and hexaploid samples across different numbers of sites for eleven different coverage amounts. The color of each point represents the best model. The shape of each point represents the approach used to simulate that sample. This represents a normal distribution with alpha and variance free.

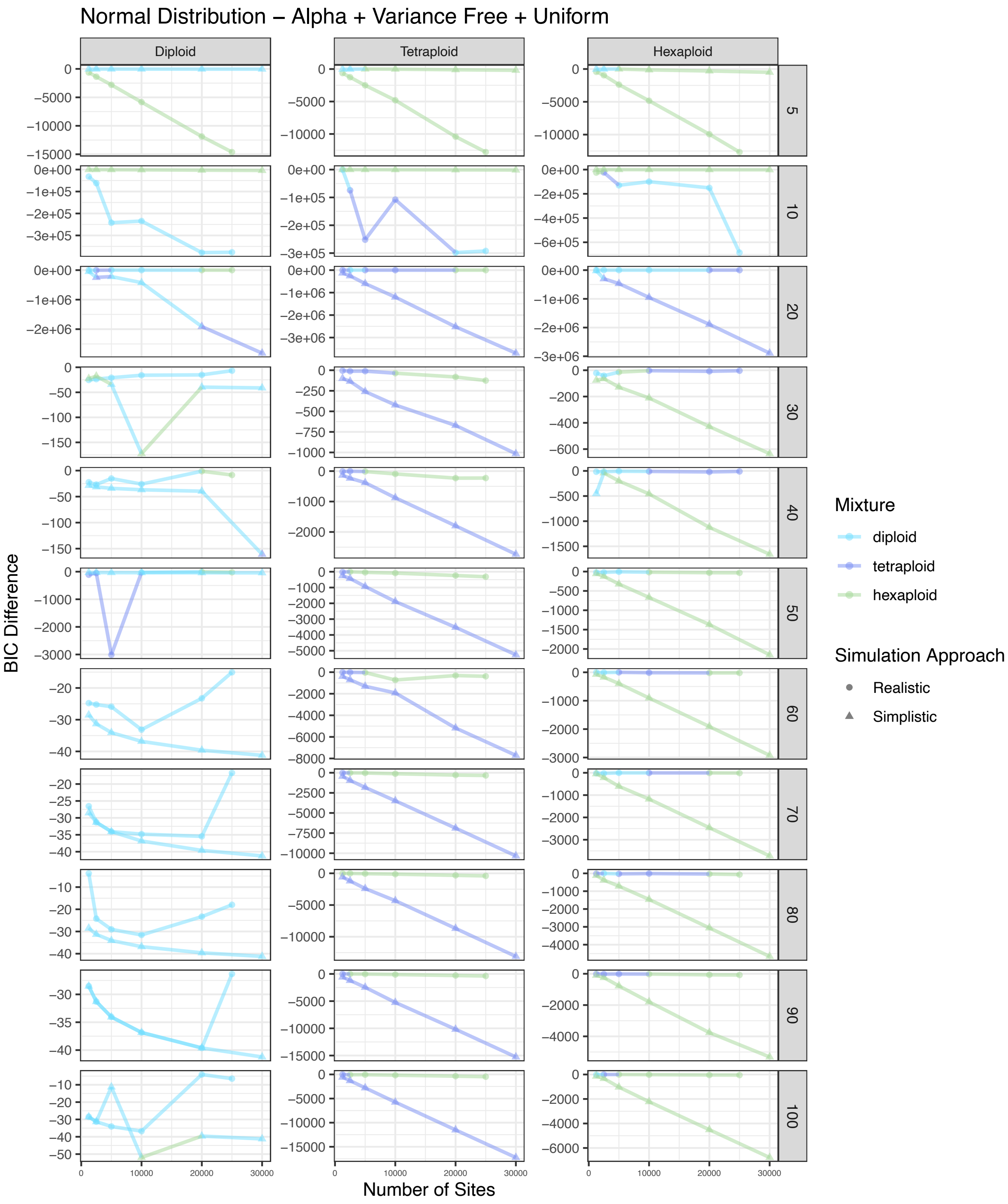


Figure S39. BIC score difference between the best and second best model for simulated diploid, tetraploid, and hexaploid samples across different numbers of sites for eleven different coverage amounts. The color of each point represents the best model. The shape of each point represents the approach used to simulate that sample. This represents a normal distribution with alpha and variance free with a uniform mixture.

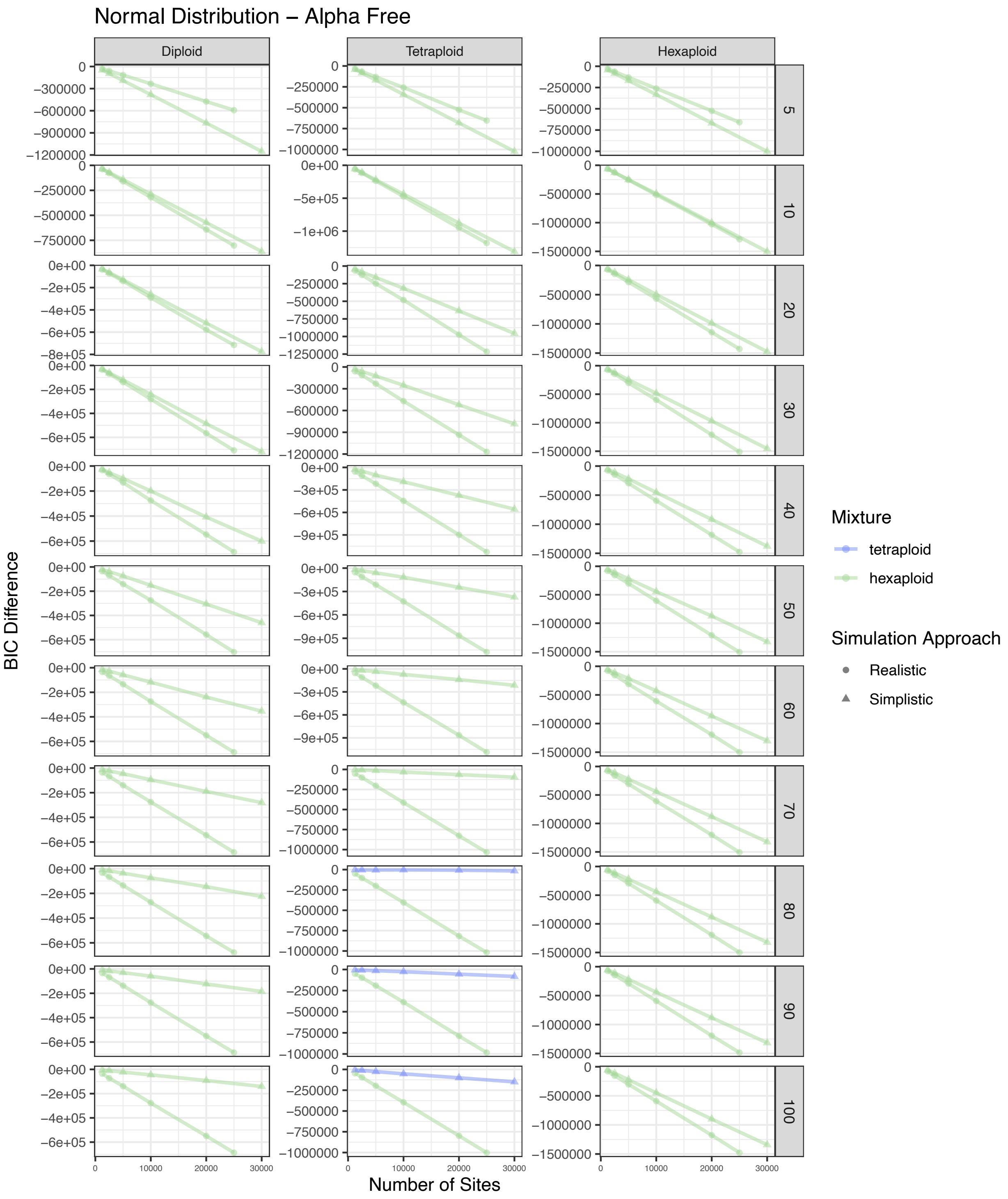


Figure S40. BIC score difference between the best and second best model for simulated diploid, tetraploid, and hexaploid samples across different numbers of sites for eleven different coverage amounts. The color of each point represents the best model. The shape of each point represents the approach used to simulate that sample. This represents a normal distribution with alpha free.

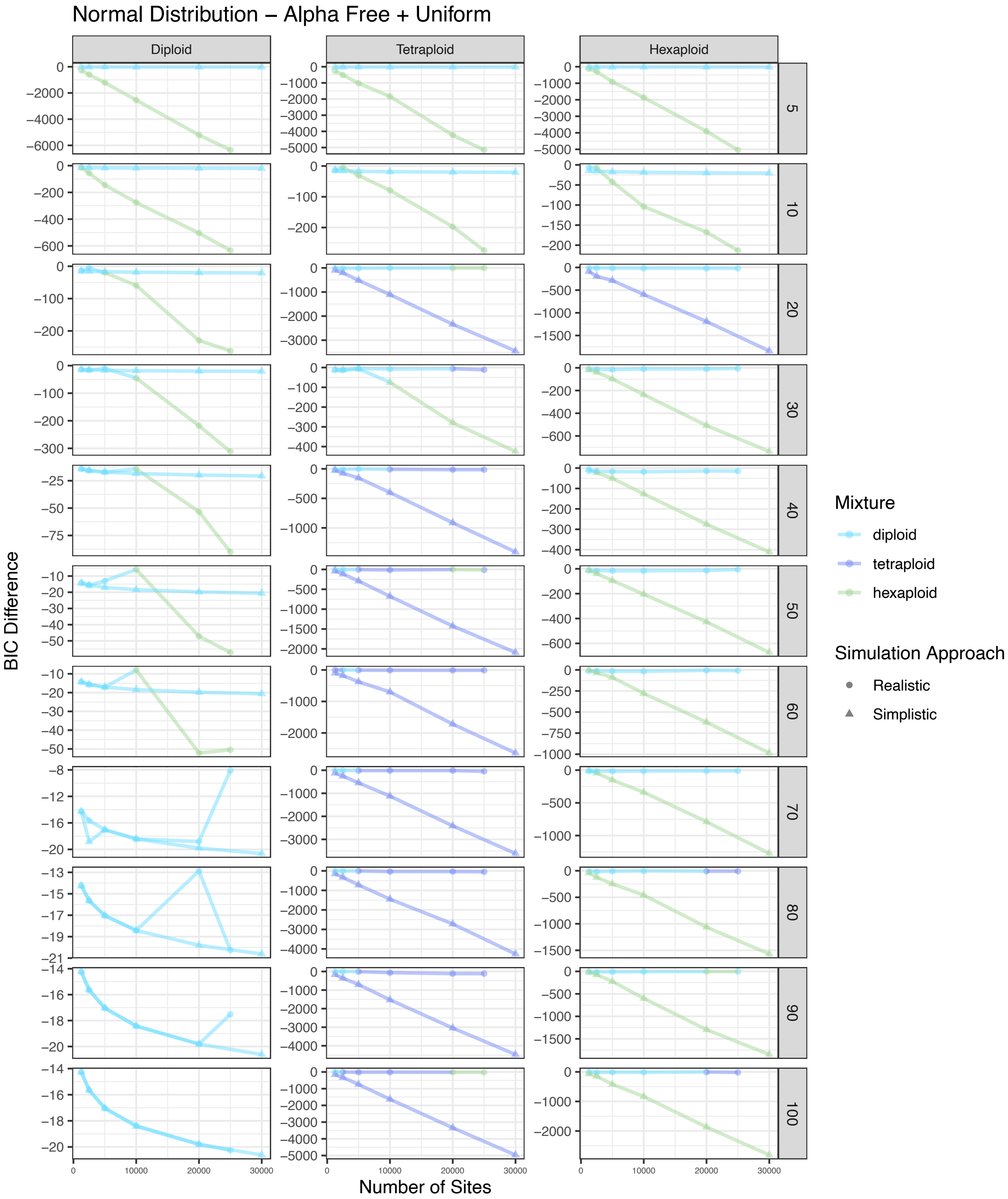


Figure S41. BIC score difference between the best and second best model for simulated diploid, tetraploid, and hexaploid samples across different numbers of sites for eleven different coverage amounts. The color of each point represents the best model. The shape of each point represents the approach used to simulate that sample. This represents a normal distribution with alpha free with a uniform mixture.

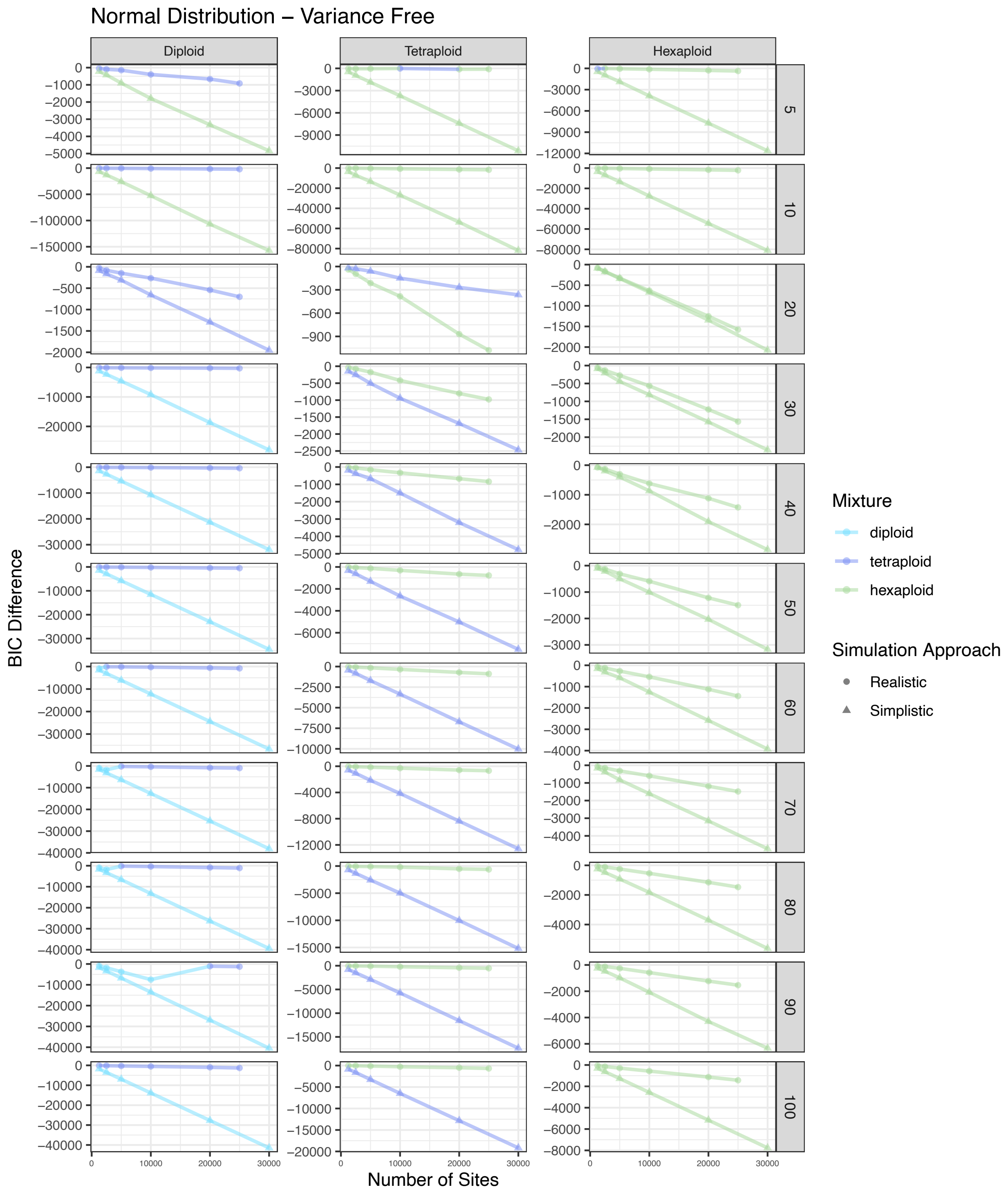


Figure S42. BIC score difference between the best and second best model for simulated diploid, tetraploid, and hexaploid samples across different numbers of sites for eleven different coverage amounts. The color of each point represents the best model. The shape of each point represents the approach used to simulate that sample. This represents a normal distribution with variance free.

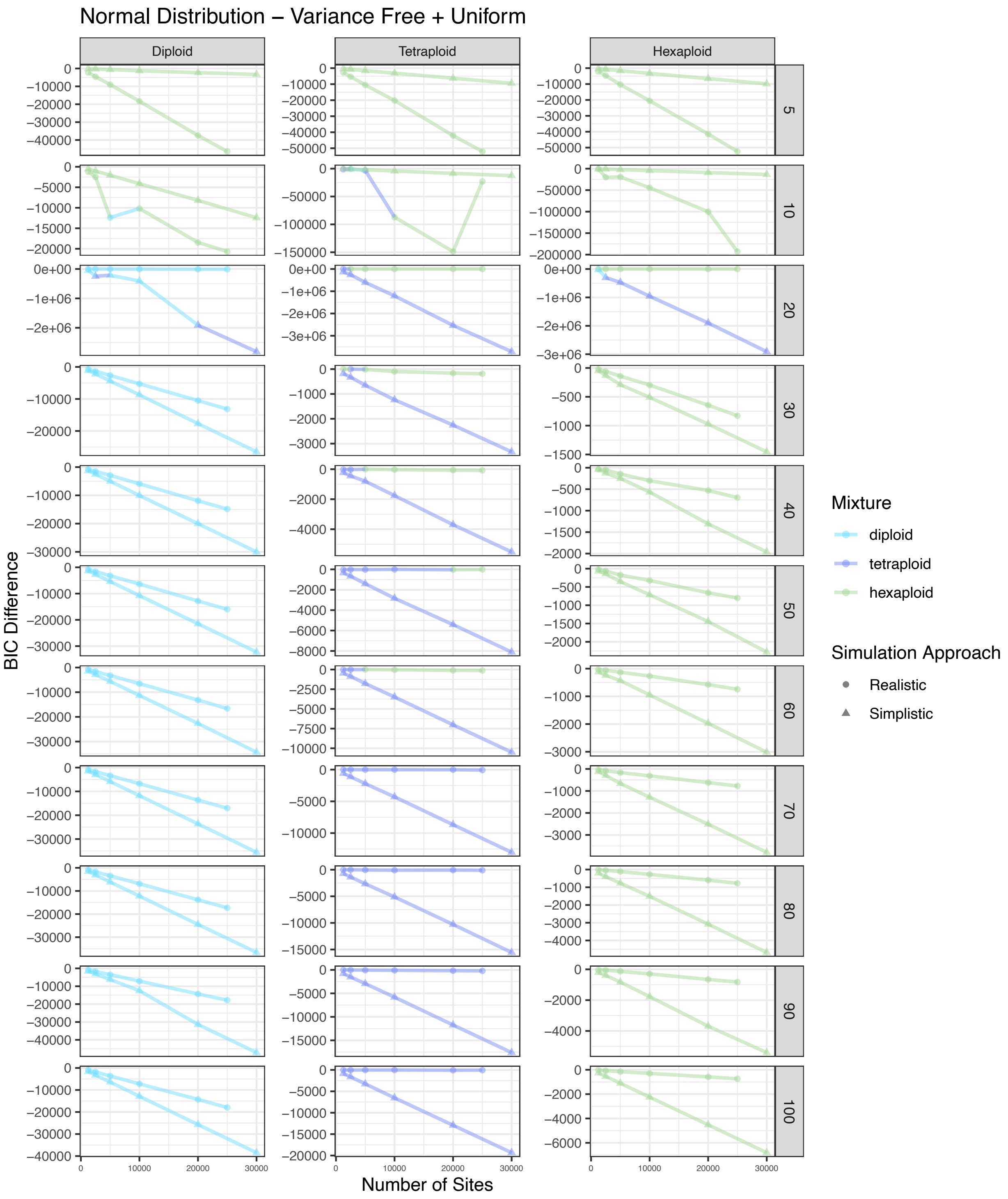


Figure S43. BIC score difference between the best and second best model for simulated diploid, tetraploid, and hexaploid samples across different numbers of sites for eleven different coverage amounts. The color of each point represents the best model. The shape of each point represents the approach used to simulate that sample. This represents a normal distribution with variance free with a uniform mixture.

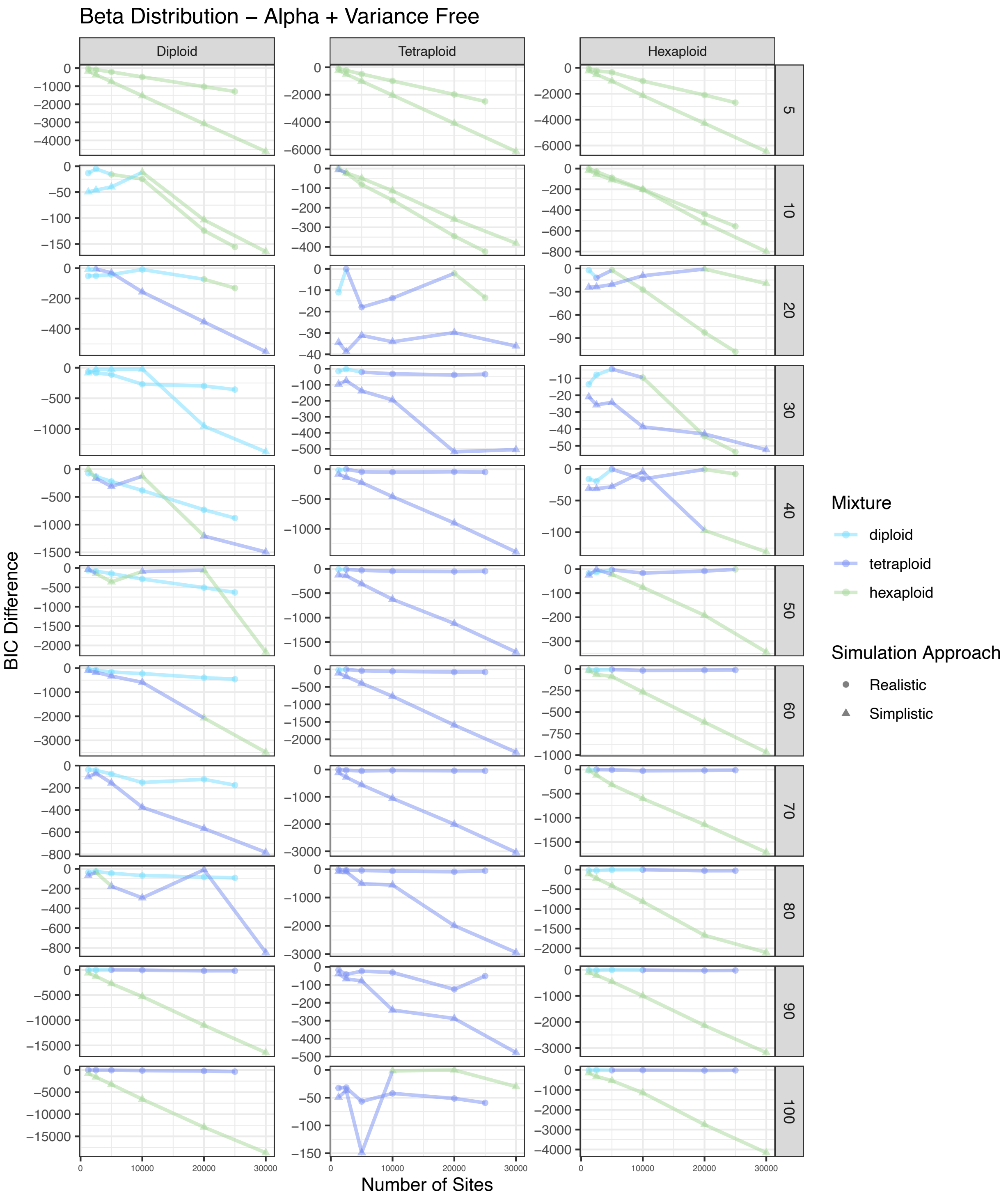


Figure S44. BIC score difference between the best and second best model for simulated diploid, tetraploid, and hexaploid samples across different numbers of sites for eleven different coverage amounts. The color of each point represents the best model. The shape of each point represents the approach used to simulate that sample. This represents a beta distribution with alpha and variance free.

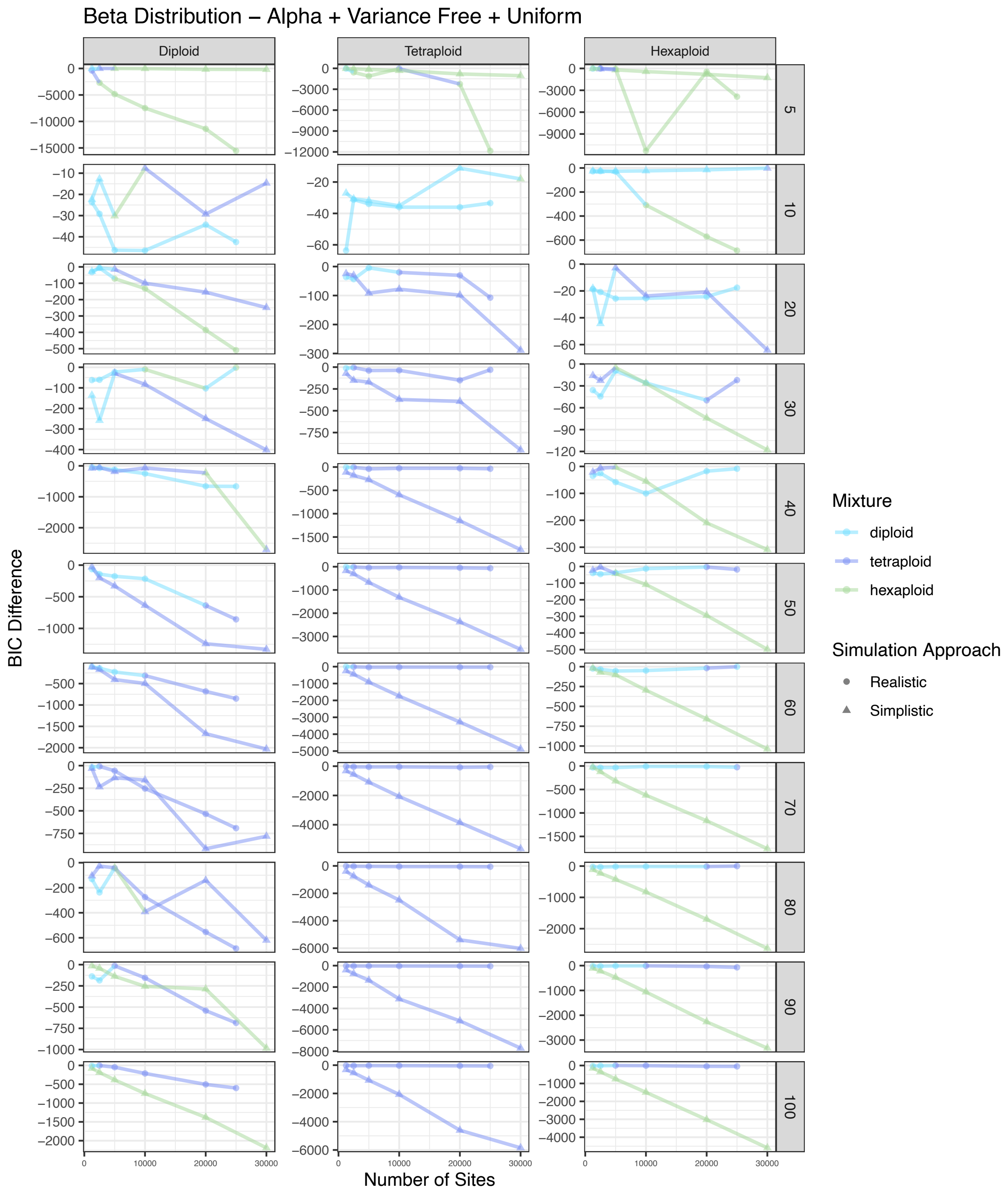


Figure S45. BIC score difference between the best and second best model for simulated diploid, tetraploid, and hexaploid samples across different numbers of sites for eleven different coverage amounts. The color of each point represents the best model. The shape of each point represents the approach used to simulate that sample. This represents a beta distribution with alpha and variance free with a uniform mixture.

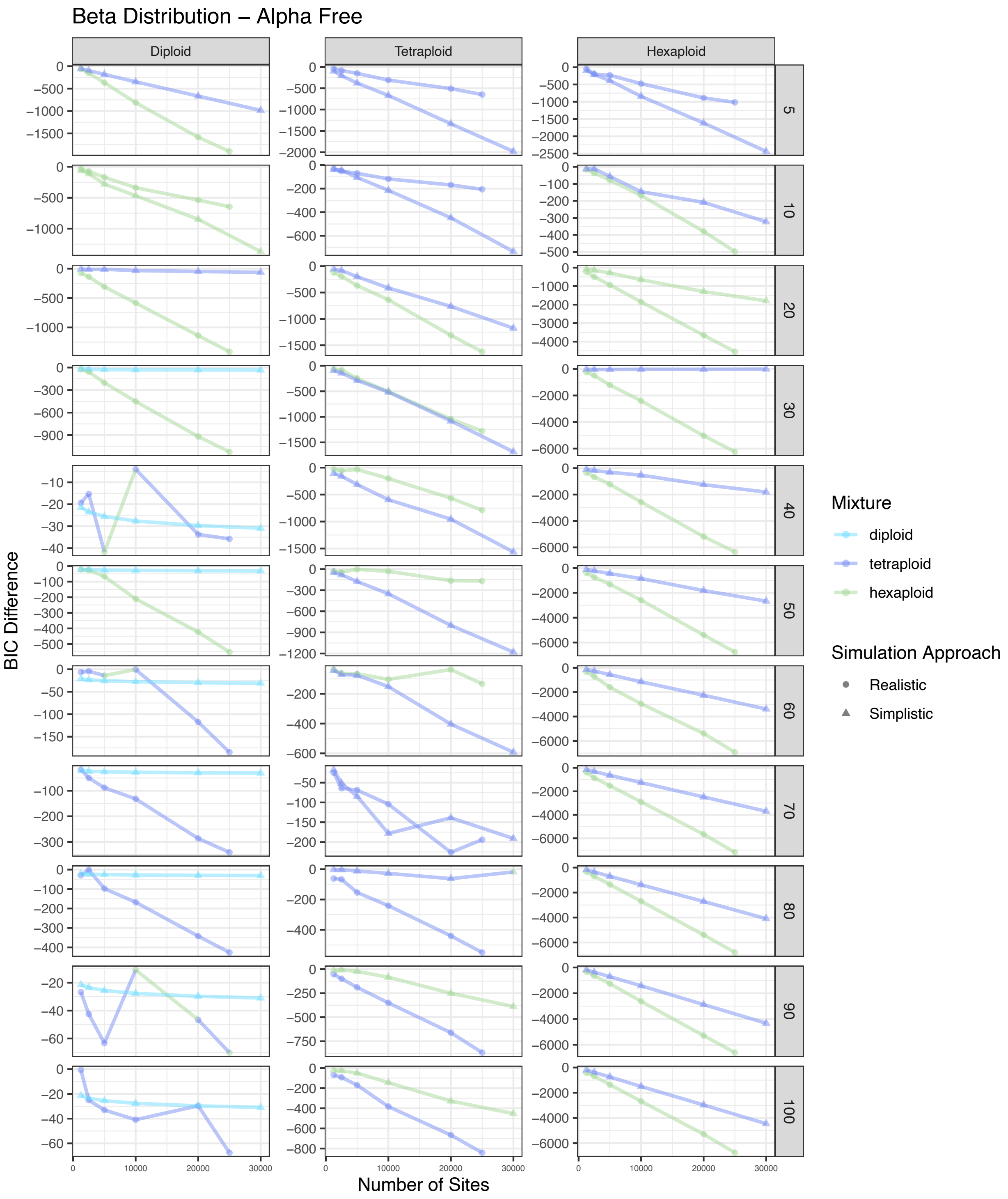


Figure S46. BIC score difference between the best and second best model for simulated diploid, tetraploid, and hexaploid samples across different numbers of sites for eleven different coverage amounts. The color of each point represents the best model. The shape of each point represents the approach used to simulate that sample. This represents a beta distribution with alpha free.

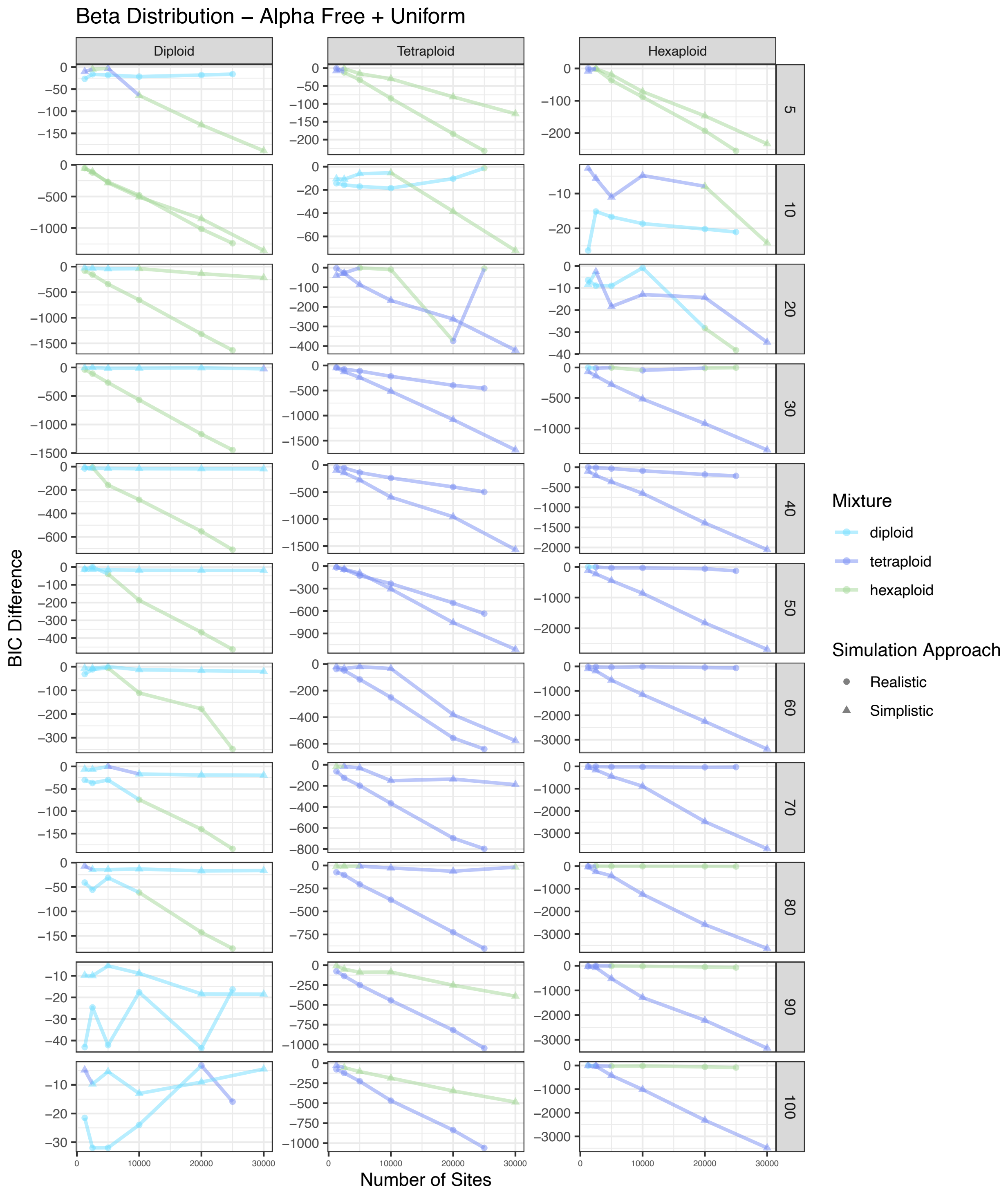


Figure S47. BIC score difference between the best and second best model for simulated diploid, tetraploid, and hexaploid samples across different numbers of sites for eleven different coverage amounts. The color of each point represents the best model. The shape of each point represents the approach used to simulate that sample. This represents a beta distribution with alpha free with a uniform mixture.

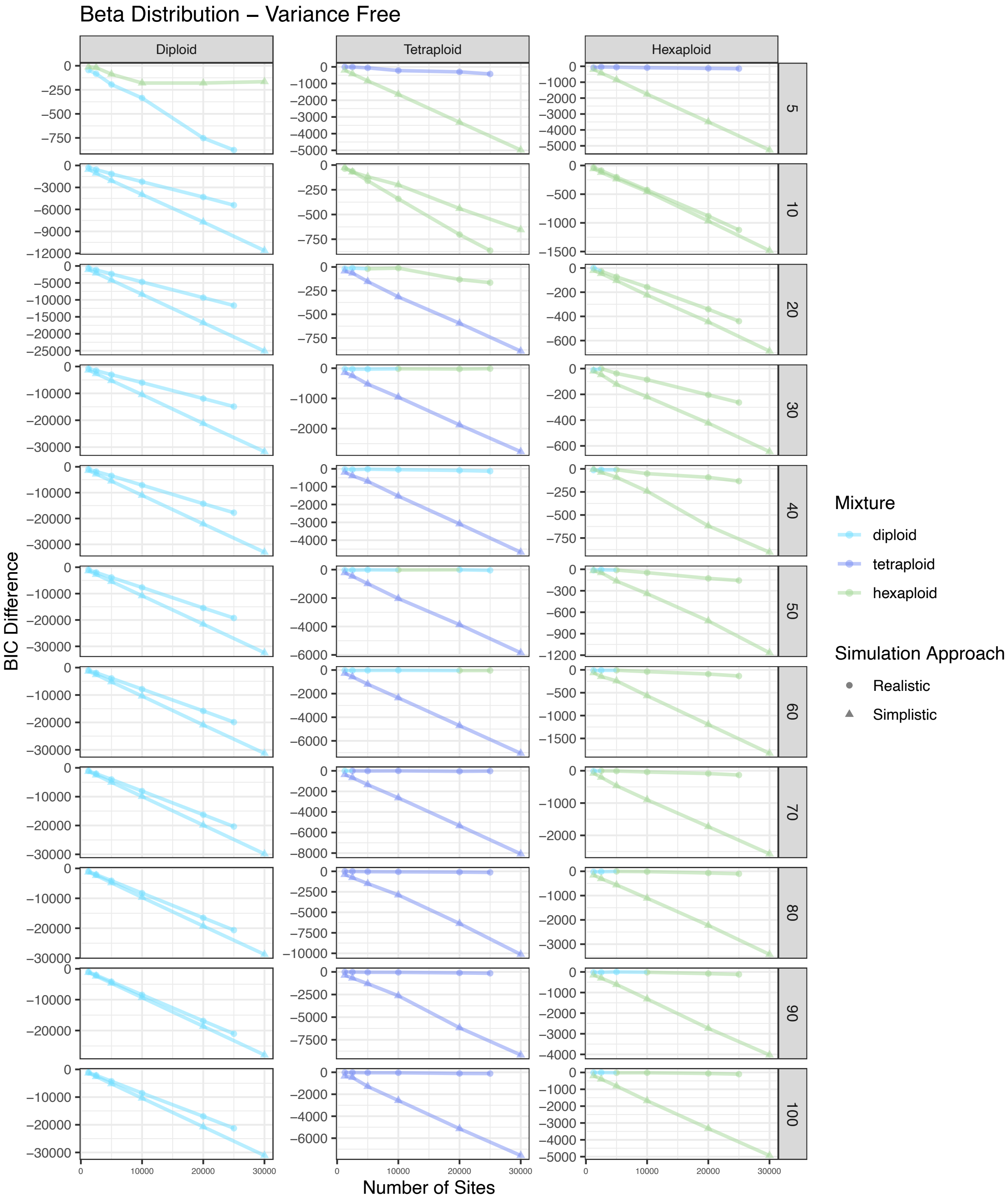


Figure S48. BIC score difference between the best and second best model for simulated diploid, tetraploid, and hexaploid samples across different numbers of sites for eleven different coverage amounts. The color of each point represents the best model. The shape of each point represents the approach used to simulate that sample. This represents a beta distribution with variance free.

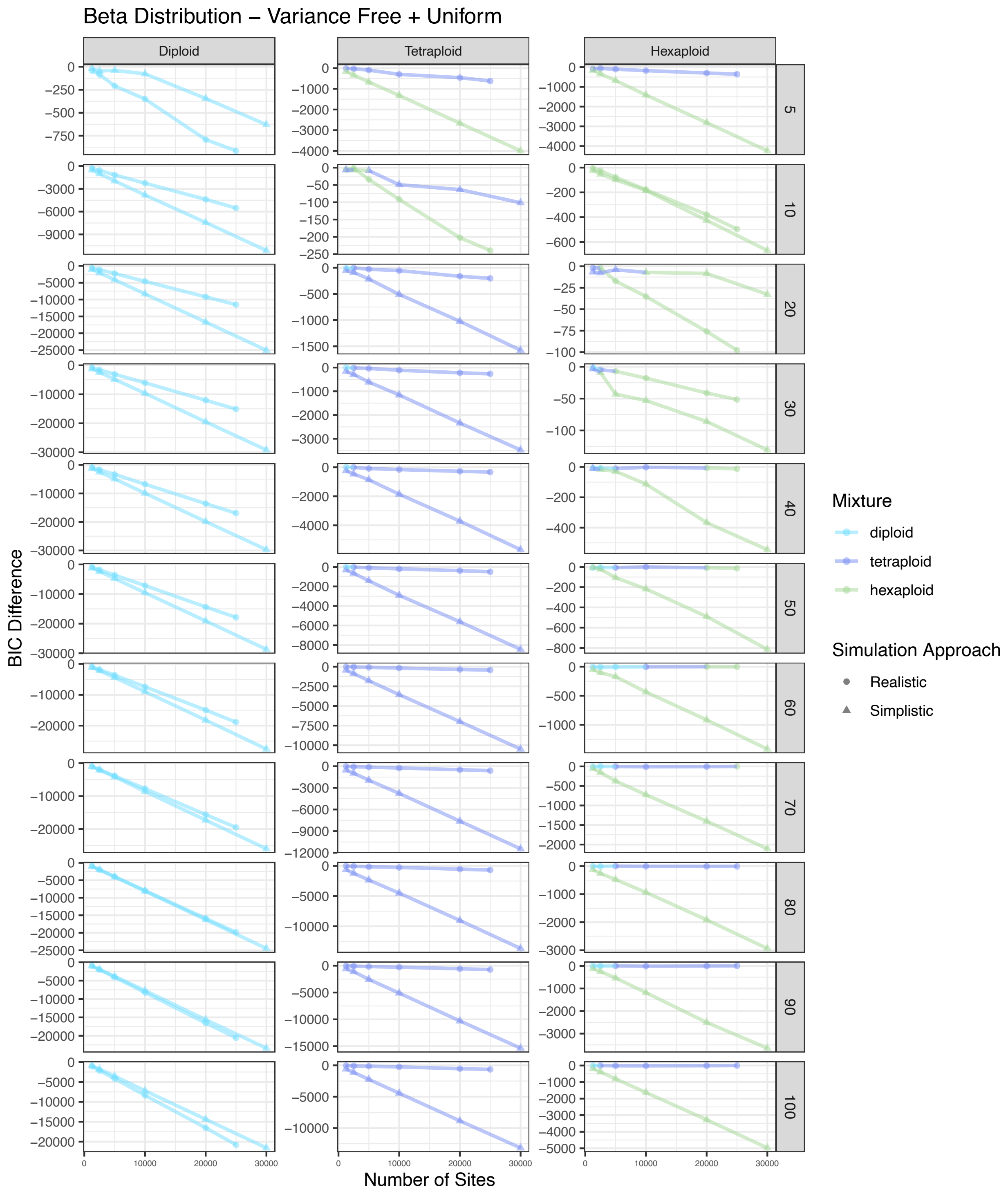


Figure S49. BIC score difference between the best and second best model for simulated diploid, tetraploid, and hexaploid samples across different numbers of sites for eleven different coverage amounts. The color of each point represents the best model. The shape of each point represents the approach used to simulate that sample. This represents a beta distribution with variance free with a uniform mixture.

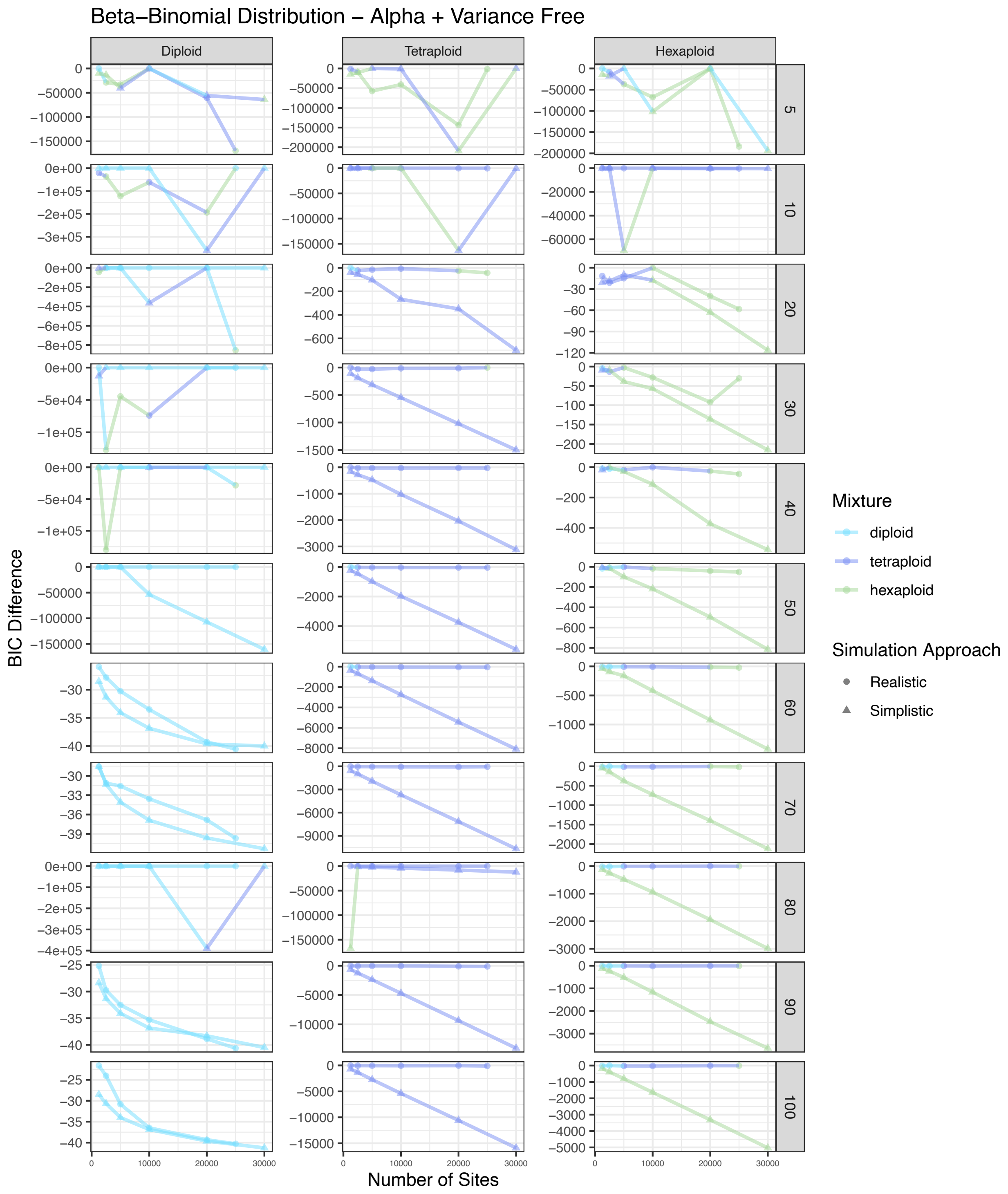


Figure S50. BIC score difference between the best and second best model for simulated diploid, tetraploid, and hexaploid samples across different numbers of sites for eleven different coverage amounts. The color of each point represents the best model. The shape of each point represents the approach used to simulate that sample. This represents a beta-binomial distribution with alpha and variance free.

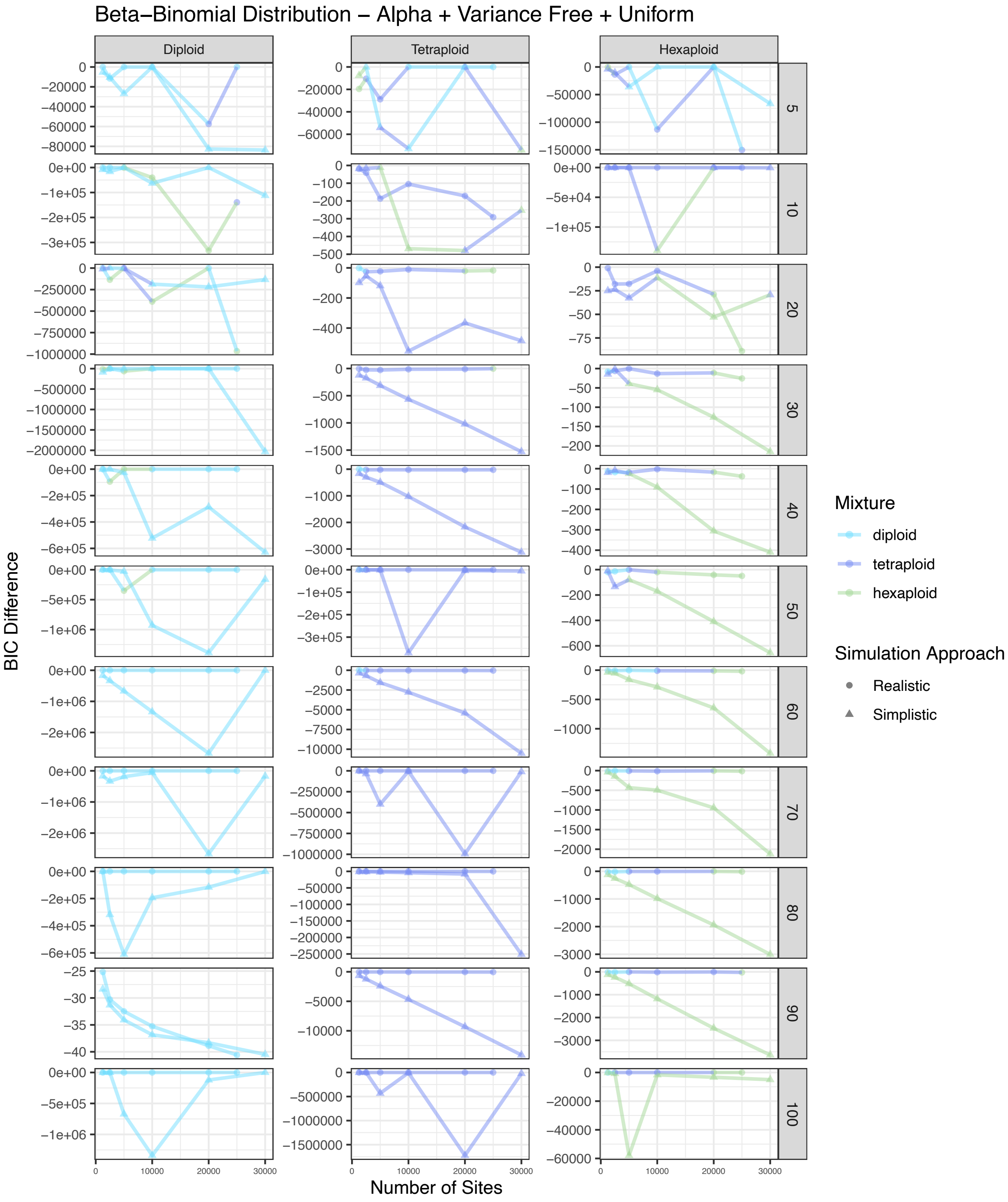


Figure S51. BIC score difference between the best and second best model for simulated diploid, tetraploid, and hexaploid samples across different numbers of sites for eleven different coverage amounts. The color of each point represents the best model. The shape of each point represents the approach used to simulate that sample. This represents a beta-binomial distribution with alpha and variance free with a uniform mixture.

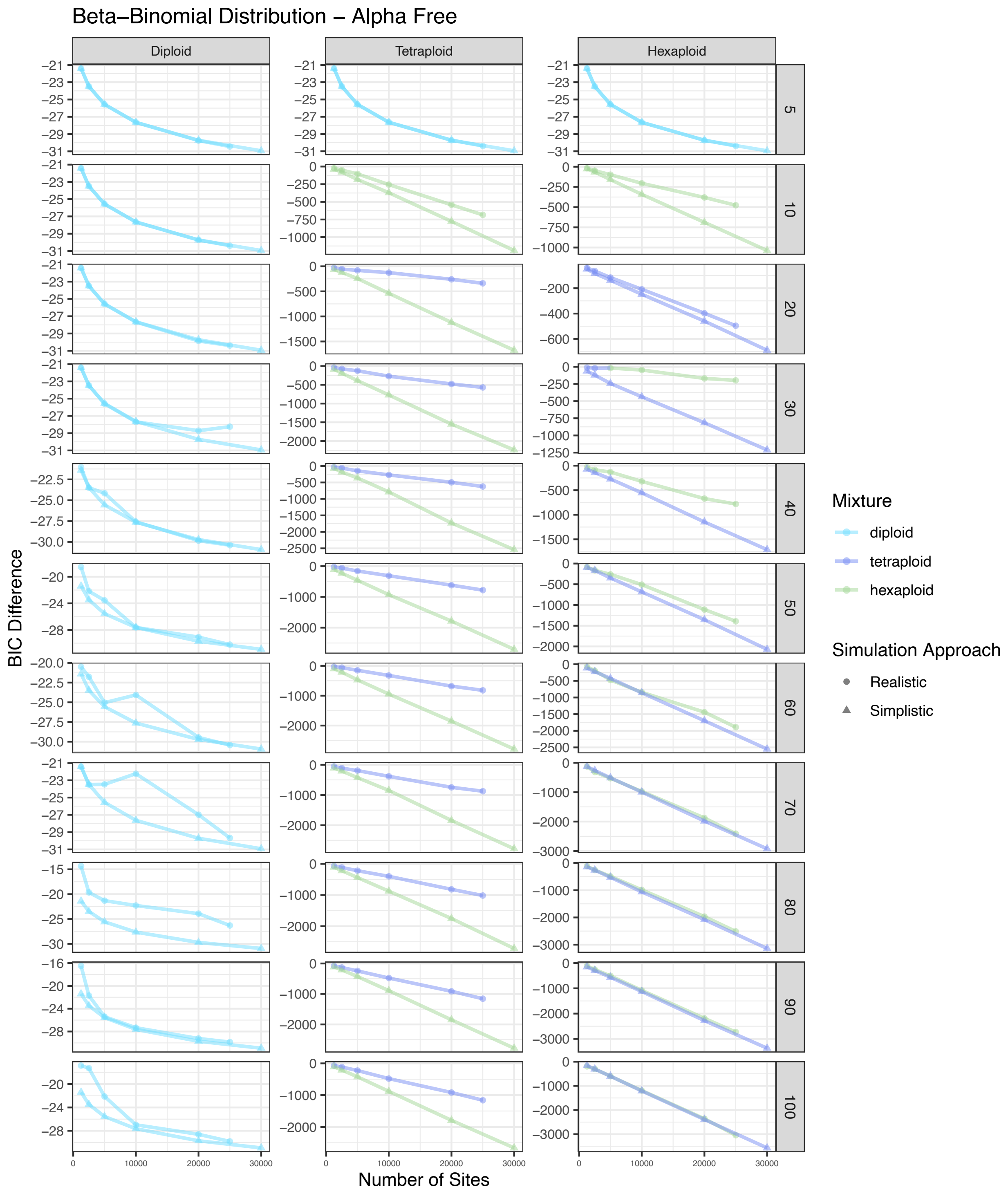


Figure S52. BIC score difference between the best and second best model for simulated diploid, tetraploid, and hexaploid samples across different numbers of sites for eleven different coverage amounts. The color of each point represents the best model. The shape of each point represents the approach used to simulate that sample. This represents a beta-binomial distribution with alpha free.

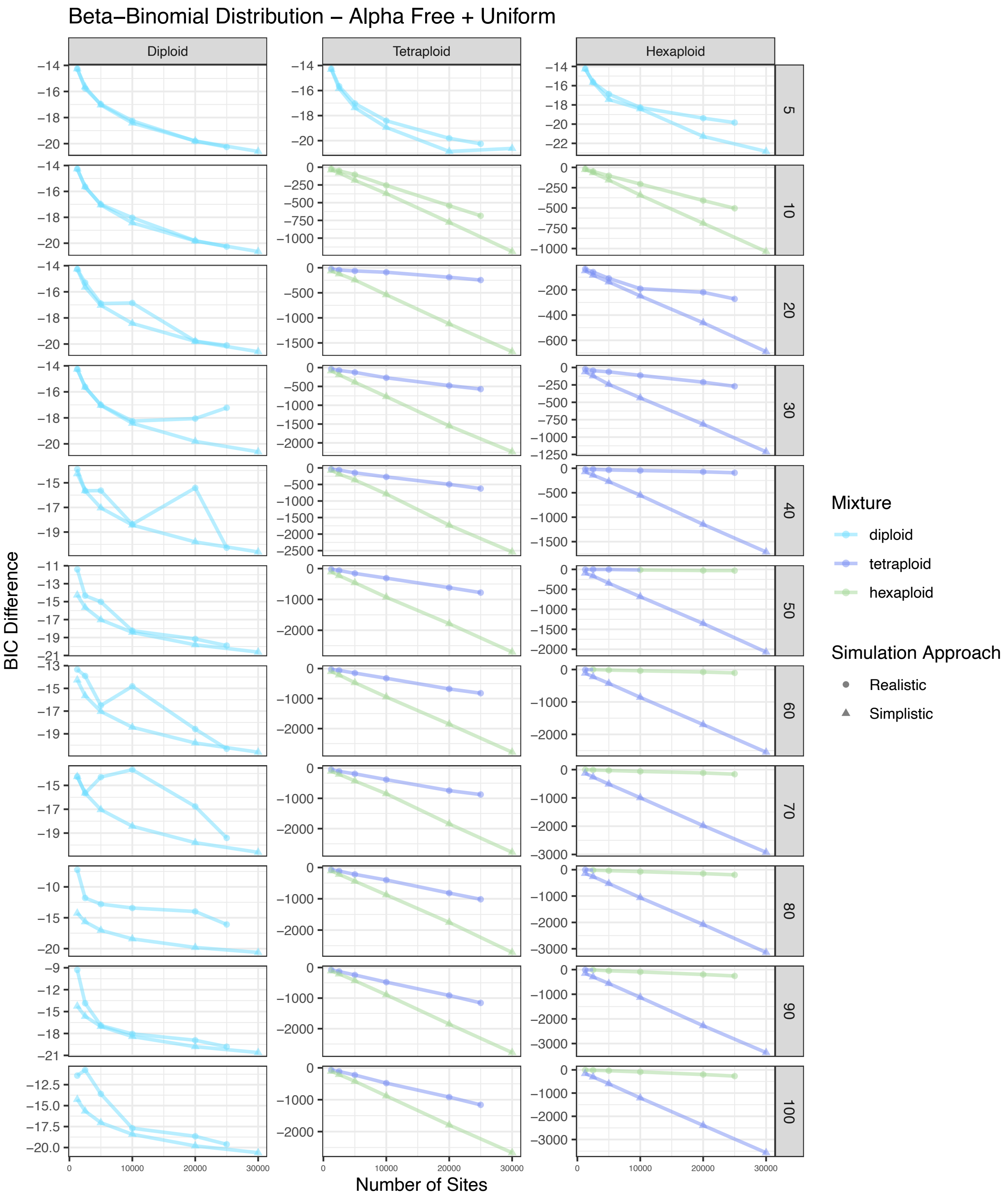


Figure S53. BIC score difference between the best and second best model for simulated diploid, tetraploid, and hexaploid samples across different numbers of sites for eleven different coverage amounts. The color of each point represents the best model. The shape of each point represents the approach used to simulate that sample. This represents a beta-binomial distribution with alpha free with a uniform mixture.

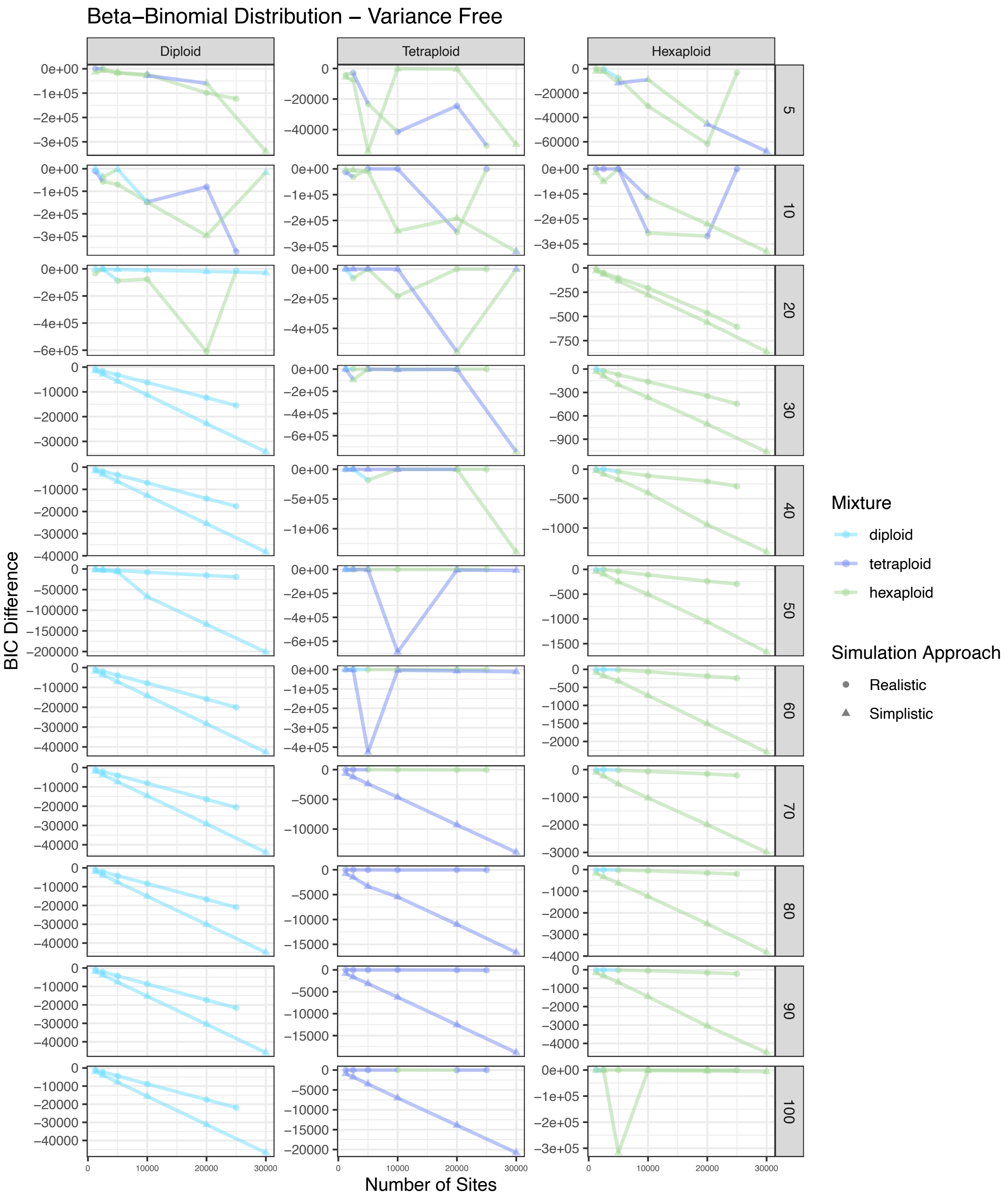


Figure S54. BIC score difference between the best and second best model for simulated diploid, tetraploid, and hexaploid samples across different numbers of sites for eleven different coverage amounts. The color of each point represents the best model. The shape of each point represents the approach used to simulate that sample. This represents a beta-binomial distribution with variance free.

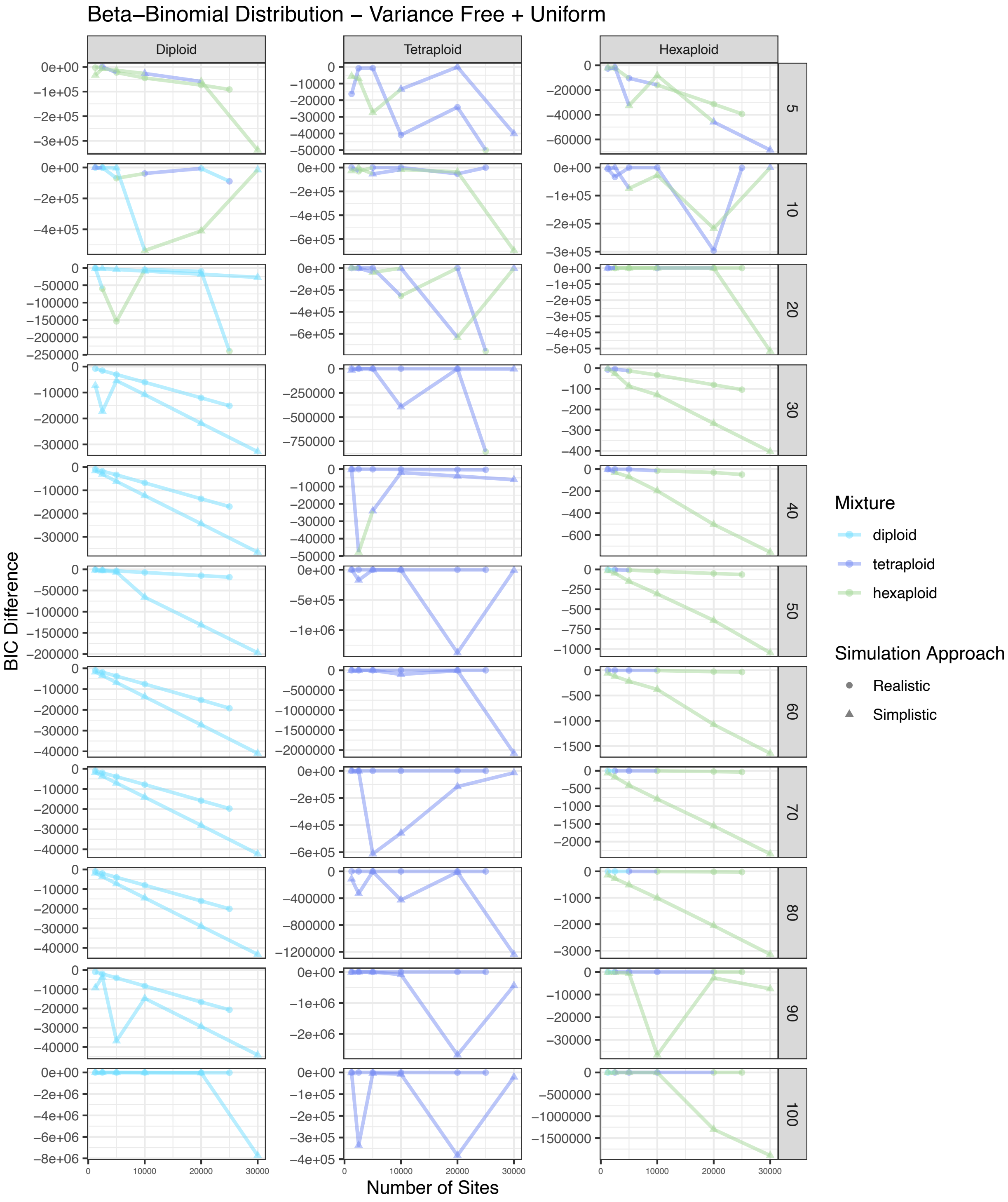


Figure S55. BIC score difference between the best and second best model for simulated diploid, tetraploid, and hexaploid samples across different numbers of sites for eleven different coverage amounts. The color of each point represents the best model. The shape of each point represents the approach used to simulate that sample. This represents a beta-binomial distribution with variance free with a uniform mixture.
