## Appendix S6 for "nQuack: An R package for predicting ploidal level from sequence data using site-based heterozygosity"

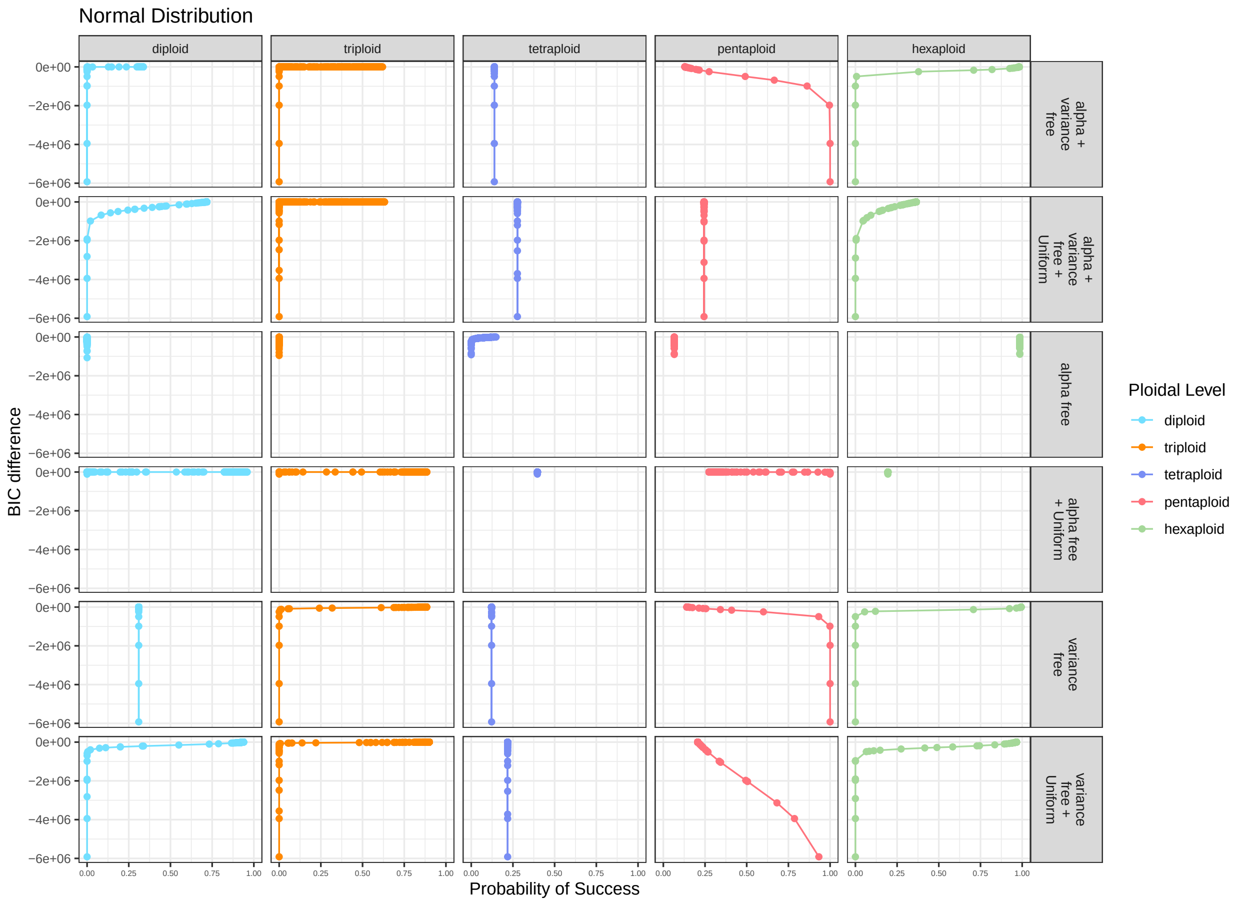


Figure S58. Probability of the correct model choice given the BIC difference between the best and second best model for all simulated diploid, triploid, tetraploid, pentaploid, and hexaploid samples. The probability of success was predicted based on a logisitic regression where accuracy is a function of BIC difference.

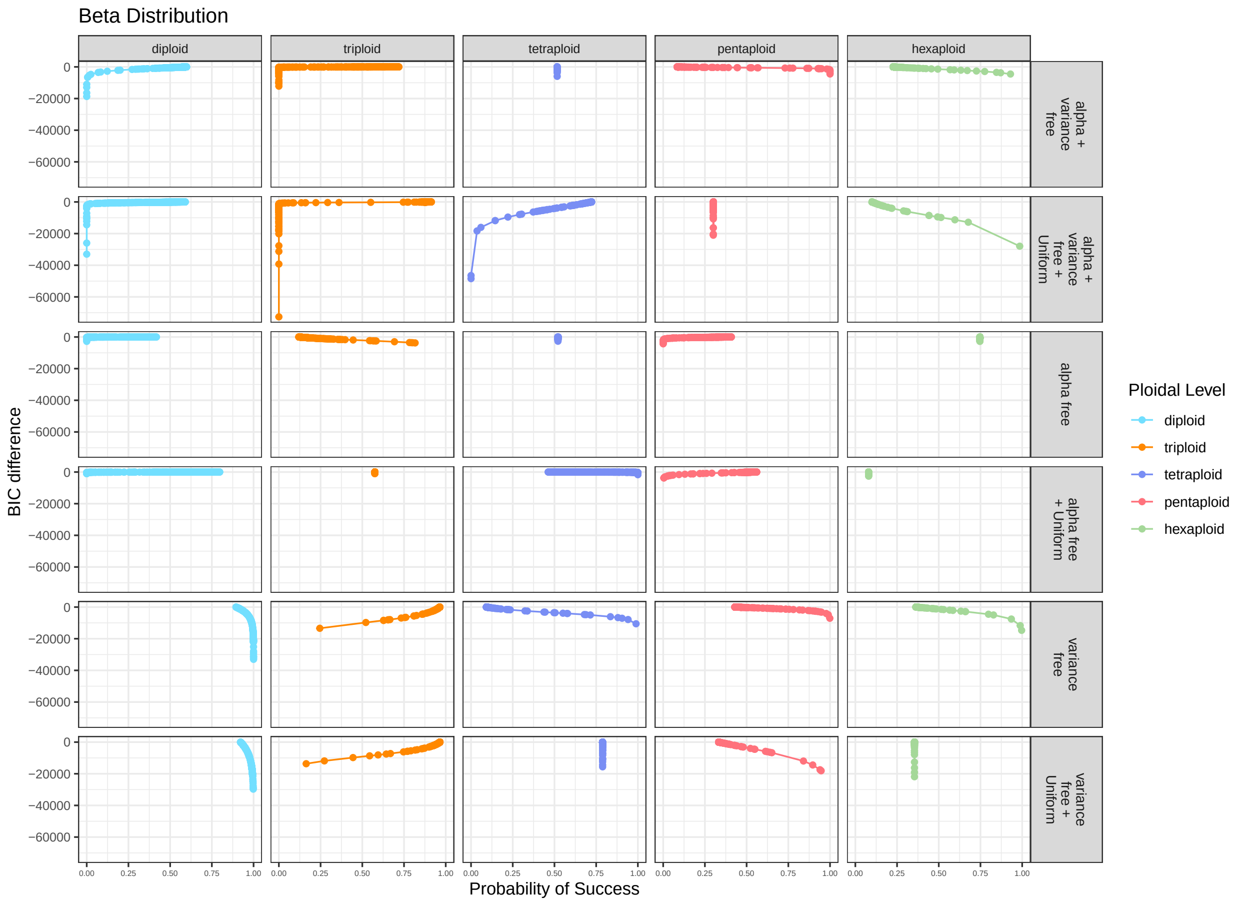


Figure S59. Probability of the correct model choice given the BIC difference between the best and second best model for all simulated diploid, triploid, tetraploid, pentaploid, and hexaploid samples. The probability of success was predicted based on a logisitic regression where accuracy is a function of BIC difference.

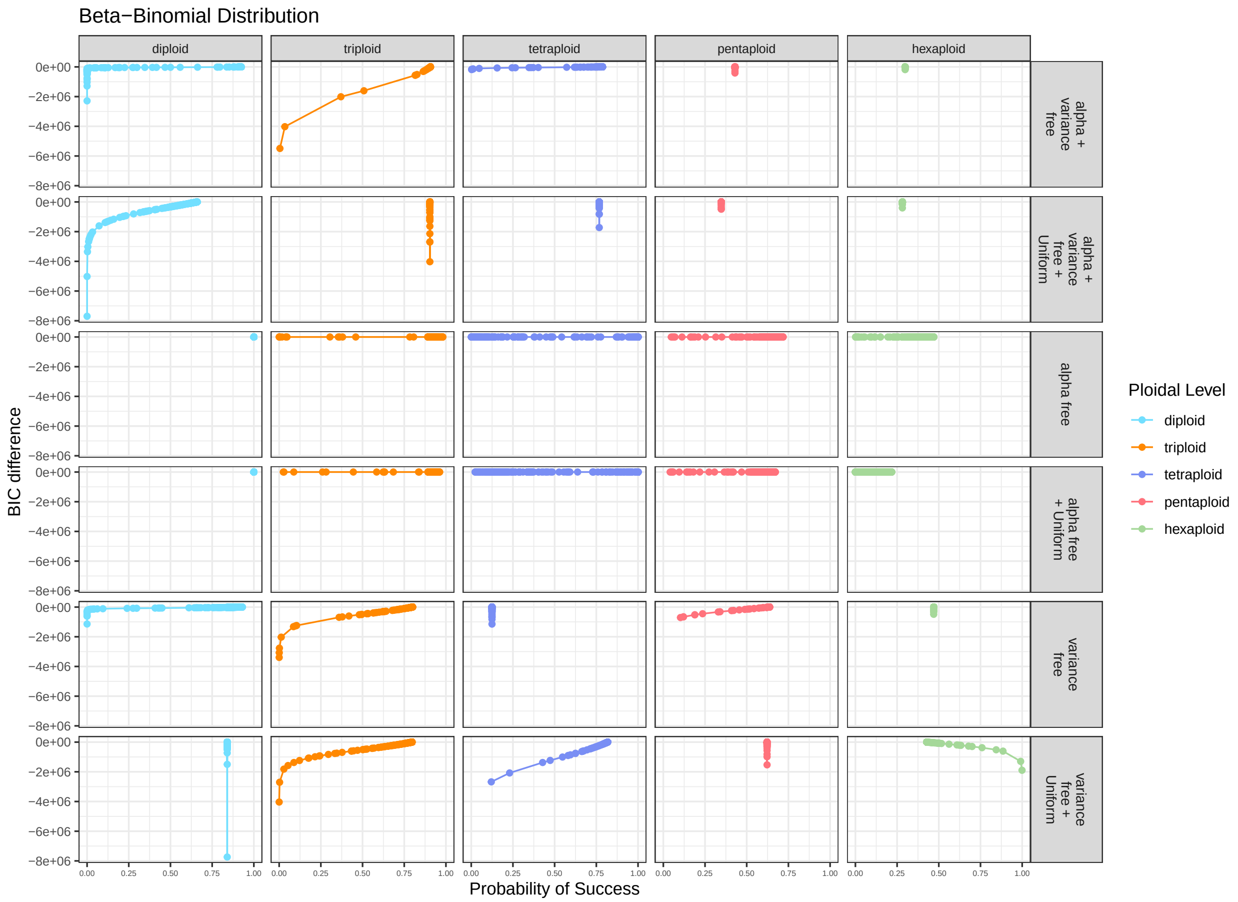


Figure S60. Probability of the correct model choice given the BIC difference between the best and second best model for all simulated diploid, triploid, tetraploid, pentaploid, and hexaploid samples. The probability of success was predicted based on a logisitic regression where accuracy is a function of BIC difference.
