## Appendix S7 for "nQuack: An R package for predicting ploidal level from sequence data using site-based heterozygosity"

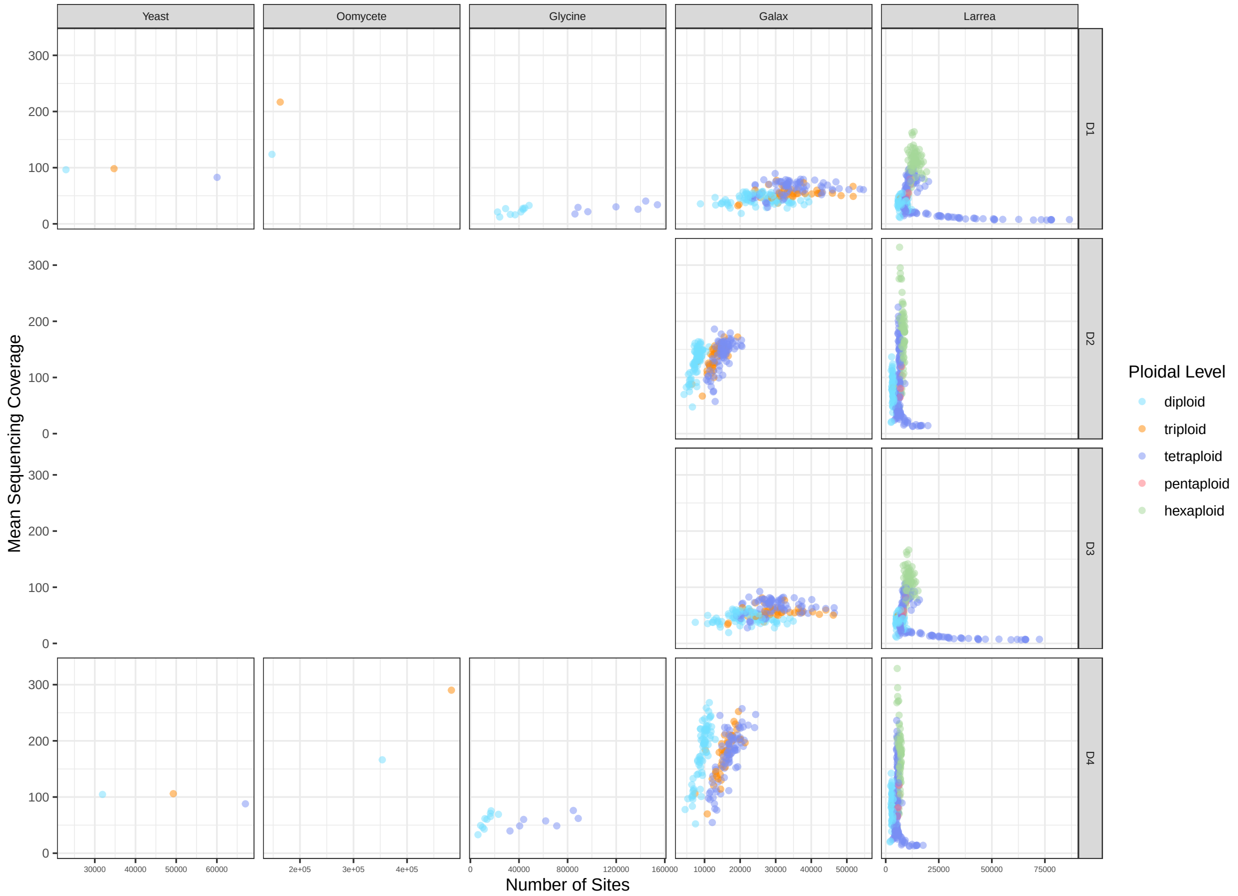


Figure S61. Number of sites and mean sequencing coverage for all data sets given the filtering type (see in-text for description).
